# Donor-matched iPSC model reveals context-dependent T2D genetic signals in fibro-adipogenic progenitors

**DOI:** 10.64898/2026.02.04.702388

**Authors:** Christa Ventresca, Arushi Varshney, Peter Orchard, Ha T.H. Vu, Yao-chang Tsan, Andre Monteiro da Rocha, Michael R. Erdos, Stephen I. Lentz, Leena Kinnunen, Timo A. Lakka, Jouko Saramies, Markku Laakso, Jaakko Tuomilehto, Karen L. Mohlke, Michael Boehnke, Laura J. Scott, Heikki A. Koistinen, Francis S. Collins, Todd Herron, Stephanie Bielas, Stephen C. J. Parker

## Abstract

Fibro-adipogenic progenitors (FAPs) in skeletal muscle have been implicated in type 2 diabetes (T2D), yet their heterogeneity and context-dependent regulation remain poorly understood. Here, we establish induced pluripotent stem cell (iPSC)-derived FAPs as a model of primary FAPs by leveraging a unique resource: iPSC lines and skeletal muscle biopsies obtained from 34 individuals, a subset of 30 profiled using single-nucleus multiomics. Donor-matched comparisons reveal iPSC-FAPs recapitulate the transcriptome, epigenome, and subtype composition of primary FAPs. Using single-nucleus multiomics, we show that high-insulin exposure drives iPSC-FAPs toward an adipogenic fate - and this adipogenic subtype is enriched for T2D GWAS signals, an enrichment undetectable at baseline. We map the T2D-associated rs3814707 non-coding signal to *LTBP3*, a gene that influences FAP adipogenic differentiation. These findings reveal how disease-relevant regulatory mechanisms can be masked in unstimulated cells and establish iPSC-FAPs as a powerful platform for dissecting the state-dependent biology of complex metabolic disease.

## Introduction

Type 2 diabetes (T2D) affects approximately 462 million individuals, representing 6% of the global population, and arises from a combination of genetic, environmental, and behavioral factors.^1–3^ Genome-wide association studies (GWAS) have identified over 1,200 independent signals associated with T2D, more than 90% of which localize to non-coding regions of the genome and individually have small effect sizes.^4,5^ Many of these variants map to regulatory elements that are active only in specific cell types,^4–12^ indicating that identifying the relevant cellular contexts is critical for interpreting GWAS signals and understanding disease mechanisms.

Skeletal muscle is a tissue where T2D-relevant regulatory activity has been identified; notably muscle regulatory elements are enriched for GWAS signals associated with fasting insulin levels.^13,14^ Within skeletal muscle, fibro-adipogenic progenitors (FAPs) have emerged as a cell population of particular interest. Genetic variants that regulate chromatin accessibility in FAPs colocalize with GWAS signals for T2D and related traits, suggesting that FAP-specific regulatory elements may harbor causal variants.^13,14^ FAPs are a heterogeneous population comprising multiple subtypes that reflect distinct differentiation trajectories: multipotent progenitors capable of following any trajectory, adipogenic cells poised to differentiate into adipocytes, and fibrogenic cells that promote fibrosis.^15–19^ The relative proportions of these subtypes shift in response to injury,^15,16^ suggesting that FAP composition is dynamically regulated by the cellular environment. However, the contribution of specific FAP subtypes in T2D pathophysiology has not yet been explored.

Beyond cell type identity, the cellular environment and cell state may also shape the regulatory landscape and thereby influence disease development. A previous study of immune cells demonstrated that exposure to different stimuli alters chromatin architecture, transcription factor (TF) binding, and gene expression.^20^ Notably, some regulatory elements exist in a “primed’ configuration that enables stimulus-specific transcriptional responses, indicating that chromatin structure can predetermine how cells respond to environmental cues. This principle suggests that studying FAPs under T2D-relevant conditions - rather than at baseline alone - may reveal disease-relevant regulatory mechanisms that would otherwise remain hidden.

To investigate how FAP subtypes and their regulatory pathways respond to a diabetogenic environment, we leveraged the Finland-United States Investigation of NIDDM Genetics (FUSION) Tissue Biopsy Study.^13^ This cohort includes skeletal muscle biopsies from 287 Finnish individuals with matched genotyping and single-nucleus multiomic profiling (snATAC-seq and snRNA-seq).^13,21^ Because FAPs constitute only a small fraction of cells in muscle biopsies, we used induced pluripotent stem cells (iPSCs) as an alternative platform. We differentiated iPSCs from 34 FUSION participants into FAPs and exposed a subset of 30 of them to a high-insulin environment, a metabolic milieu characteristic of T2D. We then performed single-nucleus multiomic profiling to characterize gene regulatory mechanisms across basal and high-insulin conditions. We found that a high-insulin state promotes the formation of the adipogenic FAP subtype and that chromatin accessibility peaks across FAP subtypes are differentially enriched for GWAS signals, establishing the relevance of this model system to T2D and related traits.

## Results

### FAP abundance in skeletal muscle biopsy and gene expression is associated with T2D status and FAP subtypes

We used skeletal muscle single-nucleus gene expression and chromatin accessibility profiles from 287 individuals in the FUSION Tissue study^13^ (Figure 1A) and identified 13 distinct cell types using the gene expression modality, of which 4.6% were annotated as FAPs (Figure 1B). We investigated whether FAP abundance was associated with donor traits including age, BMI, fasting plasma glucose, fasting serum insulin, sex, and T2D status. We modeled FAP counts across individuals while accounting for batch effects and donor traits using a negative binomial generalized linear model (NB-GLM) as implemented in DESeq2 (Figure 1C). Our analysis showed that increased FAP abundance was associated with higher fasting insulin, lower fasting plasma glucose levels, male sex, and T2D status (FDR<0.05).

**Figure 1:**
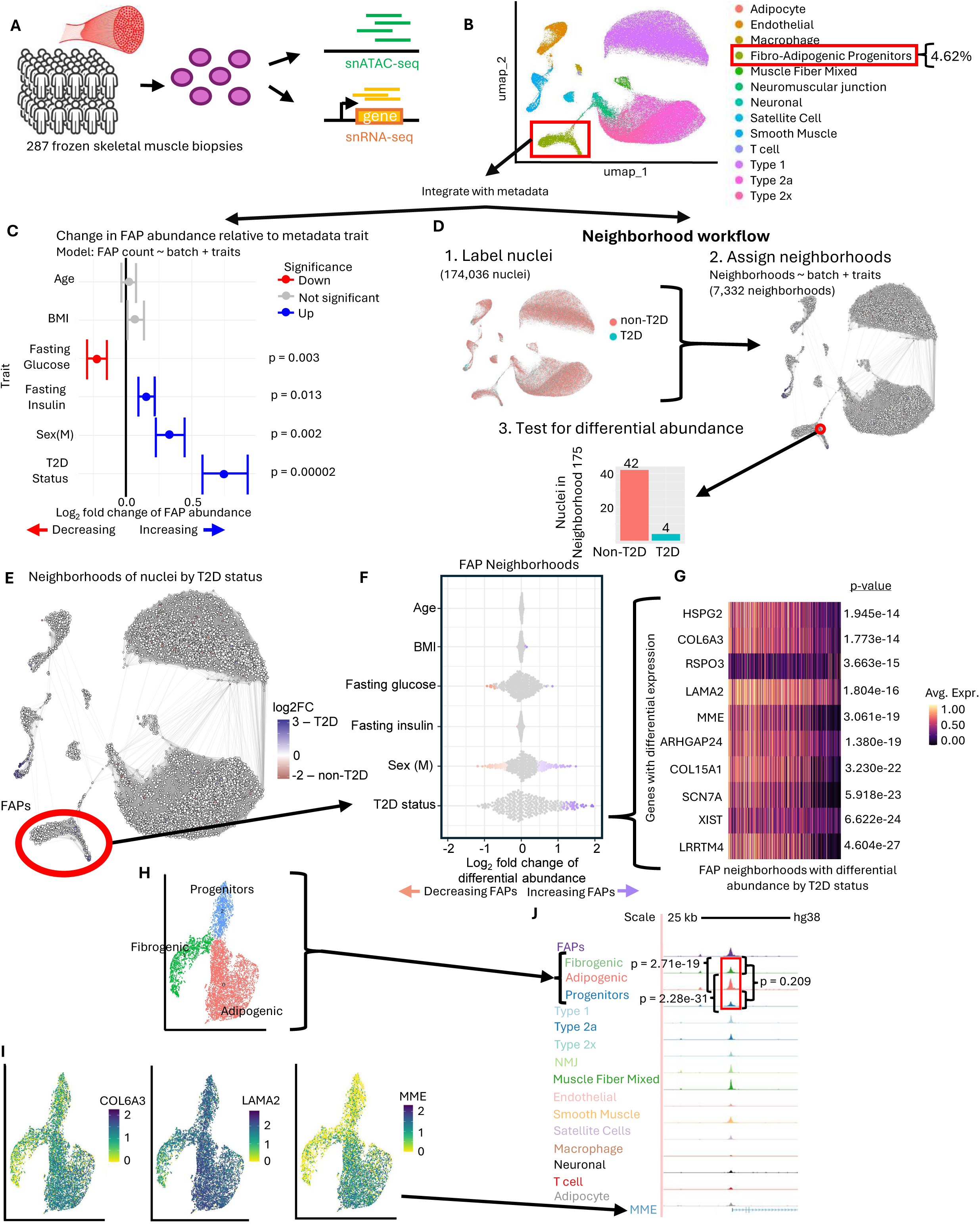
FAP abundance in skeletal muscle biopsy and gene expression is associated with T2D status and FAP subtypes. A – Schematic of the FUSION Tissue Study, a study of 287 Finnish individuals’ skeletal muscle biopsies. B – UMAP of cell types from the FUSION Tissue Study, fibro-adipogenic progenitors (FAPs) make up 4.6% of the nuclei based on gene expression. C – Comparison of FAP abundance with donor metadata, red indicates association with a decrease in FAP abundance and blue indicates an association with an increase in FAP abundance. DESeq2 was used for analysis and significance. Data represented as mean ± SEM. D – A diagram of the neighborhood workflow. E – UMAP of FUSION data indicating neighborhoods and colored by log2 fold change of differential abundance of T2D status. F – Beeswarm plots colored by log2 fold change of differential abundance for specifically the FAPs. G – Heatmap of differentially expressed genes within FAP neighborhoods based on differential abundance of T2D status. H – Clustering of FUSION FAPs revealed progenitors, fibrogenic, and adipogenic FAP subtypes. I – Multiple genes identified in G are revealed to have differential expression amongst the FAP subtypes. J – UCSC screenshot of FUSION skeletal muscle biopsy cell types, with the FAPs additionally separated by subtype. The promoter peak at *MME* has significantly different levels of chromatin accessibility amongst the FAP subtypes (DESeq2, normalized counts per 10 million reads).

We next asked whether analyzing all cell types within a skeletal muscle biopsy, agnostic of discrete cell-type cluster annotations, would recapitulate the observed FAP differential abundance. Using Milo^22^, we constructed a k-nearest neighborhood (knn) graph from the snRNA-seq data, assigned nuclei into 7,332 partially overlapping neighborhoods, and performed differential abundance testing across neighborhoods using a NB-GLM framework.^22^ We identified FAP-enriched neighborhoods with significantly (p<0.05) increased association with T2D status (80 neighborhoods), decreased fasting plasma glucose (49 neighborhoods), and association with sex (94 neighborhoods) (Figures 1E and F), consistent with our results shown in Figure 1C.

To further elucidate distinctions among FAP neighborhoods, we investigated which genes that were differentially expressed between neighborhoods with differential abundance and those without differential abundance. Specifically, we analyzed the top 2,000 most variable genes within the FAP neighborhoods, and identified the ten most significantly differentially expressed genes for each trait based on neighborhood-based differential abundance. In FAP neighborhoods with significantly higher T2D prevalence, we identified lower expression of *HSPG2, COL6A3, LAMA2, MME, ARHGAP24, COL15A1, SCN7A*, and *LRRTM4* and higher expression of *RSPO3* (FDR<0.05, Figure 1G, Figure S1). Among the ten most significantly differentially expressed genes, three - *HSPG2, COL6A3,* and *RSPO3* have been previously implicated in the development of T2D and diabetic neuropathy.^23–27^ Notably, *MME* is a regulator of cell growth and adipogenic differentiation.^28^ These observations reinforce the association between FAP abundance and T2D status and fasting plasma glucose, and highlight the potential contribution of FAP functional states, such as adipogenic differentiation, to disease-relevant phenotypes.

We next investigated whether *in vivo* FAPs comprised distinct functional subtypes. By clustering the FUSION FAP snRNA data, we identified three subtypes: fibrogenic, adipogenic, and progenitors, consistent with findings in previous studies^15–17,19^ (Figure 1H, Figure S1K). Notably, genes that were differentially expressed in FAP neighborhoods with high T2D-occurrence, including *LAMA2, COL6A3,* and *MME* (Figure 1G) also showed differential expression across FAP subtypes (Figure 1I). In addition to gene expression, we assessed the chromatin accessibility of the FAP subtypes and observed that *MME* was not only differentially expressed among FAP neighborhoods and FAP subtypes, but also exhibited significant differences in chromatin accessibility profiles between the FAP subtypes (FDR<0.05; DESeq2). Specifically, chromatin accessibility differed between adipogenic and fibrogenic subtypes (nominal p-value = 3e-19), adipogenic and progenitor subtypes (nominal p-value = 2e-31), whereas no significant difference was observed between fibrogenic and progenitor subtypes (nominal p-value = 0.2) (Figure 1J). Collectively, these findings suggest that FAP-specific gene regulatory mechanisms contribute to T2D etiology. Further investigation will enable us to pinpoint the pathways and FAP subtypes involved, offering deeper insight into their roles in the development of T2D.

### Induced pluripotent stem cell lines can be differentiated to fibro-adipogenic progenitors

For the remainder of this study different individuals/stem cell lines are utilized for different experiments. The selection of these lines was largely based on cell availability at the time, many of these experiments required large amounts of cells and therefore not every line was available to use for every experiment. Each line is indicated with an identifying number to highlight where experiments make use of the same or different lines. In total, we had access to 34 unique individual lines, however no one experiment was able to use all 34 individuals at once.

We developed a 21-day protocol to differentiate human iPSCs into FAPs by adapting the STEMdiff™ Mesenchymal Progenitor Kit (STEMCELL Technologies) with modifications based on established culture conditions for primary FAPs (Figure 2A).^29,30^ Phase one of the differentiation consists of preparing the iPSC lines (days -2 to 0). iPSCs maintained in mTeSR Plus (STEMCELL Technologies) were dissociated to single cells using Versene (STEMCELL Technologies) and plated at a density of 5 × 10L cells/cm² on Matrigel-coated plates in mTeSR Plus. During phase one, cells were cultured for 48 hours with a medium change at 24 hours, reaching approximately 30–50% confluency prior to induction. For phase two of the differentiation mesenchymal lineage was induced (days 0 to 4). On day 0, mTeSR medium was replaced with STEMdiff™-ACF Mesenchymal Induction Medium to initiate mesenchymal lineage commitment. Full medium changes were performed daily for four days. In this phase, cells transitioned from compact iPSC colony morphology toward a more dispersed, elongated appearance. The final phase consisted of FAP specification and maturation (days 4 to 21). On day 4, mesenchymal induction medium was replaced with FAP-specific medium (86% alpha-MEM, 13% FBS, 1% L-glutamine), formulated based on published protocols for primary FAP culture.^29^ On day 6, cells were passaged using the ACF Cell Dissociation Kit (STEMCELL Technologies) onto MatrixPlus plates (StemBioSys) to support FAP adhesion and expansion at a density of 5 × 10L cells/cm². Cells were maintained in FAP-specific medium with media changes every 24 hours until day 21, when they exhibited the characteristic elongated, spindle-shaped morphology of mature FAPs (Figure 2B-D). Throughout the differentiation we monitored cellular progression by live-cell imaging, capturing morphological changes throughout the 21-day protocol.

**Figure 2:**
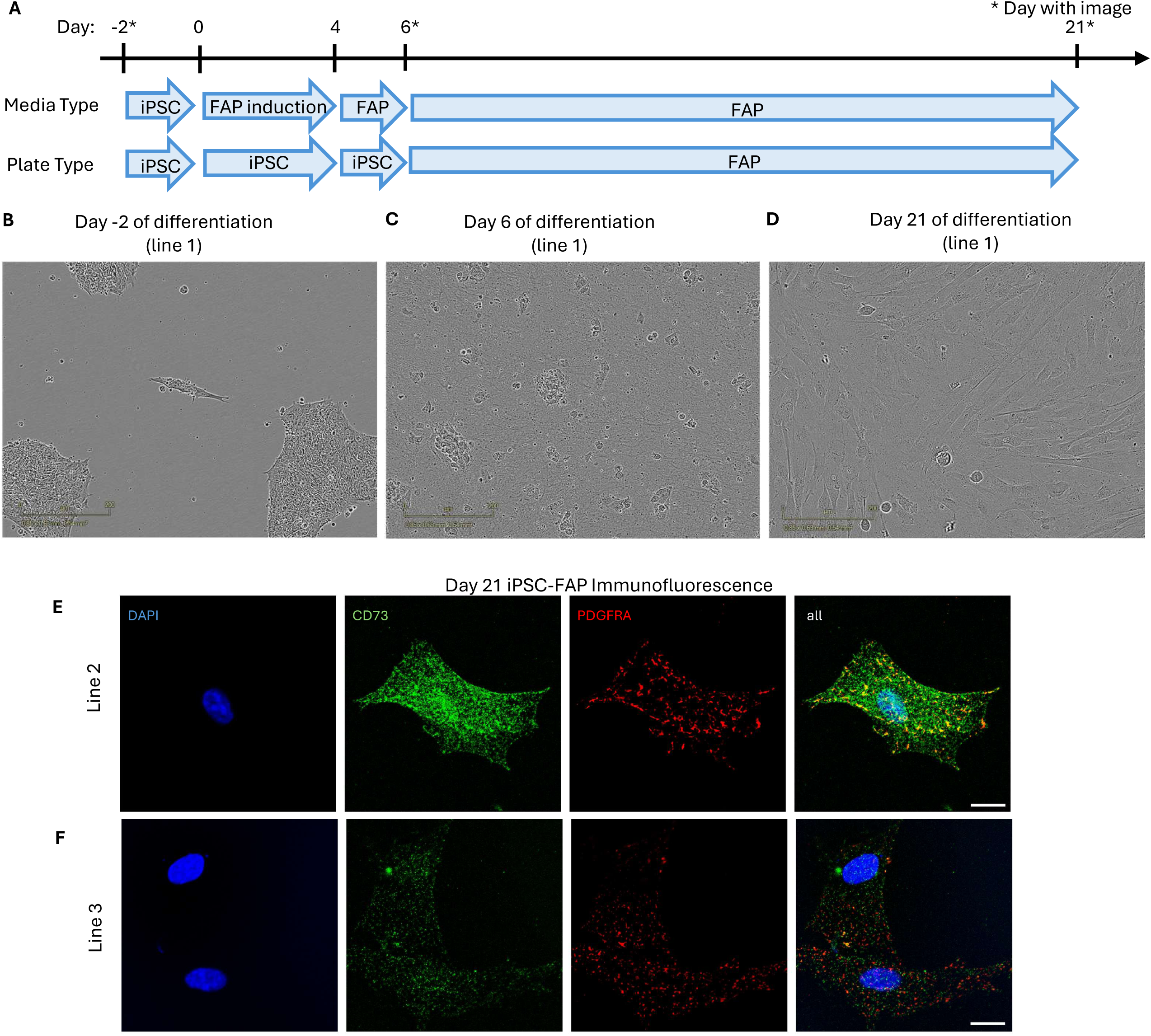
Induced pluripotent stem cell lines can be differentiated to fibro-adipogenic progenitors. A – Diagram of differentiation protocol to generate iPSC-FAPs. B – Image of day -2 iPSC-FAPs from a light microscope. Scale bar represents 200 um. C – Image of day 6 iPSC-FAPs from a light microscope. Scale bar represents 200 um. D – Image of day 21 iPSC-FAPs from a light microscope. Scale bar represents 200 um. E – Immunofluorescence of day 21 iPSC-FAP stained for two FAP markers: CD73 in green and PDGFRA in red from individual 1. Scale bar represents 20 um. F - Immunofluorescence of day 21 iPSC-FAP stained for two FAP markers: CD73 in green and PDGFRA in red from individual 2. Scale bar represents 20 um.

To further assess the efficiency of the FAP differentiation, we performed immunofluorescence staining on iPSC-FAPs from two individuals (female control and male T2D) using antibodies for two FAP markers (*CD73* and *PDGFRA*) at the day 21 timepoint. At the end of the differentiation cells showed robust staining for these markers, in contrast to undifferentiated iPSCs from the same individuals stained for the same markers which demonstrated no fluorescence (Figure 2E-F, Figure S2A-B). Together this demonstrates efficient and robust differentiation of iPSCs to FAPs using our protocol.

### iPSC-FAPs exhibit similar molecular features to *in vivo* FAPs

To further evaluate these differentiations we asked if the iPSC-FAPs recapitulate *in vivo* FAP molecular features. Successful differentiation was confirmed by flow cytometry at the terminal time point: iPSC-FAPs showed loss of the pluripotency marker *Tra-1-60* and acquisition of the FAP marker *NT5E* (*CD73*) as shown for four pilot differentiations using pools of ten lines (Figure 3A, Figure S3A). Representative examples from individual lines are shown in Figures S3B and C.

**Figure 3:**
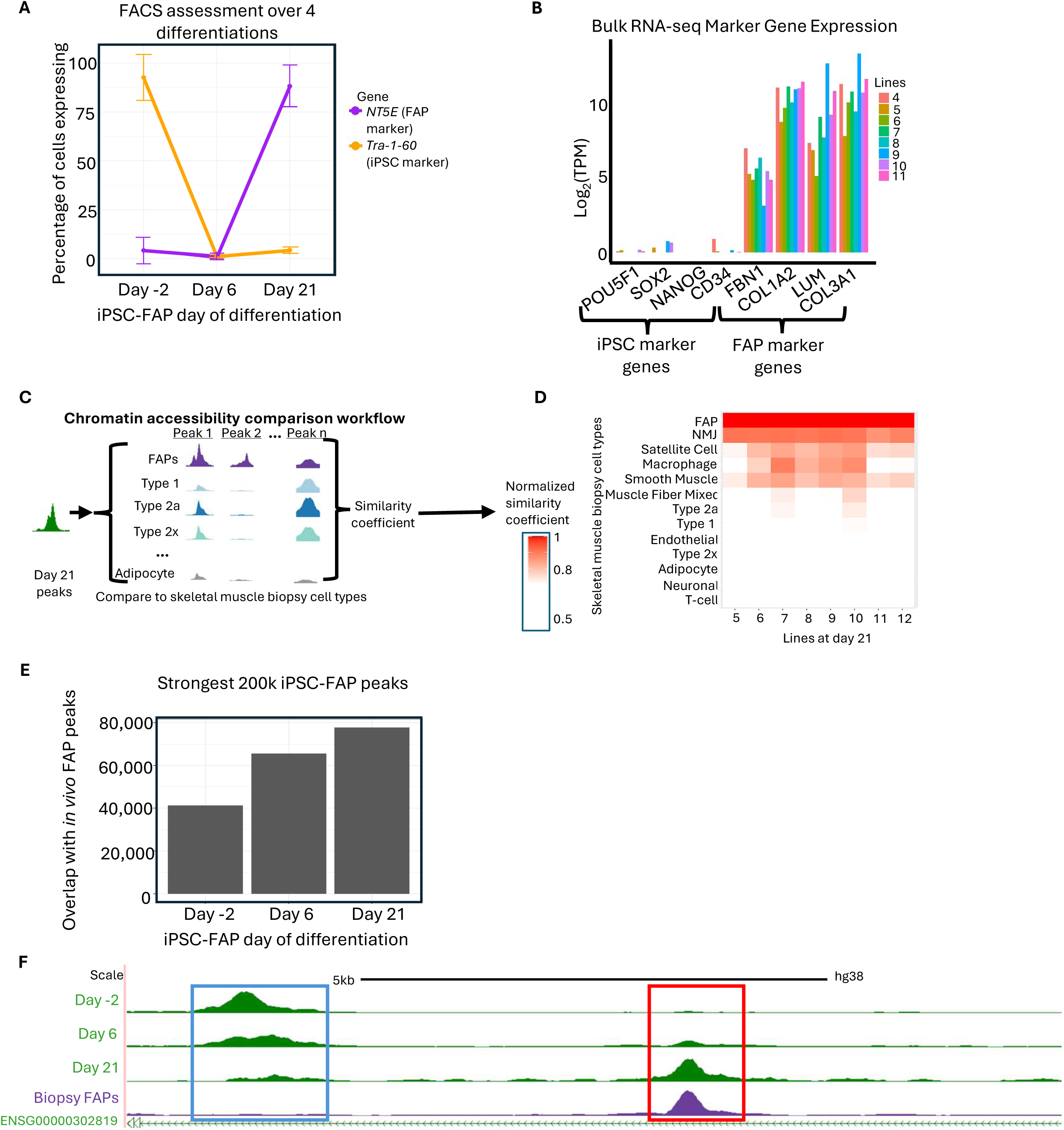
iPSC-FAPs exhibit similar molecular features to *in vivo* FAPs A – Flow cytometry examining an iPSC marker gene and a FAP marker gene over the course of four different differentiations of a pooled sample containing ten different individuals. Data represented as mean ± SEM. B – Bar plot showing expression of four FAP marker genes and four iPSC marker genes for eight individuals in bulk RNA-sequencing samples. C – Schematic illustrating the analysis performed in 3D. D – Logistic regression of chromatin accessibility peaks for eight individuals at the endpoint for FAP differentiation (day 21) to compare the differentiated cells to cell-type specific chromatin accessibility peaks from FUSION skeletal muscle biopsy cell types. Two individuals dropped out of the day 21 samples and are excluded from this analysis. E – Bar plot of the overlapping peaks between the iPSC-FAP chromatin accessibility profiles and the *in vivo* FAP chromatin accessibility profiles. F – UCSC browser screen shot of chromatin accessibility data from the beginning (day -2), middle (day 6), and end (day 21) of a pooled sample of ten individuals across the FAP differentiation. At the bottom is the chromatin accessibility data from the FUSION FAPs. Two peaks of interest are highlighted with the blue and red boxes.

We next assessed whether iPSC-FAPs expressed canonical FAP marker genes. Specifically, we generated bulk RNA-seq data and quantified the expression of four iPSC marker genes (*POU5F1, SOX2, NANOG,* and *CD34)* as well as four FAP marker genes (*FBN1, COL1A2, LUM,* and *COL3A1)*. Across all individuals in the preliminary group, there was robust expression of the FAP marker genes and very little expression of the iPSC marker genes (Figure 3B). We further explored these differentiations by examining the chromatin accessibility in iPSC-derived FAPs and assessing their similarity to *in vivo* FAPs. To this end, we generated single-nucleus chromatin accessibility (snATAC) data for a pooled sample of 10 lines at three timepoints during differentiation: early (day -2), intermediate (day 6), and late (day 21). To compare the chromatin accessibility profiles of the differentiated iPSC-FAPs with those of *in vivo* skeletal muscle cell types, we performed a logistic regression using ATAC peaks from the terminal time point and corresponding peaks from the cell types profiled in FUSION skeletal muscle biopsies (Figure 3C).^14^ We observed that the iPSC-FAPs were the most similar to *in vivo* FAPs compared with other skeletal muscle cell types (Figure 3D).

We next evaluated which differentiation stage of iPSC-FAPs most closely resembled biopsy-derived FAPs by computing Jaccard similarity scores using the top 200,000 most significant peaks for each iPSC-FAP timepoint, ranked by p-value. As expected, peak overlap increased progressively with differentiation stage, the early time point (day -2) showed the fewest overlapping peaks (41,230), increasing to 77,704 peaks at the terminal time point (day 21, Figure 3E). This corresponds to the emergence of at least 36,474 new, shared accessible regions over the course of the differentiation. For example, we observed a chromatin accessibility peak (blue box; chr4:12,884,865-12,885,328) present at early stages that gradually diminished as differentiation advanced (Figure 3F). Conversely, a nearby peak (red box; chr4:12,889,003-12,889,441) emerged during differentiation and ultimately recapitulated the accessibility profile seen in biopsy-derived FAPs (Figure 3F); examples of individual lines are shown in Figure S3D.

### iPSC-FAPs respond to insulin stimulation

To confirm the identities of the 30 iPSC lines used for iPSC-FAP stimulatory state specific data generation out of the total of 34 individuals, we performed low-pass whole-genome sequencing, which verified that all lines matched their respective donor reference data with high confidence (Figure 4A). We then differentiated all 30 lines into iPSC-FAPs and evaluated the proportion of cells expressing FAP marker genes at the terminal time point (day 21). An average of 74% of cells expressed the FAP marker gene *NT5E* based on FACS analysis, confirming that the majority of lines successfully differentiated to FAPs (Figure 4B).

**Figure 4:**
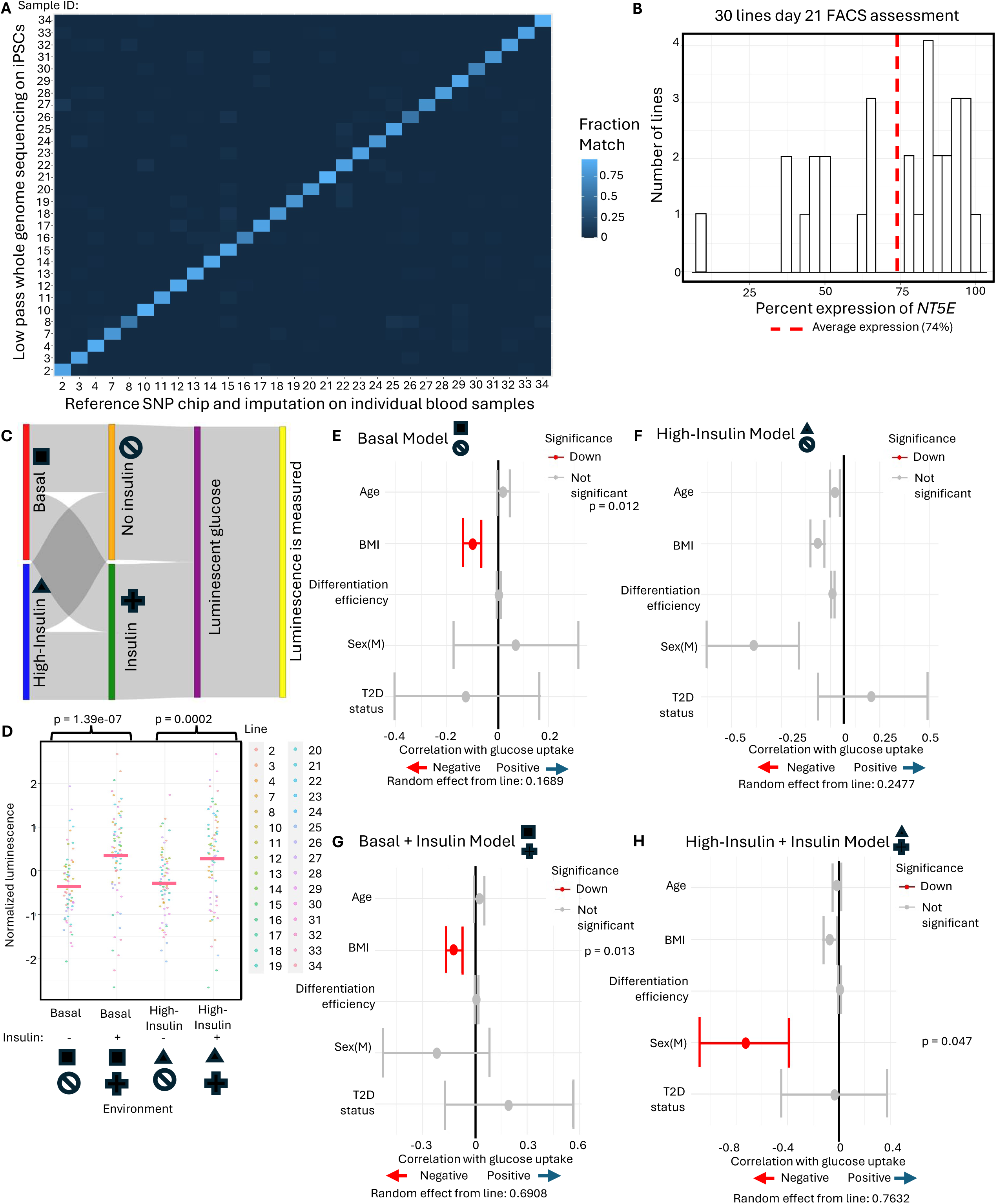
iPSC-FAPs respond to insulin stimulation. A – A heatmap comparing reference genotyping data from the FUSION study to low-pass genotyping data generated as part of QC metrics for this study. The fraction match indicates the fraction of reads that aligned from the low pass WGS data to the reference SNP Chip. B – Histogram of FAP marker gene expression based on FACS assessment of 30 different lines. The X-axis indicates the percentage of cells per line expressing *NT5E*. The average across all cell lines was 74% of cells expressing the FAP marker gene. C – Sankey plot representing the insulin-stimulated glucose uptake assay. D – Comparison of normalized luminescence from all 30 donors (with replicates) in basal, basal with a small dose of insulin, high-insulin, and high-insulin with a small additional dose of insulin. Pink lines indicate the average luminescence value for that environmental condition. A paired Student’s t-test was used to assess statistical significance. E – An interval plot comparing donor metadata to luminescence levels in basal conditions. Red indicates a negative correlation with glucose uptake. Data represented as mean ± SEM. F – An interval plot comparing donor metadata to luminescence levels in high-insulin conditions. Red indicates a negative correlation with glucose uptake. Data represented as mean ± SEM. G – An interval plot comparing donor metadata to luminescence levels in basal conditions with an additional small dose of insulin. Data represented as mean ± SEM. H – An interval plot comparing donor metadata to luminescence levels in high-insulin conditions with an additional small dose of insulin. Red indicates a negative correlation with glucose uptake. Data represented as mean ± SEM.

To assess insulin responsiveness, we subjected all 30 iPSC-derived lines to an insulin-stimulated glucose uptake assay (Figure 4C, Figure S4A). In response to the dose of insulin, we observed a significant increase in luminescence indicating enhanced glucose uptake (Figure 4D, individual example in Figure S4B). Next, we examined how donor-specific metadata traits such as age, BMI, FACS-based differentiation efficiency (Figure 4B), sex, and T2D status correlated with cellular glucose uptake using a linear mixed model. We found that BMI is inversely correlated with glucose uptake under basal conditions (p<0.05, Figure 4E and G) consistent with previous studies linking higher BMI to impaired glucose uptake.^31,32^ Under high-insulin conditions, male sex emerged as a significant predictor of decreased glucose uptake, but only following insulin stimulation (p<0.05, Figure 4F and H). Previous studies have suggested that men are more likely to develop insulin resistance than women, and subsequent T2D.^33^ These results highlight how changes in stimulatory context between basal and high-insulin environments modulate cellular responses to the same stimulus. Overall, our present experiment provides line-specific physiological response data, confirms differential responses to environmental perturbations, and establishes a foundation for identifying associated gene regulatory mechanisms underlying insulin responsiveness.

### The cell village of 30 lines demonstrates responsiveness to high-insulin state

Maintaining all 30 iPSC-FAP lines in separate plates can result in plate-specific effects that may confound downstream analyses; therefore, we pooled cells from all 30 individuals into a single “cell village” to minimize this source of technical variation.^34^ The cell village included 12 males and 18 females, including 12 individuals with T2D and 18 non-T2D controls (Figure 5A). We then divided the cell village into two pools, one maintained under basal conditions and the other exposed to a high-insulin environment. To assess the representation of each donor within the cell village prior to omics profiling, we performed low-pass whole genome sequencing (WGS) and applied census-seq to quantify donor representation (Figure S5A). Donor representation across basal and high-insulin environments did not differ with at least 25 individuals each contributing ≥1% of nuclei per sample, enabling robust comparisons between conditions.

**Figure 5:**
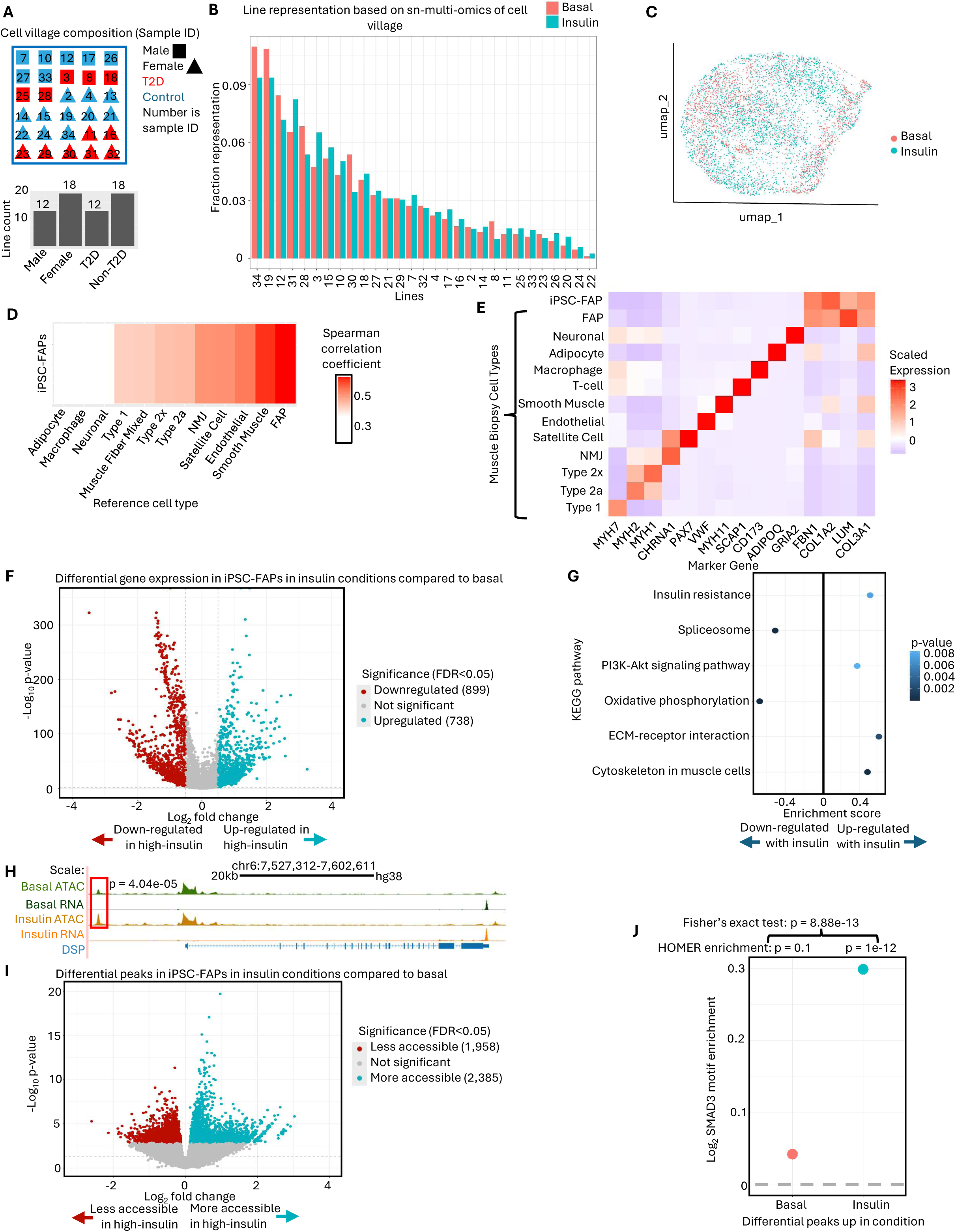
The cell village of 30 donors demonstrates responsiveness to a high-insulin state. A – Schematic of the 30 donors included in the cell village. B – Bar plot of line representation in basal and high-insulin states as indicated by color and assessed by sn-multi-omics data. C – Clustering of the snRNA data, labelled by environment. D – Spearman correlation comparing *in vivo* gene expression TPM values of FAPs to the iPSC-FAP cell village. E – Heatmap of marker gene expression levels for skeletal muscle biopsy cell types and iPSC-FAPs (top row) with four FAP marker genes shown (right columns). F – Volcano plot of gene expression comparing insulin conditions to basal as assessed by MAST. G – Gene set enrichment pathways using KEGG analysis within the differential gene expression data comparing the high-insulin state to the basal state. H – UCSC browser screenshot of an extended view of a differential peak (left side). This peak is near the gene *DSP* (ENSG00000096696), which is known to be involved in the cytoskeleton in muscle cells and is included in Table S1 and in the pathways in Fig 5G (normalized counts per 10 million reads). I – A volcano plot comparing differential peaks between high-insulin and basal states. J – Plot of log2 SMAD3 motif enrichment (percent of reads with motif in target sequences divided by percent of reads with motif in background sequences) in differential peaks colored by subset of peaks (basal or insulin). Significance determined by both HOMER analysis of motif enrichment and Fisher’s exact test of counts.

Next, we performed single-nucleus gene expression and chromatin accessibility profiling using the 10x Genomics Epi Multiome ATAC + Gene Expression platform to investigate transcriptional and regulatory features of the iPSC-FAPs across different environmental conditions. We used demuxlet to assess donor representation in the single nucleus data and observed similar representation levels (Figure 5B), consistent with previous estimates from WGS-based donor quantification (Figure S5B). After quality control, we jointly clustered 1,990 high-quality basal nuclei and 3,038 high-insulin nuclei, which integrated homogeneously across conditions (Figure 5C). To assess transcriptional similarity between iPSC-FAPs and *in vivo* FAPs, we computed Spearman correlations using normalized TPM values. Across all individuals in the cell village, iPSC-FAPs showed the highest correlation with *in vivo* FAPs, indicating strong concordance in gene expression profiles (Figure 5D and S5C). We further compared expression levels of marker genes for FAPs and other skeletal muscle biopsy cell types, and observed that the *in vivo* FAPs and *in vitro* iPSC-FAPs exhibited similar marker expression trends, reinforcing their similarities in the gene expression modality (Figure 5E). This indicated that the cell village recapitulated expected FAP molecular features.

To explore differences in gene expression between the basal and high-insulin environments, we performed differential gene expression analysis, identifying 738 up-regulated genes (higher expression in high-insulin compared with basal conditions) and 899 down-regulated genes (Figure 5F). Up-regulated genes were enriched for pathways including “insulin resistance,” “PI3K-Akt signaling pathway,” and “ECM-receptor interaction” (FDR<0.05), consistent with previous findings characterizing cellular responses to insulin.^35–37^ We also observed enrichments of the “cytoskeleton in muscle cells” pathway among up-regulated genes, suggesting continued differentiation under high-insulin conditions. Conversely, downregulated genes were enriched for “spliceosome” and “oxidative phosphorylation” pathways (Figure 5G, full results in Table S1). Reduced oxidative phosphorylation is a well-established feature of high-insulin states,^38^ and prior work has shown that spliceosome components are misregulated - and specifically downregulated - in T2D, particularly in pancreatic beta cells.^39^ Additional GO term enrichment analysis, presented in the supplemental data, were consistent with KEGG pathway results (Figure S5D).

Next, we used a Negative Binomial Generalized Linear Mixed Model (NB-GLMM) to identify differentially accessible chromatin between basal and high-insulin environments (Figure 5H). We identified 1,958 (0.9%) peaks with greater accessibility in the basal state, and 2,385 (1%) peaks with greater accessibility under high-insulin conditions (FDR<0.05, Figure 5I). Notably, a regulatory element near *DSP*, a gene involved in the cytoskeleton in muscle cells KEGG pathway (Table S1), showed increased accessibility in the high-insulin condition (Figure 5H). To identify transcription factors that may mediate insulin-responsive chromatin remodeling, we tested whether differentially accessible chromatin peaks were enriched for known transcription factor binding motifs. We identified 139 motifs significantly enriched in peaks more accessible under high-insulin conditions and 151 peaks more accessible under basal conditions. (FDR<0.05, Table S2 and S3). Peaks with more accessible chromatin under high insulin were strongly enriched for the SMAD3 binding motif (Figure 5J, Table S2, HOMER p = 1e-12; Fisher’s exact test comparing basal vs. insulin enrichment, p = 9e-13). Consistent with this finding, *SMAD3* expression was also upregulated under high-insulin conditions (Figure 5F).

Given the established role of *SMAD3* in adipogenesis,^40^ these results suggest that SMAD3 may link insulin exposure to the adipogenic fate in iPSC-FAPs. Taken together, these analyses established a high-quality multi-omics dataset for dissecting how iPSC-FAPs respond to high-insulin through coordinated gene expression and chromatin accessibility.

### The high-insulin state induces the FAPs to cultivate the adipogenic subtype

We also explored the FAP subtypes in our population of iPSC-FAPs. Consistent with the FUSION FAPs, we identified three distinct sub-populations characterized by the expression of *E2F1*, *SEMA3C*, and *THY1* markers (Figure 6A, Figure S6A–C). Additionally, we performed pseudotime analysis to further refine subtype identification by leveraging comparisons between nuclei in the gene expression modality to assemble trajectories of development. This analysis revealed developmental trajectories corresponding to FAP progenitors, adipogenic cells, and a fibrogenic cluster (Figure 6B). Interestingly, when pseudotime analysis was conducted separately on basal and high-insulin samples, we observed that the basal condition comprised only progenitor and fibrogenic subtypes, while the high-insulin condition encompassed progenitor, fibrogenic, and adipogenic subtypes (Figure S6D and E).

**Figure 6:**
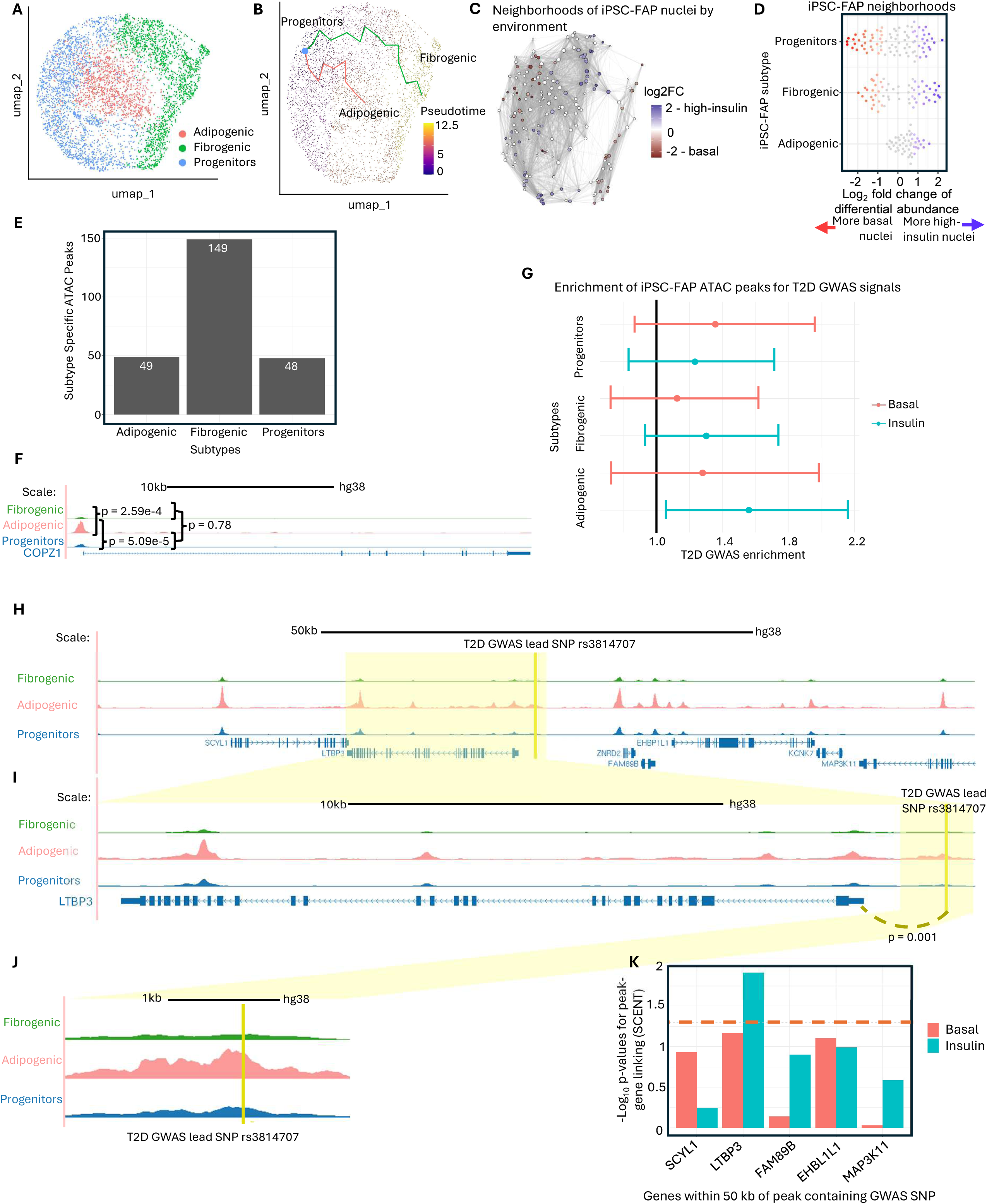
The high-insulin state induces the FAPs to cultivate the adipogenic subtype. A – Clustering of the snRNA data, labelled by subtype. B – Pseudotime trajectory within the clustering, colors indicate FAP subtypes. The FAP subtypes of progenitors, adipogenic, and fibrogenic are labelled. C – UMAP of iPSC-FAP data indicating milo neighborhoods and colored by log fold change of differential abundance of environment (basal or insulin). D – Beeswarm plot from milo colored by log2 fold change of differential abundance for environment. P-values comparing peaks by subtype from GLMNB analysis comparing subtypes to peak sizes. E – Bar plot of chromatin accessibility peaks specific to iPSC-FAP subtypes. F – UCSC screenshot of gene *COPZ1* and FAP subtype chromatin accessibility peaks. Chromatin accessibility peaks are significantly different amongst the subtypes (FDR < 0.05). G – Plot of enrichment by FAP subtype chromatin accessibility peaks, looking at Suzuki *et al.*, Nature, 2024 loci. Color indicates sample (basal or high-insulin). H – UCSC screenshot showing the *LTBP3* locus along with 50 kb to either side. The dark yellow line indicates the location of rs3814707. I – UCSC screenshot showing the *LTBP3* locus and its location on chromatin accessibility peaks of the iPSC-FAP subtypes. Dark yellow line indicates location of rs3814707, dark yellow dashed line shows proposed chromatin looping with p-value from SCENT analysis. J – UCSC screenshot showing the *LTBP3* locus and its location on chromatin accessibility peaks of the iPSC-FAP subtypes. The dark yellow line indicates the location of rs3814707. K – Bar plot showing p-values of correlation between chromatin accessibility peak shown in 6H-J and expression of genes within 50 kb based on SCENT analysis (two genes were removed from analysis due to low expression in iPSC-FAPs) in the basal sample and high-insulin sample. The orange dashed line indicates p-value = 0.05.

Nuclei from the basal and high-insulin samples integrated well; however, the relative contributions of nuclei from each condition differed across subtypes. We therefore hypothesize that the high-insulin sample would contain more nuclei within the adipogenic FAP subtype. Indeed, the neighborhood analyses on the joint clustering of basal and insulin samples revealed that adipogenic neighborhoods were enriched for high-insulin nuclei (Figure 6C and D). Furthermore it showed that the adipogenic subtype contained significantly more high-insulin than basal nuclei (binomial test, p-value = 4e-10, Figure S6F). These findings are consistent with previous studies showing that insulin serves as a key trigger for the adipogenic pathway in FAPs.^41,42^ Additionally, the iPSC-FAP subtypes exhibit distinct chromatin accessibility landscapes. When examined using an NB-GLMM, we observed 246 subtype-specific peaks - namely 49 adipogenic, 149 fibrogenic, and 48 progenitor specific peaks (Figure 6E). For example, the *COPZ1* promoter shows significantly higher accessibility in the adipogenic subtype (FDR<0.05; Figure 6F). Specifically, chromatin accessibility differed between adipogenic and fibrogenic subtypes (NB-GLMM, nominal p-value = 2.59e-4) and adipogenic and progenitor subtypes (nominal p-value = 5.09e-5) while the fibrogenic and progenitors did not significantly differ (nominal p-value = 0.78, Figure 6F).*COPZ1* is involved in lipid transport and cellular growth, making it relevant to adipogenic differentiation and this subtype.^43,44^ Collectively these findings highlight how changes in the cellular environment can alter FAP differentiation trajectories, with the high-insulin conditions favoring adipogenic differentiation.

To assess the overlap of iPSC-FAP subtype-specific chromatin accessibility peaks with existing T2D GWAS data, we performed an enrichment analysis which revealed that only the adipogenic subtype is significantly enriched for T2D GWAS signals (Figure 6G).^4,45^ These findings highlight the particular relevance of the adipogenic subtype to T2D, and underscore the importance of studying cells under varied environmental conditions. Notably, without the addition of insulin to the cellular environment, our samples would contain significantly fewer nuclei belonging to the adipogenic cluster (p-value = 4e-10, Figure S6F), limiting our ability to detect this enrichment. Consistent with this observation, when the basal and high-insulin FAP subtypes were tested separately for enrichment only high-insulin adipogenic FAPs showed significant enrichment for T2D GWAS signals (Figure 6G).

Within this subset of peaks overlapping a T2D GWAS loci, the lead SNP rs3814707 signal emerged as a notable example (GWAS p-value = 3e-09^4^). This signal is located within 50 kb of seven genes (Figure 6H). To determine possible gene interactions, we leveraged co-accessibility analyses from *in vivo* FAPs, which indicated that this peak is connected to the nearby gene LTBP3.^13^ Additionally, using SCENT we assessed correlations between chromatin accessibility at this peak and expression of genes within 50 kb. Among these genes, only *LTBP3* showed a significant association with accessibility at this locus, with proposed chromatin looping shown (Figure 6I). Notably, this GWAS signal resides at the summit of an adipogenic subtype accessibility peak (Figure 6J). Importantly this association was only significant when testing high-insulin nuclei (Figure 6K), under basal conditions no genes within 50 kb are significantly associated with this peak (Figure 6K). *LTBP3* is known to regulate the extracellular matrix and cellular growth, and previously has been implicated in the regulation of adipogenic differentiation in FAPs.^46,47^

## Discussion

This study establishes two main findings. First, we demonstrate that iPSC-derived FAPs faithfully recapitulate the molecular and cellular properties of *in vivo* FAPs isolated from skeletal muscle. This validation is uniquely rigorous, because our iPSC lines and muscle biopsies were derived from the same individuals, enabling direct donor-matched comparisons that have not been done in previous studies. Second, we show that the adipogenic subtype of FAPs is enriched for T2D GWAS signals, and that this enrichment emerges only when cells are exposed to a high-insulin environment. Together, these findings establish iPSC-FAPs as a robust model system for studying FAP biology and demonstrate how disease-relevant regulatory mechanisms can be uncovered through context-specific profiling.

The donor-matched design of our data allowed us to benchmark iPSC-FAPs against *in vivo* FAPs with unprecedented precision. iPSC-FAPs closely resemble their *in vivo* counterparts in morphology, marker gene expression, and chromatin accessibility profiles. Critically, the same major FAP subtypes - progenitors, adipogenic, and fibrogenic - are identifiable in both systems using the same marker genes. This molecular and cellular concordance demonstrates that iPSC differentiation captures the essential features of FAP biology, rather than merely producing a superficial resemblance. Our data resolve this uncertainty and provide a foundation for using iPSC-FAPs to investigate aspects of FAP biology that are impractical to study using primary tissue, including responses to controlled environmental perturbations.

Beyond establishing a robust model system, this study reveals that cellular context is critical for identifying disease-relevant regulatory mechanisms. The adipogenic FAP subtype showed significant enrichment for T2D GWAS signals only under high-insulin conditions, while this enrichment was not detectable at baseline. iPSC-FAPs are functionally responsive to insulin, as evidenced by increased glucose uptake and a shift in subtype composition toward the adipogenic fate. This state dependence is consistent with evidence from other systems that regulatory elements can exist in “primed” configurations, enabling context-specific transcriptional responses.^20^ Our findings reinforce a broader principle: GWAS interpretation efforts focused exclusively on baseline cellular states may overlook regulatory mechanisms active only under disease-relevant conditions.

FAP abundance in skeletal muscle is associated with T2D status, fasting glucose, and fasting insulin, consistent with a role for this cell population in disease etiology. Our results support a model in which T2D risk variants act through FAPs to promote adipogenic differentiation in response to elevated circulating insulin levels, demonstrated by the specific enrichment of the adipogenic iPSC-FAP subtype under high-insulin conditions. One locus of particular interest is rs3814707, where a T2D GWAS signal lead SNP resides within a chromatin accessibility peak in the adipogenic subtype and chromatin looping analyses indicate this peak interacts with the gene *LTBP3*. Although LTBP3 has not been directly linked to T2D prior to this study, we propose that such variants may bias FAP differentiation toward adipogenesis under high-insulin conditions, thereby contributing to increased intramuscular adipose tissue in individuals with T2D. This model generates testable hypotheses regarding how non-coding variants influence T2D pathophysiology through effects on progenitor cell fate decisions.

Several questions remain for future investigation. Future studies should investigate the downstream transcriptional consequences of GWAS-implicated regulatory variants in FAPs to elucidate the pathways through which they contribute to the disease development. It will also be of interest to explore additional stimulatory conditions to determine whether other FAP subtypes exhibit GWAS enrichment in distinct environmental conditions. The iPSC-FAP platform we established here provides a tractable system for addressing these questions through systematic profiling across diverse environmental contexts.

## Limitations

Our study has some limitations. Although iPSC-derived FAPs closely resemble *in vivo* FAPs, they are not identical to primary cells. Our donor-matched design mitigates this limitation by enabling direct comparison to skeletal muscle biopsies from the same individuals; however, iPSC-FAP findings should ultimately be validated in primary tissue where feasible. Additionally, all participants in our study are of Finnish ancestry, and our findings may not generalize to other populations. Extending this work to cohorts of diverse ancestries will be important for external validity and assessing broader applicability.

## Resource availability

### Lead Contact

Requests for further information and resources should be directed to Stephen C. J. Parker.

### Materials Availability

This study did not generate new unique reagents.

## Acknowledgements

The authors acknowledge several undergraduate researchers who helped cultivate and differentiate the iPSC-FAPs: Usman Siddiqui, Ashna Atukuri, Chelsea Kehoe, Benjamin Means, and Elena Wood.

We thank the Advanced Genomics Core at the University of Michigan for their assistance in generating the data used throughout this study. The project used the Microscopy and Image Analysis Core, which is supported by Grant Number P30 DK020572 (Michigan Diabetes Research Center, MDRC) from the National Institute of Diabetes and Digestive and Kidney Diseases.

CV was supported by T32GM149391 (Michigan Predoctoral Training in Genetics).

This research was supported by National Institutes of Health (NIH) grants F31DK135342, DK062370, and ZIAHG000024. The content is solely the responsibility of the authors and does not necessarily represent the official views of the NIH.

## Author contributions

CV led the experiments, analysis, and manuscript preparation. AV performed the FUSION FAP subtype identification and led the FUSION analyses used here. PO assisted with the sn-multiomic analysis including developing pipelines used. HV assisted with the iPSC-FAP nuclei QC and subtype identification. YT, AMR, TH, and SB provided essential mentorship and guidance for cultivating iPSCs and differentiating to FAPs, with SB providing additional guidance for forming the cell village and performing the insulin stimulated glucose uptake assay. SIL provided technical expertise for immunofluorescence staining and confocal microscopy. ML, JS, JT, TAL, LK, and HAK contributed biopsy samples. KLM, MB, JLS, and FSC provided supervision and mentoring. SCJP organized the project, provided mentorship, and supervised all aspects of data generation, analysis, and interpretation.

## Supplemental Information

Supplemental Table 1: All significant pathways based on KEGG enrichment analysis. Table of the significantly different pathways when comparing the basal to high-insulin environment, selected pathways shown in Figure 5G are pulled from these results. Table includes the KEGG ID, description, set size, enrichment score, NES, p-value, adjusted p-value, q-value, leading edge, and list of genes associated with those pathways. ENSG00000096696 in Figure 5H is indicated by a footnote.

Supplemental Table 2: Enriched transcription factors for peaks more accessible under high-insulin conditions.

Table of transcription factors significantly enriched within chromatin accessibility peaks that are larger under high-insulin conditions. Table includes overall ranking, motif, name, nominal p-value, log p-value, q-value, number of target sequences with the motif, percentage of target sequences with the motif, number of background sequences with motif, and percentage of background sequences with the motif. SMAD3 is indicated by a footnote (rank 31).

Supplemental Table 3: Enriched transcription factors for peaks more accessible under basal conditions.

Table of transcription factors significantly enriched within chromatin accessibility peaks that are larger under basal conditions. Table includes overall ranking, motif, name, nominal p-value, log p-value, q-value, number of target sequences with the motif, percentage of target sequences with the motif, number of background sequences with motif, and percentage of background sequences with the motif.

Supplemental Figure 1

A – UMAP of FUSION data indicating neighborhoods and colored by log fold change of differential abundance of age.

B – Heatmap of differentially expressed genes within FAP neighborhoods based on differential abundance of age.

C – UMAP of FUSION data indicating neighborhoods and colored by log fold change of differential abundance of BMI.

D – Heatmap of differentially expressed genes within FAP neighborhoods based on differential abundance of BMI.

E – UMAP of FUSION data indicating neighborhoods and colored by log fold change of differential abundance of fasting glucose.

F – Heatmap of differentially expressed genes within FAP neighborhoods based on differential abundance of fasting glucose.

G – UMAP of FUSION data indicating neighborhoods and colored by log fold change of differential abundance of fasting insulin.

H – Heatmap of differentially expressed genes within FAP neighborhoods based on differential abundance of fasting insulin.

I – UMAP of FUSION data indicating neighborhoods and colored by log fold change of differential abundance of sex.

J – Heatmap of differentially expressed genes within FAP neighborhoods based on differential abundance of sex.

K – UMAP of FUSION FAPs, colored by marker genes for the fibrogenic, adipogenic, and progenitor subclusters.

Supplemental Figure 2:

A – Immunofluorescence of iPSCs stained for two FAP markers: CD73 in green and PDGFRA in red from individual 1. Scale bar represents 20 um.

B – Immunofluorescence of iPSCs stained for two FAP markers: CD73 in green and PDGFRA in red from individual 2. Scale bar represents 20 um.

Supplemental Figure 3

A – Flow cytometry examining an iPSC marker gene and a FAP marker gene over the course of four different differentiations of a pooled sample containing ten different lines.

B – Flow cytometry examining an iPSC marker gene and a FAP marker gene over the course of one differentiation for one line.

C – Flow cytometry examining an iPSC marker gene and a FAP marker gene over the course of one differentiation for one line.

D - UCSC browser screen shot of chromatin accessibility data from the beginning (day -2), middle (day 6), and end (day 21) of a pooled sample of ten lines across the FAP differentiation as well as two individual chromatin accessibility profiles from the mix. At the bottom is the chromatin accessibility data from the FUSION FAPs (normalized counts per 10 million reads).

Supplemental Figure 4:

A – Alternate representation of the insulin stimulated glucose uptake assay.

B – Normalized luminescence from insulin stimulated glucose uptake assay for single donor in basal, basal with a small dose of insulin, high-insulin, and high-insulin with a small additional dose of insulin environments.

Supplemental Figure 5:

A – Bar plot of line representation in basal and high-insulin states as indicated by color and assessed by light whole genome sequencing of the cell village.

B – Scatterplot comparing line representation from light WGS (Figure S5A) and line representation from single-nucleus multi-omics (Figure 5B) of cell village.

C – Spearman correlation comparing *in vivo* gene expression TPM values of FAPs to individuals in iPSC-FAP cell village, sorted by representation in the cell village.

D – Gene set enrichment pathways using GO term analysis within the differential gene expression data comparing the high-insulin state to the basal state.

Supplemental Figure 6:

A – Violin plot of a marker gene of FAP progenitor subtype, p-value from MAST differential expression analysis.

B – Violin plot of a marker gene of FAP fibrogenic subtype, p-value from MAST differential expression analysis.

C – Violin plot of a marker gene of FAP adipogenic subtype, p-value from MAST differential expression analysis.

D – Basal sample pseudotime analysis with subtypes labelled. Color indicates FAP subtypes

E – Insulin sample pseudotime analysis with subtypes labelled. Color indicates FAP subtypes.

F – Table of sample composition by subtype, with binomial test p-values comparing the make-up of the subtype to the whole sample.

## Methods

### EXPERIMENTAL MODEL AND STUDY PARTICIPANT DETAILS

#### FUSION Tissue Study Individuals

The FUSION Tissue Study consists of 287 individuals aged 40 to 80 years, including 123 female participants, all of whom are of Finnish ancestry. The FUSION study was approved by the coordinating ethics committee of the Hospital District of Helsinki and Uusimaa and written informed consent was obtained from all participants.

From this cohort, a subset of 30 was selected for the cell village and iPSC experiments. The age range of this subset was 51-69 years and included 12 male and 18 female participants. We included sex as a factor in all analyses ran.

#### iPSC Growth Conditions

We grew induced pluripotent stem cell (iPSC) lines on 6 well Matrigel plates with mTeSR Plus (Stem Cell Technologies) media replaced on a daily basis. Cell lines were passaged once a week by incubating the wells in 1 mL of Versene and either incubating at room temperature for ten minutes or 37 °C for 5 minutes. Cells were then scraped off of the plates and added at a 1:3 ratio to the new plates. Cells were incubated at 37 °C, 5% CO2 overnight.

### QUANTIFICATION AND STATISTICAL ANALYSIS

P-values are reported within figures, with values <0.05 indicating statistical significance for correlations and differences. Details of statistical methods used and sample sizes are provided in the figure legends, and plots display mean ± SEM. Sex was included as a covariate in all analyses.

### METHOD DETAILS

#### Percent FAP per skeletal muscle biopsy analysis

In order to compare changes in the number of nuclei per FAPs in relation to donor traits in the FUSION Tissue study skeletal muscle biopsies, DESeq2 (v. 1.42.1) was used. We used raw nucleus counts per cell type as input and the abundance of FAPs was then compared to donor metadata variables. The model included batch, sex, BMI, age, T2D status (all donors were encoded as a one if diagnosed with T2D and zero otherwise), fasting glucose, and fasting insulin.

#### Neighborhood analysis

To perform neighborhood analyses, MiloR (v. 2.1.3) was used.^22^ The model used included age, BMI, batch, fasting glucose at time of biopsy, fasting insulin at time of biopsy, T2D status (1 if an individual was diagnosed with T2D at time of biopsy, 0 otherwise), and sex. In order to identify differentially expressed genes between neighborhoods, we used Milo’s function testDiffExp and pinpointed genes whose expression levels were associated with the metadata of interest. The testDiffExp function tests this association by using a negative binomial GLM from edgeR. Milo’s function plotNhoodExpressionGroups was then used to plot the top ten most significantly differently expressed genes (based on adjusted p-values) that differ between neighborhoods with differential abundance within the FAPs.

#### Differential Chromatin Accessibility Across *in vivo* FAP Subtypes

Differential peaks in Figure 1J were compared using DESeq2 (v. 1.42.1). Once the FAP subtypes were identified, we generated a consensus peak set by merging the narrowPeak files from all three subtypes. Specifically, we used bedtools^48^ (v. 2.30.0) to consolidate overlapping features into a single region that encompasses the combined peaks. We then used featureCounts^49^ (v. 2.0.3) to quantify the read counts in peaks per individual per subtype. This was used as input for DESeq2, where a paired analysis was performed, and the *MME* promoter peak was examined.

#### Differentiation of iPSC-FAPs

We differentiated iPSCs to FAPs by plating and growing as iPSCs for 2 days (day -2 to day 0). Mesenchymal Induction media (Stem Cell Technologies) was then added, and cells were grown in that for 4 days (day 0 to day 4). After that cells were then grown in FAP media (86% alpha-MEM, 13% FBS, and 1% L-glutamine). On day 6, cells are passaged to MatrixPlus plates (StemBioSys). Cells are grown in FAP media and on MatrixPlus plates until day 21 when they are then considered to be iPSC-FAPs. FAPs are passaged using the ACF Cell Dissociation Kit (Stem Cell Technologies).

#### Immunofluorescence of iPSC-FAPs and iPSCs

##### Staining

To prep for immunofluorescence staining, cells were grown on glass coverslips overnight. Cells were then washed with PBS and fixed in 3% PFA for ten minutes on a shaker before being washed again with PBS three times for five minutes each wash. The cells were then blocked with 5% BSA in 1x PBS + 0.1% Triton X-100 for 1 hour at room temperature before adding primary antibody diluted in 5% BSA in 1x PBS + 0.1% Triton X-100 for two hours at room temperature. The antibody for *CD73* (Thermo Fisher Scientific, Cat# 41-0200, RRID:AB_2533492) was diluted to 1:50 while *PDGFRA* (Thermo Fisher Scientific, Cat# 710169, RRID:AB_2532600) was diluted to 1:500. Cells were then washed with 1x PBS + Triton X-100 3 times for 6 minutes each before secondary antibody incubation for one hour at room temperature. Secondary antibodies goat anti-rabbit 488 (Thermo Fisher Scientific Cat# A-11008, RRID:AB_143165) and goat anti-mouse 594 (Thermo Fisher Scientific Cat# A-11005, RRID:AB_2534073) were diluted to 1:2000 in 5% BSA in 1x PBS + 0.1% Triton X-100. The plate with the coverslips was covered in foil to preserve the fluorescence. Cells were then washed with 1x PBS + Triton X-100 3 times for 6 minutes each. After that cells were washed in PBS alone for five minutes, PBS with a 1:1000 dilution of DAPI for five minutes, and PBS alone for another five minutes. Coverslips were then mounted on slides using Prolong Gold Antifade mounting media before storing at 4C.

##### Imaging

Immunofluorescence images were acquired on a Nikon A1 confocal with a 40x/1.3 NA oil immersion objective. Identical settings were used for all fluorescence microscopy and post-acquisition adjustments to brightness and contrast. Images were cropped to isolate individual cells.

#### FACS analysis

For flow cytometry analysis, we dissociated cells using the previously mentioned passaging methods. Cells are put through a 70 um filter and divided into a negative control and a stained sample. Cells are then centrifuged down, media is aspirated, and the positive sample is stained with antibodies for *Tra-1-60* (iPSC marker) (Miltenyi Biotec Cat# 130-122-921, RRID:AB_2801969) and *NT5E* (FAP marker) (Miltenyi Biotec Cat# 130-111-913, RRID:AB_2784275) for 10 minutes. Cells are centrifuged down and then fixed in 3% PFA for 15 minutes before getting resuspended in PBS. All reagents are kept on ice throughout until FACS data collection.

#### Bulk RNA-sequencing

##### Data generation

We isolated RNA for bulk RNA-sequencing from iPSC-FAPs using the AllPrep DNA/RNA Mini Kit (Qiagen). In brief, cells were frozen in aliquots of 500,000 cells that were then thawed and centrifuged for 5 minutes at 300g. The supernatant was then removed and 600 ul of Buffer RLT Plus (from kit) was added before vortexing for one minute. Cells were transferred to an AllPrep DNA spin column and centrifuged for 30 seconds at 8000 x g. The supernatant was then used for RNA isolation. To the supernatant, 600 ul of 70% ethanol was added before transfer of 700 ul to an RNeasy spin column. This was centrifuged at 8000 x g for 15 seconds and flow through was discarded. Next 700 ul of Buffer RW1 (from kit) was added before being centrifuged at 8000 x g for 15 seconds and flow through was discarded. Then 500 ul of Buffer RPE (from kit) was added before being centrifuged at 8000 x g for 15 seconds and flow through was discarded. Again 500 ul of Buffer RPE (from kit) was added before being centrifuged for two minutes at 8000 x g. The spin column was then placed in a new 1.5 mL collection tube and 50 ul of RNase-free water was added to the membrane. This was centrifuged for one minute at 8000 x g before submission to the sequencing core.

##### Data analysis

Bulk RNA-sequencing data was analyzed to examine marker gene expression levels. In brief, we trimmed sequencing adapters using cta (v. 0.1.2) and mapped reads to hg38 using STAR (v. 2.7.10). featureCounts (v. 2.0.3) was then used to quantify reads per GENCODE annotated gene before calculating TPM of selected marker genes.

#### Nuclei isolation and snATAC data generation

We pooled ten iPSC lines together for a joint differentiation and split the cells into two replicates after day 6. Cells were frozen down into cryovials of 500,000 cells. For nuclear isolation, cells were thawed and pelleted at 500 RCF at 4°C for 5 min in a fixed-angle centrifuge. Cells were then resuspended in 50 ul of ATAC-Resuspension Buffer (RSB) (1M Tris-HCl pH 7.4, 5M NaCl, 1M MgCl2, and water to 50 mL) containing 0.1% NP40, 0.1% Tween-20, and 0.01% Digitonin and pipetted up and down 3 times before incubating on ice for 3 minutes. ATAC-RSB was then washed out with 1 mL of ATAC-RSB containing 0.1% Tween-20, and tube was inverted 3 times to mix. Nuclei are then pelleted at 500 RCF for 10 min at 4°C in a fixed-angle centrifuge, then resuspended in 1000ul PBS + 0.1% Tween-20 and 1% BSA with a 1:1000 dilution of 7-AAD and incubated for 30 minutes on ice before sorting with a Miltenyi Tyto machine. Five thousand nuclei were then loaded onto the 10x platform in two replicates to generate snATAC data using the 10x multi-omics kit.

#### Comparing chromatin accessibility profiles with a Jaccard statistic

To compare chromatin accessibility peaks between the iPSC-FAPs and *in vivo* FAPs, we used bedtools^48^ (v. 2.30.0) function “bedtools jaccard” to calculate the Jaccard similarity statistic. For this analysis, we restricted the iPSC-FAPs to the top 200,000 strongest peaks based on p-value ranking within the broadPeak file and defined intersection as peaks overlapping at least 75% of the iPSC-FAP input peak (bedtools jaccard -f 0.75).

#### Comparing chromatin accessibility profiles with a logistic regression

We used a logistic regression approach to compare cell-type-specific chromatin accessibility profiles between iPSC-FAPs from the eight most abundant individuals at the end time point and corresponding peaks from the cell types from the FUSION skeletal muscle biopsies. For each of the eight individuals, we took the day -2 peaks (called separately on each individual) and day 21 peaks (called separately for each individual) and generated a master peak list by merging them (taking the union, peaks called using MACS2 v. 2.2.4). We then filtered to TSS-distal master peaks (peaks not overlapping the 5kb window upstream of a gene) and restricted to master peaks present in only one of the two timepoints. Next, for each of the 13 FUSION cell types, we fit a logistic regression model:

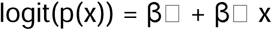

where p(x) is the probability that the master peak is accessible at day 21, and x is the probability that the master peak overlaps a peak from the fusion cell type. We then divided the 13 coefficients by the maximum coefficient to obtain get a peak “similarity score” capped at 1.

#### Insulin-stimulated glucose uptake assay

To investigate the response of the FAPs to insulin, we used an insulin-stimulated glucose uptake assay. To do this, we incubated cells in 100 mM of insulin for two hours or kept them at basal levels (constituting the basal and high-insulin samples) prior to using the Promega insulin-stimulated glucose uptake kit. Each line was measured in triplicate in both of the stimulatory states. Lines were plated in replicates on four 96-well plates (20 lines were assessed in triplicate, 8 in duplicate, and 2 as singlets based on cell availability at the time). Within each environment, one plate was given an additional dose of 100 nM of insulin for 30 minutes before all wells were given fluorescent glucose, and luminescence was measured to assess glucose uptake, allowing us to compare glucose uptake in response to insulin under basal and high-insulin conditions.

##### Statistical analysis

After measuring luminescence levels, we subtracted the background (averaged from empty wells) from each well. We normalized luminescence levels to the number of cells, measured at plating, and performed inverse rank normalization. We used a paired t-test to assess the significance of the difference in fold change between samples that received additional insulin from the Promega kit or not.

##### Response to stimulatory state modeling

To compare normalized luminescence values to donor metadata variables, we used a linear mixed model. The model included age, BMI, differentiation efficiency (from the FACS analysis of that specific line’s differentiation), sex, and T2D status.

#### DNA isolation

We isolated DNA for light whole genome sequencing from cell lines using the DNeasy Blood & Tissue Kit (Qiagen). In brief, cells were frozen down in aliquots of 500,000 cells that were then thawed and centrifuged for 5 minutes at 300g. Pellet was resuspended in 200 ul PBS and Proteinase K (from kit). 200 ul of Buffer AL (from kit) was added before vortexing and incubating at 56 C for 10 minutes. 200 ul of ethanol was added before vortexing and adding the mixture to a DNeasy spin column. The spin column was centrifuged at 6000g for 1 pm, flow-through was discarded, and the spin column was placed in a new collection tube. 500 ul of Buffer AW1 (from the kit) was added before centrifuging for 1 minute at 6000g, flow through was again discarded, and a new collection tube was added. 500 ul of Buffer AW2 (from kit) was then added, and centrifuged for 3 minutes at 20,000g, flow-through was again discarded. The spin column was then placed in a 1.5 mL tube, 200 ul of water was added before incubating at room temperature for 1 minute, and then centrifuging at 6000g for 1 minute to elute the DNA.

#### Census-seq

For census-seq (v. 2.5.1) analysis,^50^ we trimmed reads using cta (v. 0.1.2, https://github.com/ParkerLab/cta), then aligned to the hg38 reference genome using BWA (v. 0.7.17). We then marked duplicates using Picard Tools MarkDuplicates (v. 3.4.0, https://broadinstitute.github.io/picard/) and removed them prior to running Census-Seq, Roll Call, and CSI Analysis from the Census-Seq tools.^50^ Specifically, census-seq’s Roll Call analysis was used to determine the fraction match of each individual iPSC line’s low pass whole genome sequencing to reference SNP Chip and Imputation data for each donor. This tool compares individual reads from whole-genome sequencing to the reference data for each donor and determines a fraction match based on how many reads of the whole genome sequencing match the reference.

#### Nuclei isolation and multi-omics data generation

The cell village was frozen down into aliquots of 200,000 cells. The cells were thawed on ice for 30 minutes before being transferred to 1 mL of pre-warmed alpha-MEM and centrifuged at 300 rcf for 5 minutes. The cells were then resuspended in 1 mL DPBS with 0.04% BSA before proceeding. To isolate nuclei, cells were pelleted at 500 rcf at 4 C for 5 minutes in a fixed-angle centrifuge. 600 ul of lysis buffer (10 mM Tris-HCl pH 7.4, 10 mM NaCl, 3 mM MgCl2, 0.1%

NP40, 1% of 10% BSA, 0.1% Tween-20, 1 mM DTT, 50 mg/mL 7-AAD, 1 U/ul RNase inhibitor, and water to volume) was added before transferring to a pre-chilled dounce homogenizer and homogenizing with 5 strokes. The mixture was then incubated on ice for 7 minutes, with pipetting once per minute. Lysis was then washed out with 1.3 mL of wash buffer (1% of 10% BSA, 0.1% Tween-20, 1 mM DTT, 0.5 mg/mL 7-AAD, and DPBS to volume) before pelleting at 500 rcf for 5 minutes at 4 C in a swinging bucket centrifuge. Nuclei were resuspended in 0.1% Tween-20, 1% BSA, 1 mM DTT, 1 U/ul RNase Inhibitor, 1 mg/mL 7-AAD, and DPBS to the desired volume then incubated for 15 minutes. Due to issues with the Miltenyi Tyto machine, the basal sample was sorted through the Tyto, and the insulin sample was pelleted at 500 rcf for 5 minutes at 4 C in a swinging bucket centrifuge, then the nuclei were resuspended in 450 ul of wash buffer before spinning down at 500 rcf for 5 minutes in a fixed-angle centrifuge. The resuspension in wash buffer and the spin-down was repeated once more. Finally, both samples were resuspended in 1 mL nuclear permeabilization buffer (10 mM Tris-HCl pH 7.5, 10 mM NaCl, 3 mM MgCl2, 0.1% Tween-20, 0.1% IGEPAL, 0.01% Digitonin, and water to volume) before pipetting 10 times. From there, 10,000 basal nuclei and 20,000 insulin nuclei were loaded onto the 10x Genomics multi-omics kit for data generation.

#### Multi-omics analysis

##### snATAC-seq processing

The chromatin accessibility (ATAC) component of the multiome library was processed using the pipeline at https://github.com/porchard/snATACseq-NextFlow (commit b81ce91) and the TOPMed version of the hg38 reference genome (described here: https://github.com/broadinstitute/gtex-pipeline/blob/master/TOPMed_RNAseq_pipeline.md). In brief, we trimmed sequencing adapters using cta (v. 0.1.2) and mapped reads to hg38 with bwa^51^ (v.0.7.15; bwa mem -I 200,200,5000 -M). Cell barcodes were corrected using a custom implementation of the barcode correction algorithm described on the 10X Genomics website. Duplicates were marked using picardtools (http://broadinstitute.github.io/picard/; v 2.26.9; MarkDuplicates READ_ONE_BARCODE_TAG=CB READ_TWO_BARCODE_TAG=CB VALIDATION_STRINGENCY=LENIENT). BAM files were filtered to properly mapped, non-duplicate autosomal read pairs using samtools (samtools view -h -b -f 3 -F 4 -F 8 -F 256 -F 1024 -F 2048 -q 30 chr{1..22}). Peaks were called using MACS2^52^ (macs2 callpeak -t $bed --outdir . --SPMR -f BED -g hs --nomodel --shift -100 --extsize 200 -B --broad --keep-dup all, with a bed file generated using the bedtools^48^ bamtobed command) and filtered against the ENCODE blacklist (https://www.encodeproject.org/files/ENCFF356LFX/). QC metrics were generated using ataqv^53^ (v. 1.5.0; --ignore-read-groups --nucleus-barcode-tag CB, with the ENCODE blacklist.

##### snRNA-seq processing

The gene expression (RNA) component of the multiome library was processed using the pipeline at https://github.com/porchard/snRNAseq-NextFlow (commit baea274). In brief, reads were mapped and gene count matrices generated using STARsolo^54^ (v. 2.7.10a, with hg38 fasta file and GENCODE v30-derived genome annotations from the TOPMed consortium (https://github.com/broadinstitute/gtex-pipeline/blob/master/TOPMed_RNAseq_pipeline.md); default options except --soloBarcodeReadLength 0 --outSAMattributes NH HI nM AS CR CY CB UR UY UB sM GX GN --outSAMtype BAM SortedByCoordinate --outSAMunmapped Within KeepPairs --soloType CB_UMI_Simple --soloUMIlen 12 --soloFeatures GeneFull_ExonOverIntron --soloUMIfiltering MultiGeneUMI --soloCBmatchWLtype 1MM_multi_pseudocounts --soloCellFilter None --soloCBwhitelist 10X-whitelist.txt). We removed ambient RNA using cellbender ^55^ (v. 0.3.0, options --fpr 0.05) and emptyDrops^56^, and calculated per-nucleus QC metrics using a custom python script.

##### Multiome QC

After processing the ATAC and RNA components as described above, we selected pass-QC nuclei by applying the following thresholds for each nucleus. For the basal sample, we required a minimum of 499 UMIs for the RNA modality, maximum 15 percent mitochondrial reads from the RNA modality, minimum 3000 properly paired and mapped non-duplicate autosomal reads where mapping quality is at or above 30 in the ATAC modality, minimum TSS enrichment of 2 in the ATAC modality, and minimum CellBender cell probability of 0.99 for the RNA modality. For the high-insulin sample, we required a minimum of 1445 UMIs for the RNA modality, maximum 15 percent mitochondrial reads from the RNA modality, minimum 6672 properly paired and mapped non-duplicate autosomal reads where mapping quality is at or above 30 in the ATAC modality, minimum TSS enrichment of 2 in the ATAC modality, and minimum CellBender cell probability of 0.99 for the RNA modality.

##### Multiome doublet detection

We relied on both the RNA and the ATAC modalities to detect doublets. The multiome library’s doublets detection was processed using the pipeline at https://github.com/porchard/Multiome-Doublet-Detection-NextFlow (commit 72df9ea). For both modalities we ran demuxlet^57^ for detecting doublets and assigning singlets to donors. If a barcode was found to be a doublet based on either method or modality (or if the inferred donor identity differed between the ATAC and RNA modalities), it was removed prior to clustering. Clustering was performed using Seurat v5.^58^ After subsetting to pass-QC nuclei and adding metadata, the nuclei were split into different layers based on sample (basal or insulin). scTransform was then used to normalize counts (<u>vars.to</u>.regress = c(“donor”, “age”, “sex”, “bmi”, “ogtt”). The layers were then integrated using Harmony. 10 PCs were included in the final clustering (any more would lead to donor-specific clusters) with a resolution of 0.5 used in FindClusters. An additional separate cluster of less than 100 nuclei was additionally removed to limit analyses to strictly FAPs. A binomial test was used to assess the composition of different subtypes for significance.

#### Pseudotime analysis

To assess pseudotime trajectories in the gene expression data of the iPSC-FAPs, Monocle3 (v. 1.3.7) was used.^59^

#### Spearman correlation

To assess the similarity of the iPSC-FAPs to FUSION skeletal muscle cell types, TPM values were calculated from average counts per gene for each cell type. The comparison was then restricted to genes shared between the cell types, and a Spearman correlation was used to quantify similarities.

#### Differential gene expression

Differential gene expression was performed using MAST (v. 1.28.0) using a model that took into account donor metadata traits of sex, age, BMI, and T2D status.^60^ Genes expressed in less than 0.35 of nuclei were removed prior to analysis. Pathway analysis was then conducted by clusterProfiler (v. 4.16.0).^61^

#### Differential peak analysis

##### Comparing basal vs high-insulin

To identify differential chromatin accessibility peaks between the basal and high-insulin samples, a Negative Binomial Generalized Linear Mixed Model (NB-GLMM) was used. First, peak lists were compiled using snapATAC2 (v. 2.7.0).^62^ Differential peaks were then identified by the NB-GLMM controlling for donor traits and testing for basal vs insulin samples (called “sample” in the below models) at an FDR level of 5% (using Storey Lab’s qvalue package, v. 2.15.0). Two models were ran, both with and without the variable of interest:

model_with <- as.formula(“PEAK ∼ sample + sex + age + ogtt + bmi + tss_enrichment + offset(log(total_counts)) + (1|donor)”)

model_without <- as.formula(“PEAK ∼ sex + age + ogtt + bmi + tss_enrichment + offset(log(total_counts)) + (1|donor)”)

After running both models for each peak, likelihood ratio tests (LRTs) identified differentially accessible peaks between the basal and high-insulin samples.

##### Comparing iPSC-FAP subtypes

To identify differential chromatin accessibility peaks between subtypes, we used a NB-GLMM framework where:

model_with <- as.formula(“PEAK ∼ subtype + sex + age + ogtt + bmi + tss_enrichment + offset(log(total_counts)) + (1|donor)”)

model_without <- as.formula(“PEAK ∼ sex + age + ogtt + bmi + tss_enrichment + offset(log(total_counts)) + (1|donor)”)

Comparing the two models with LRT identified peaks significantly associated with FAP subtype (FDR < 5%, Storey qvalue package, v. 2.15.0). For these subtype-associated peaks, we extracted pairwise subtype coefficients and corresponding Wald test P values (null hypothesis = 0) from the full model. Given three FAP subtypes, the full model yielded two coefficients representing the accessibility of each subtype relative to a reference; we then releveled the data to obtain the third coefficient comparing the two non-reference subtypes directly. A peak was designated as subtype-specific if it showed a positive coefficient and Wald test P < 0.05 in both comparisons involving that subtype, and a non-significant Wald P (> 0.05) for the comparison between the other two subtypes. For example, a peak was called adipogenic-specific if it had positive coefficients and P < 0.05 for adipogenic vs. fibrogenic and adipogenic vs. progenitor comparisons, but P > 0.05 for fibrogenic vs. progenitor. This two-step process allowed us to first identify overall peaks impacted by subtype and then those specific to each subtype.

### Motif enrichment analysis

Transcription factor motif enrichment was performed using HOMER (v. 4.11.1).^63^ The command findMotifsGenome.pl was used with default settings separately on bed files of peaks more accessible under basal and high-insulin conditions.

### GWAS Enrichment analysis

To perform enrichment analyses comparing chromatin accessibility peaks of the iPSC-FAP subtypes to GWAS data, fgwas (v. 0.3.6) was used.^45^ In this case, summary statistics from the 2024 T2D GWAS (specifically the multi-ancestry analysis) were used for the enrichment.^4^ The summary statistics were formatted as required by the program, and default parameters were used.

#### Chromatin looping analysis

To identify genes our peak of interest was likely interacting with, we used SCENT (v. 1.0.1).^64^ Genes within 50 kb of the selected peak were tested with covariates log(UMI), fraction mitochondrial reads, sample (basal/insulin), donor, sex, BMI, T2D status, and age.

