## Supplementary figures and images for "Donor-matched iPSC model reveals context-dependent T2D genetic signals in fibro-adipogenic progenitors"

### Supplemental Figure 1

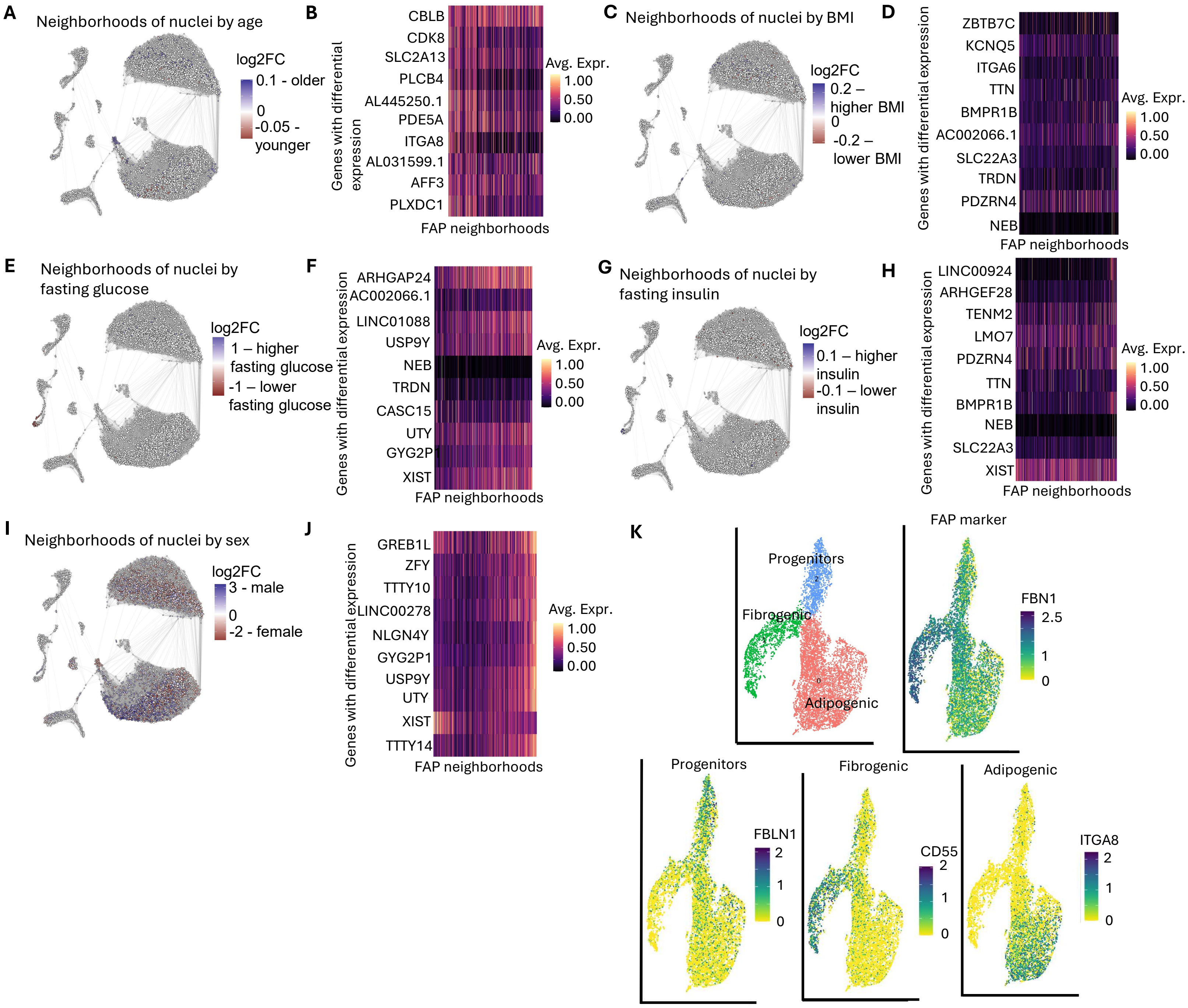

### Supplemental Figure 2

Day -2 iPSC Immunofluorescence

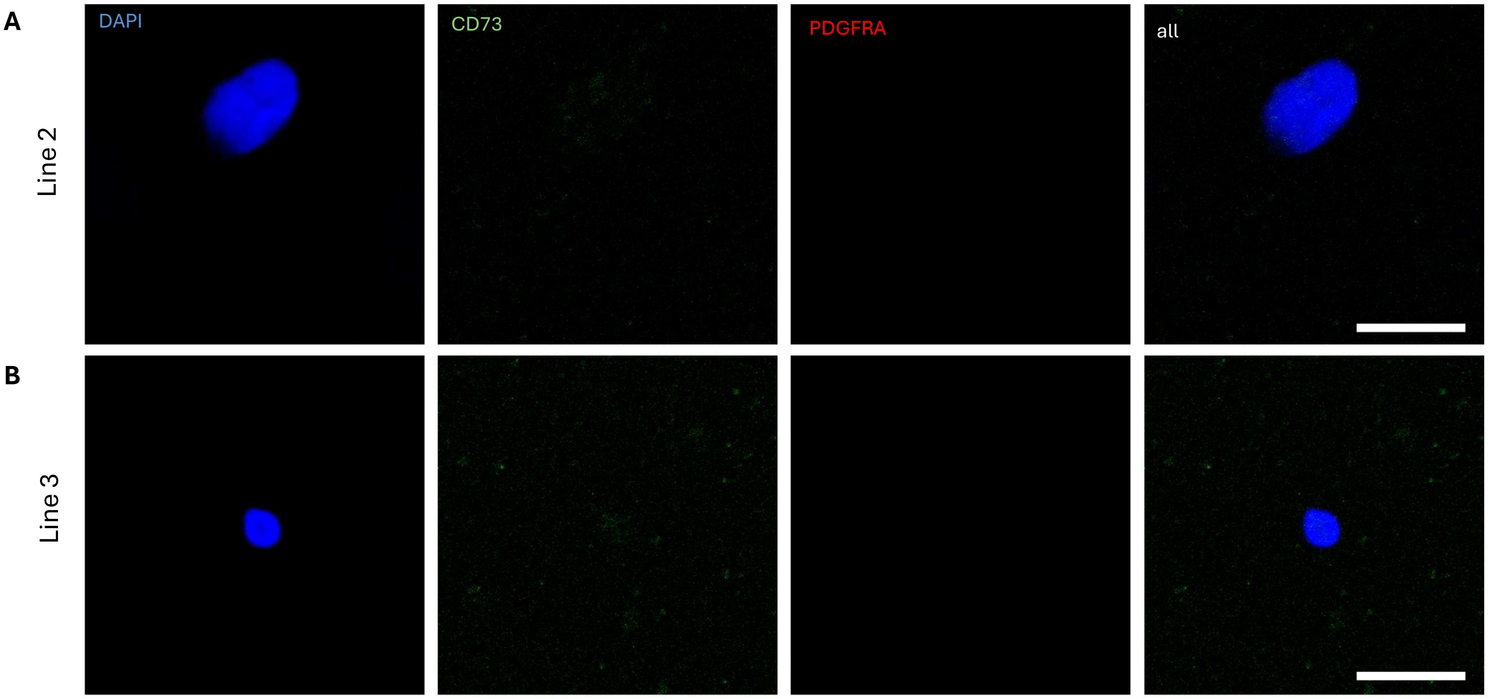

### Supplemental Figure 3

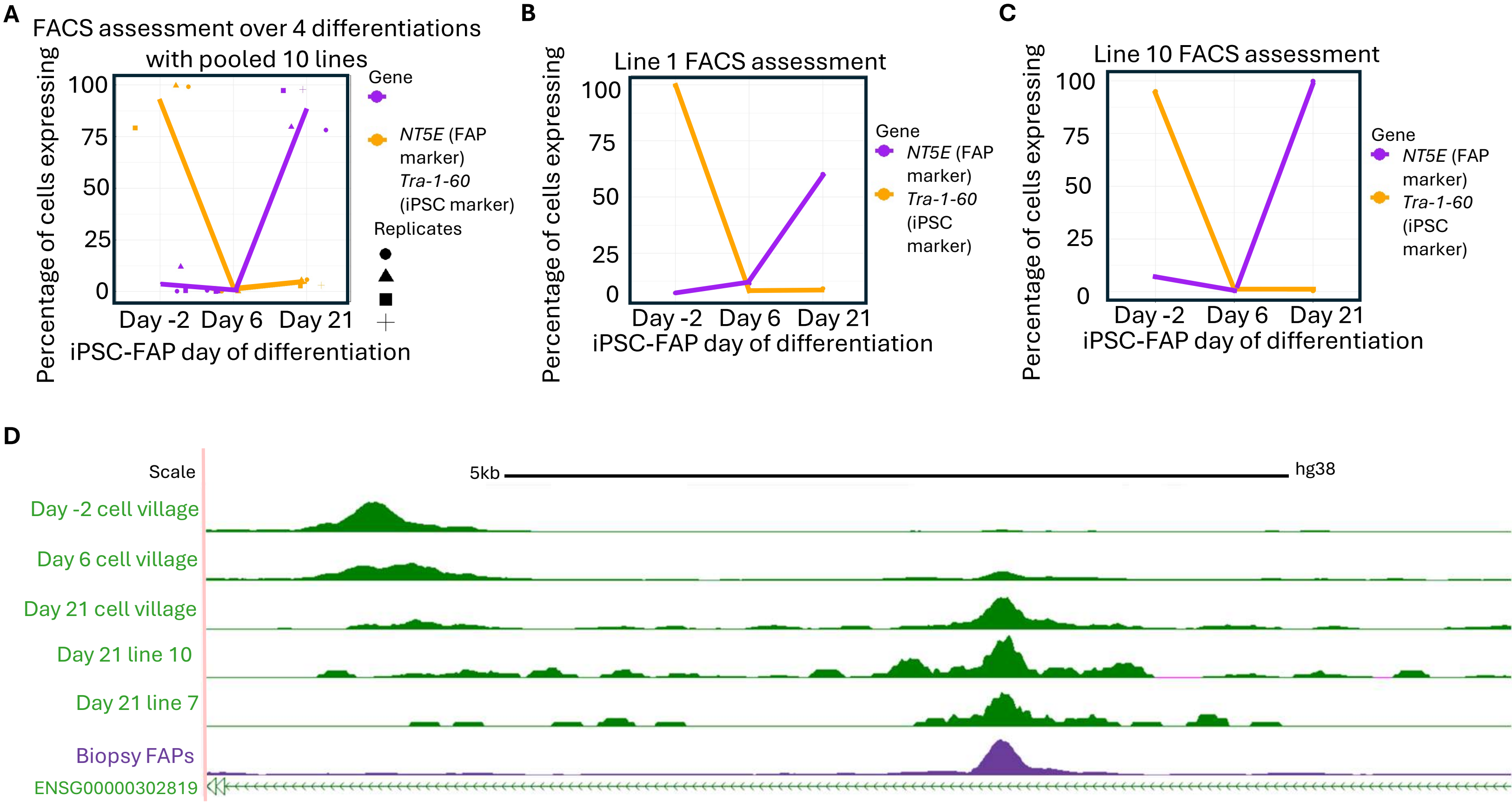

### Supplemental Figure 4

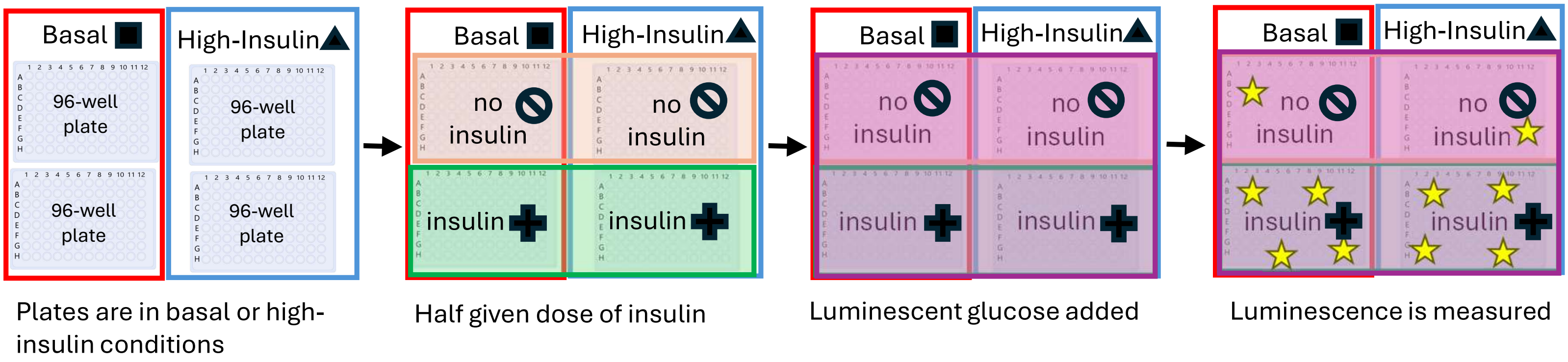

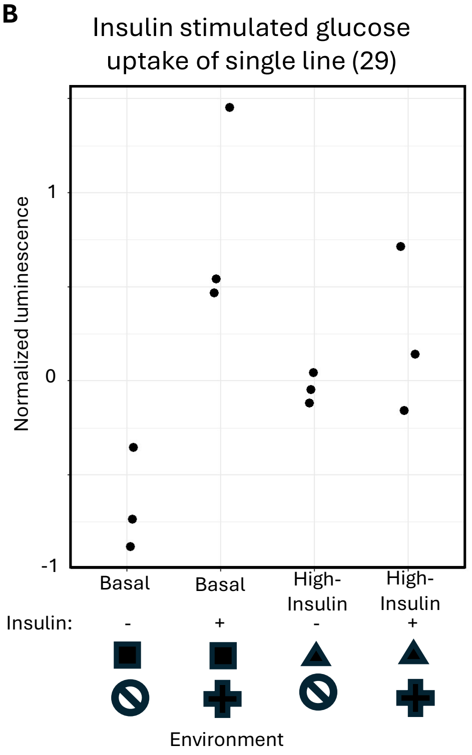

### Supplemental Figure 5

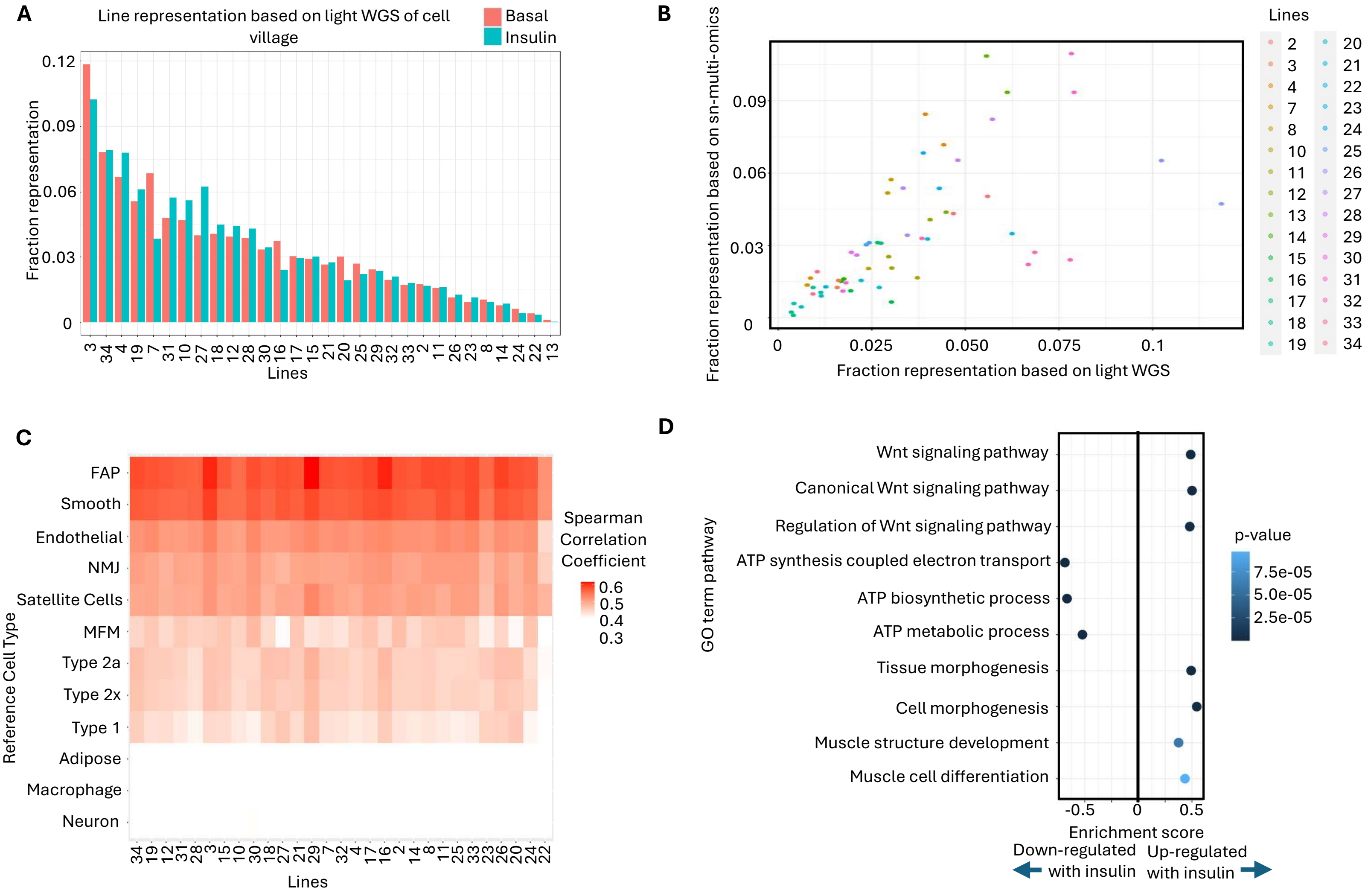
