## Supplemental Figure 6 for "Donor-matched iPSC model reveals context-dependent T2D genetic signals in fibro-adipogenic progenitors"

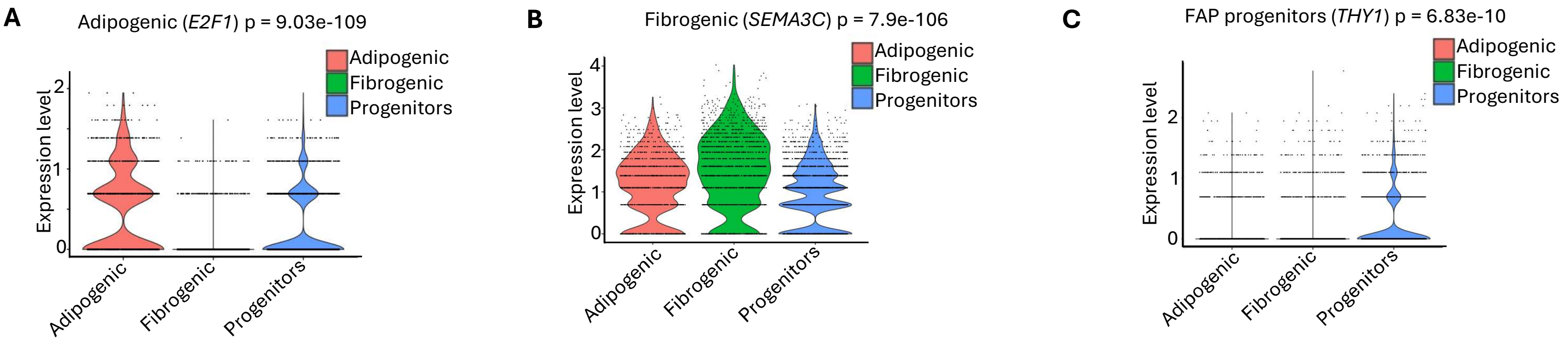


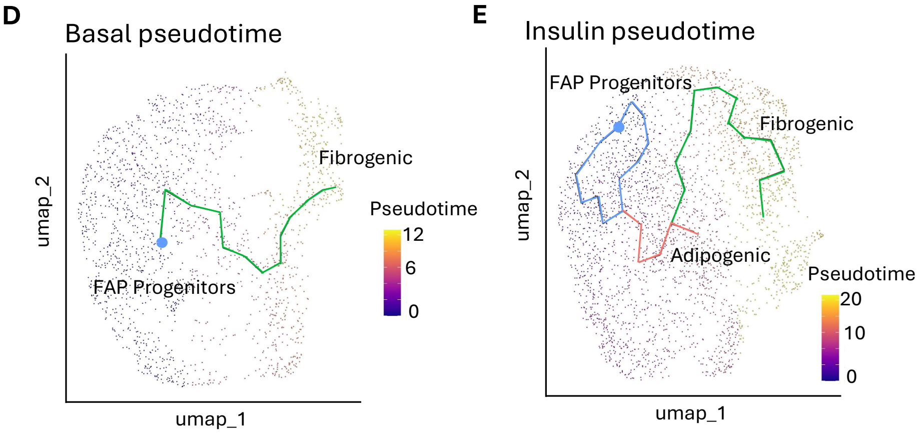


**F**

| **Subtype** | **Basal nuclei** | **Insulin nuclei** | **Binomial test p-value** |
|---|---|---|---|
| Progenitors | 0.43 (897) | 0.57 (1,208) | 0.015 |
| Fibrogenic | 0.42 (710) | 0.58 (992) | 0.151 |
| **Adipogenic** | **0.31 (383)** | **0.69 (838)** | **4.478e-10** |
| *Total* | *0.4 (1,990)* | *0.6 (3,038)* | -- |
