## Supplemental Table 1 for "Donor-matched iPSC model reveals context-dependent T2D genetic signals in fibro-adipogenic progenitors"

| ID | Description | setSize | enrichmentScore | NES | pvalue | p.adjust | qvalue | leading_edge | core_enrichment |
| --- | --- | --- | --- | --- | --- | --- | --- | --- | --- |
| hsa03010 | Ribosome | 132 | -0.845288971768744 | -3.00835282917901 | 1e-10 | 4.01428571428571e-09 | 3.03759398496241e-09 | tags=83%, list=12%, signal=75% | 122704/200916/64983/9801/6191/6122/10399/29088/6166/6188/6128/51065/6203/6124/4736/6135/6125/6139/64949/6138/3921/9045/6132/2197/6175/6130/6209/57129/63875/51023/6129/28973/23521/6201/6146/6187/6141/219927/6156/6194/6189/6229/64968/6137/11224/6164/6202/6232/6205/6169/6142/6152/6155/6224/51073/6173/6233/6143/6158/6222/6193/6134/65005/6234/6136/6157/9349/6223/6133/51081/7311/6144/6168/6230/6217/6207/6208/6159/6176/6204/6154/25873/6181/6161/6160/6228/79590/6165/6170/29093/6227/6167/6231/6235/6210/51121/6147/6171/6218/9553/6206/84545/51253/51649/51264/29074/64979/6183/64928/51021 |
| hsa05171 | Coronavirus disease - COVID-19 | 117 | -0.805118690227416 | -2.80413660539623 | 1e-10 | 4.01428571428571e-09 | 3.03759398496241e-09 | tags=68%, list=9%, signal=64% | 6166/6188/6128/51065/6203/6124/4736/6135/6125/6139/6138/3921/9045/6132/2197/6175/6130/6209/6129/23521/6201/6146/6187/6141/6156/6194/6189/6229/6137/11224/6164/6202/6232/6205/6169/6142/6152/6155/6224/6173/6233/6143/6158/6222/6193/6134/6234/6136/6157/9349/6223/6133/7311/6144/6168/6230/6217/6207/6208/6159/6176/6204/6154/25873/6181/6161/6160/6228/6165/6170/6227/6167/6231/6235/6210/51121/6147/6171/6218/6206 |
| hsa00190 | Oxidative phosphorylation | 105 | -0.675251349596165 | -2.3250876698649 | 1e-10 | 4.01428571428571e-09 | 3.03759398496241e-09 | tags=71%, list=23%, signal=56% | 4714/51079/1537/7385/10476/506/529/5464/4707/4694/513/54539/55967/9377/4713/1329/4700/7386/4711/4708/4724/8992/4718/4704/539/4716/54205/4706/4726/4715/4720/1327/9551/522/516/4710/126328/1340/29796/4712/4723/4697/1345/1337/374291/517/521/4725/27089/518/7381/533/7388/4709/10975/10632/1350/9550/1351/1347/4696/4728/514/509/4695/9296/4702/4698/6390/1349/4722/9167/4717/10063/4729 |
| hsa05012 | Parkinson disease | 187 | -0.580069389099006 | -2.13765698150589 | 1e-10 | 4.01428571428571e-09 | 3.03759398496241e-09 | tags=59%, list=24%, signal=47% | 7417/7345/581/5685/5690/5714/4714/51079/1537/5684/7385/6647/10476/506/7494/1965/5689/5691/4707/4694/513/54539/5682/7332/55967/9377/5692/4713/1329/4700/5694/7386/5683/2773/4711/203068/4708/10383/4724/292/4718/5693/5719/4704/539/4716/2778/54205/4706/4726/4715/4720/801/1327/5701/11315/522/84790/293/516/4710/7314/126328/25828/10131/10376/1340/29796/7979/4712/7295/4723/4697/1345/5688/1337/374291/517/4725/27089/518/7381/7388/4709/805/118424/10975/1350/1351/1347/4696/4728/514/9817/5715/6233/509/4695/10381/27173/4702/7311/4698/6390/7419/5686/1349/4722/9167/4717/4729 |
| hsa05020 | Prion disease | 177 | -0.565892599102523 | -2.07484814475183 | 1e-10 | 4.01428571428571e-09 | 3.03759398496241e-09 | tags=58%, list=24%, signal=46% | 7417/3304/581/5685/5690/5714/4714/51079/1537/5684/7385/6647/10476/506/1965/5689/5691/4707/4694/513/54539/5682/858/55967/3303/9377/5692/4713/1460/1329/4700/5694/7386/5683/857/4711/5534/203068/4708/10383/4724/292/4718/5693/5719/4704/539/4716/54205/4706/4726/4715/4720/5879/1327/5701/572/522/84790/293/516/4710/126328/3312/10376/1340/29796/7979/4712/4723/4697/1345/5688/1337/374291/517/4725/27089/518/7381/7388/4709/10975/1350/1351/1347/4696/4728/514/5715/509/4695/10381/4702/4698/6390/7419/5686/1349/4722/9167/4717/4729 |
| hsa05014 | Amyotrophic lateral sclerosis | 232 | -0.548152521914591 | -2.07058819487786 | 1e-10 | 4.01428571428571e-09 | 3.03759398496241e-09 | tags=52%, list=24%, signal=42% | 581/5685/5690/5714/4714/9818/51079/3178/311/1537/5684/79902/7385/6647/10476/506/7494/1965/29978/71/5689/5691/55626/10189/4707/4694/513/23279/54539/5682/55967/9377/5692/4713/1329/4700/637/5694/7386/5683/4711/5534/203068/4708/10383/4724/642659/4718/5693/5719/4704/6428/539/4716/5861/54205/4706/4726/4715/4720/5879/1327/5701/572/522/8480/84790/516/4710/2876/126328/10121/10376/10540/1340/29796/7979/4712/55706/4723/4697/1345/310/5688/1337/374291/517/4725/5216/27089/518/7381/7388/4709/10975/4218/5217/81631/1350/1351/1347/4696/11258/4728/10280/514/5715/509/4695/10381/4702/10671/10762/4698/6390/5686/1349/4722/9167/4717/4729 |
| hsa05016 | Huntington disease | 197 | -0.551031225481673 | -2.04646618132718 | 1e-10 | 4.01428571428571e-09 | 3.03759398496241e-09 | tags=56%, list=24%, signal=44% | 581/5685/5690/5714/4714/51079/1537/5684/7385/6647/10476/506/5689/5691/55626/4707/4694/513/54539/3065/5682/55967/9377/5692/4713/1173/5435/1329/4700/5694/7386/5683/160/4711/203068/4708/10383/4724/292/4718/5693/5719/4704/539/4716/54205/4706/4726/4715/4720/1327/5701/522/7019/84790/293/516/1212/5437/4710/2876/126328/10121/10376/10540/1340/1175/29796/7979/4712/4723/4697/1345/5688/1337/374291/517/4725/27089/518/7381/7388/4709/10975/1350/1351/1347/4696/11258/4728/514/5715/5441/509/4695/10381/4702/10671/4698/6390/7419/5686/1349/4722/5436/9167/5440/5439/4717/4729 |
| hsa04714 | Thermogenesis | 142 | -0.560465016989588 | -2.01210486105953 | 6.5001858468378e-09 | 2.28319027870178e-07 | 1.72768097508057e-07 | tags=56%, list=23%, signal=44% | 4714/6602/51079/1537/7385/10476/506/3265/71/4707/4694/513/54539/55967/9377/4713/1329/4700/7386/4711/284184/4708/4724/4718/4704/539/4716/2778/4706/4726/4715/4720/1327/9551/522/516/4893/4710/126328/1340/29796/4712/4723/4697/1345/1337/374291/86/517/521/4725/27089/518/7381/7388/4709/10975/6598/10632/1350/1351/25915/1347/4696/4728/6194/514/509/4695/4702/4698/6390/51287/1349/4722/116228/9167/4717/10063/4729 |
| hsa05010 | Alzheimer disease | 224 | -0.476760521945183 | -1.79145838035503 | 2.36971841586727e-07 | 7.39878749843004e-06 | 5.59863298251683e-06 | tags=50%, list=24%, signal=40% | 5685/5690/5714/4714/51079/1537/5684/7385/10476/506/7494/1965/5605/3265/5689/5691/55626/4707/4694/513/54539/5682/55967/9377/5692/4713/1460/1329/4700/637/5694/207/7386/5683/4711/2535/5534/203068/4708/10383/4724/292/4718/1454/5693/5719/4704/539/4716/54205/1855/4706/4726/4715/4720/801/1327/5701/572/522/84790/293/516/4893/4710/126328/10376/1340/29796/7979/4712/8883/4723/4697/1345/5688/1337/374291/517/4725/2597/27089/518/7381/7388/4709/805/10975/1350/51107/1351/1347/4696/4728/514/5715/509/4695/10381/27173/3028/4702/55851/4698/6390/7419/5686/1349/4722/9167/4717/4729 |
| hsa05022 | Pathways of neurodegeneration - multiple diseases | 261 | -0.453264495030268 | -1.7336924188567 | 4.64831708702115e-07 | 1.30617710145294e-05 | 9.88379001661339e-06 | tags=49%, list=24%, signal=39% | 7417/7345/581/5685/5690/5714/4714/51079/1537/5684/7385/6647/10476/506/7494/1965/5605/3265/5689/5691/55626/4707/4694/513/54539/5682/7332/55967/9377/5692/4713/1460/1329/4700/637/5694/7386/5683/4711/2535/5534/203068/4708/10383/4724/292/4718/1454/5693/5719/4704/539/4716/5861/54205/1855/4706/4726/4715/4720/801/5879/1327/5701/572/11315/522/84790/293/516/4893/4710/7314/2876/126328/10121/10131/10376/10540/1340/29796/7979/4712/4723/4697/1345/5688/1337/374291/517/4725/27089/518/7381/7388/4709/805/118424/10975/4218/81631/1350/1351/1347/4696/11258/4728/10280/514/5715/6233/509/4695/10381/3028/4702/7311/10671/4698/6390/7419/5686/1349/4722/9167/4717/4729 |
| hsa04932 | Non-alcoholic fatty liver disease | 95 | -0.55889014612787 | -1.91020216979094 | 2.27464953864993e-06 | 5.8106956396421e-05 | 4.39693020868217e-05 | tags=67%, list=24%, signal=52% | 3667/998/581/4714/51079/1537/7385/7494/1965/4707/4694/4296/54539/55967/9377/4713/1329/4700/637/207/7386/4711/4708/4724/4718/4704/4716/54205/4706/4726/4715/4720/5879/1327/4710/126328/1340/29796/4712/4723/4697/1345/1337/374291/4725/27089/7381/7388/4709/10975/1350/1351/1347/4696/4728/4695/4702/4698/6390/1349/4722/9167/4717/4729 |
| hsa05208 | Chemical carcinogenesis - reactive oxygen species | 145 | -0.493823238277519 | -1.78291330585936 | 8.64968503486354e-06 | 0.000202546791233055 | 0.000153266348863372 | tags=54%, list=23%, signal=43% | 51079/1537/4259/7385/6647/10476/506/5605/3265/4707/4694/513/54539/55967/9377/4713/1329/4700/207/7386/4711/4708/4724/292/4718/4704/539/4716/4706/4726/4715/4720/5879/1327/9446/572/522/293/516/4893/4710/126328/1340/29796/4712/4723/4697/1345/1337/374291/517/4725/27089/518/7381/7388/4709/10975/1350/1351/873/1347/4696/4728/514/9817/509/4695/4702/4698/6390/7419/1349/52/4722/9167/4717/4729 |
| hsa05415 | Diabetic cardiomyopathy | 137 | -0.477037768648199 | -1.70194086062892 | 1.60475253533661e-05 | 0.000346873432638143 | 0.000262477742621858 | tags=50%, list=22%, signal=40% | 7385/10476/506/5160/4707/4694/513/54539/55967/9377/4713/1329/4700/207/7386/4711/4708/4724/292/4718/5162/4704/539/4716/4706/4726/4715/4720/5879/1327/522/293/516/4710/126328/1340/29796/4712/4723/4697/1345/1337/374291/517/4725/2597/27089/518/7381/7388/4709/10975/1350/1351/1347/4696/4728/514/509/4695/4702/4698/6390/7419/1349/4722/9167/4717/4729 |
| hsa05202 | Transcriptional misregulation in cancer | 65 | 0.561925909633827 | 1.88845307600136 | 6.7941372030751e-05 | 0.00136368039576007 | 0.00103189151505351 | tags=40%, list=18%, signal=33% | 6935/2120/4790/7403/5747/6760/221037/329/8148/9611/64332/3480/4297/51274/4211/2321/4193/5327/3486/3728/5218/861/8464/55589/1655/4616 |
| hsa03040 | Spliceosome | 107 | -0.507769892627527 | -1.76198585073316 | 7.54999715179349e-05 | 0.00141436613310265 | 0.00107024521028932 | tags=48%, list=24%, signal=37% | 27316/3304/10262/9939/6626/3178/5093/25804/10189/26121/9128/6631/3303/6627/51729/642659/6428/51690/23658/27258/6634/51639/5356/4809/3312/6427/6632/10713/6628/55110/8683/25949/9416/8175/6633/6635/10465/6636/51691/22916/6637/51503/9410/10084/84844/4116/11157/10907/8896/83443/57819 |
| hsa04010 | MAPK signaling pathway | 109 | 0.467679077590276 | 1.72121699643576 | 0.000110536258302652 | 0.00194129303644032 | 0.00146896869586419 | tags=39%, list=20%, signal=32% | 9693/4790/51347/51701/6788/5922/1843/5599/4763/781/10746/6722/2247/8491/4254/10000/6197/5530/627/6789/5894/3480/4217/3727/4216/5606/22800/2260/2321/5921/56034/5578/5908/57551/5494/7424/1946/1956/4616/6654/7046/9448/1969 |
| hsa04390 | Hippo signaling pathway | 75 | 0.51419181562853 | 1.77540838971712 | 0.000129950391289367 | 0.0021480035266066 | 0.00162538569909921 | tags=37%, list=20%, signal=30% | 2932/7003/1490/6788/26524/10413/1739/23291/25937/2736/329/1495/64398/6934/4092/659/56288/5520/5054/122786/8323/7159/657/7474/4088/324/7046/83439 |
| hsa04820 | Cytoskeleton in muscle cells | 94 | 0.487556193065631 | 1.74602410356479 | 0.000183623919771739 | 0.00286657341421437 | 0.00216912466630943 | tags=38%, list=22%, signal=30% | 23345/27063/10611/1282/23353/6645/3685/1284/7414/3339/114793/2200/1289/1730/4627/56776/7168/9672/83660/3688/1462/81624/1290/481/4628/3728/29766/1832^a^/2201/7058/1281/2318/7057/1278/1729/6709 |
| hsa04928 | Parathyroid hormone synthesis, secretion and action | 46 | 0.572215121707306 | 1.8093678324556 | 0.000287249942935237 | 0.00424827547183167 | 0.00321465310099268 | tags=43%, list=20%, signal=35% | 27347/2776/4205/4325/23236/4040/11214/5144/5894/3727/9943/1839/1385/2260/5578/4323/1956/65125/6446/5142 |
| hsa04340 | Hedgehog signaling pathway | 20 | 0.736814742143444 | 1.95356311719312 | 0.000333270962475444 | 0.00468245702277999 | 0.00354319654842314 | tags=35%, list=7%, signal=33% | 2932/8452/1452/57154/23291/2736/64750 |
| hsa04723 | Retrograde endocannabinoid signaling | 64 | -0.558954357572372 | -1.80322089471281 | 0.000359760078201166 | 0.0048139324749775 | 0.00364268349857822 | tags=55%, list=20%, signal=44% | 4707/4694/54539/2783/55967/4713/4700/2773/4711/4708/4724/4718/4704/4716/4706/4726/4715/4720/4710/126328/4712/4723/4697/374291/2787/4725/4709/4696/4728/4695/4702/4698/4722/4717/4729 |
| hsa04510 | Focal adhesion | 110 | 0.440474282905621 | 1.63658326515236 | 0.000444181797241667 | 0.00567341295567766 | 0.00429304894941707 | tags=37%, list=22%, signal=30% | 2932/1793/5728/1282/5599/3685/4659/5747/6093/1284/7414/394/329/55742/10000/3371/2909/5894/3480/3915/7410/87/83660/3688/2317/2321/56034/5578/5908/7424/331/1956/81/3913/9475/6654/7058/7057/1278/53358/1729 |
| hsa04514 | Cell adhesion molecules | 38 | 0.598191873760459 | 1.83074835085834 | 0.00059720741951882 | 0.00729631673412124 | 0.00552109376397262 | tags=47%, list=22%, signal=37% | 1000/9369/22829/3685/1002/29126/9672/25945/214/3688/1462/2734/3133/257194/23705/22871/4756/80381 |
| hsa05200 | Pathways in cancer | 205 | 0.375765813345849 | 1.51207290345865 | 0.000767426307438865 | 0.00898528301626338 | 0.00679912781151977 | tags=29%, list=20%, signal=24% | 2776/2932/4790/2113/3280/5728/27436/1613/867/817/1282/5599/3685/5747/6093/1284/8202/23236/1021/1871/2736/2247/4792/4254/329/4040/25/1495/10000/1387/6934/7187/23365/6789/5894/3480/3915/3716/3688/8323/2260/4780/4193/5578/3728/861/8454/7424/7474/331/4088/1956/324/3913/4616/9475/6654/4853/7046/83439 |
| hsa04360 | Axon guidance | 79 | 0.466128330949497 | 1.61828871095082 | 0.000893025575028989 | 0.0100376074633258 | 0.00759541752235183 | tags=38%, list=20%, signal=31% | 2932/57522/817/85464/55740/5747/6093/6259/9037/8482/25/10512/659/5530/23365/56288/6091/5894/54434/2044/1948/9353/3688/5921/5578/1946/7474/9475/5362/1969 |
| hsa04668 | TNF signaling pathway | 42 | 0.572615228516827 | 1.80126085983299 | 0.00105733312555914 | 0.0114273310877738 | 0.00864701584465369 | tags=36%, list=18%, signal=30% | 4790/3726/5599/83737/10059/4792/329/10000/7187/4217/5606/1385/4323/7424/331 |
| hsa04658 | Th1 and Th2 cell differentiation | 24 | 0.656934577876372 | 1.81824918946963 | 0.00125327347563461 | 0.0130433276538269 | 0.00986983399914978 | tags=33%, list=13%, signal=29% | 4790/5599/3516/84441/4792/5530/55534/3716 |
| hsa04974 | Protein digestion and absorption | 28 | 0.615141586372829 | 1.77472126368132 | 0.00134955789674365 | 0.0135437774637487 | 0.0102485223737675 | tags=46%, list=26%, signal=35% | 84570/1282/1284/1289/1290/481/1295/54407/1281/6520/1278/1303/6546 |
| hsa03420 | Nucleotide excision repair | 31 | -0.625560699844899 | -1.75638742766355 | 0.00157720741961451 | 0.0150591102466493 | 0.0113951686451514 | tags=48%, list=18%, signal=40% | 5886/5435/3978/5111/5983/5437/9978/2067/5425/404672/5441/54107/5436/5440/5439 |
| hsa05206 | MicroRNAs in cancer | 84 | 0.46210623826939 | 1.62707644798467 | 0.00160773419003374 | 0.0150591102466493 | 0.0113951686451514 | tags=37%, list=22%, signal=29% | 2744/6935/4790/5728/4325/6093/23414/1021/1871/4363/25/1387/659/3371/5894/7168/27086/6541/4193/5578/1946/9839/1956/324/6654/57521/4853/7078/7057/6659/960 |
| hsa04520 | Adherens junction | 64 | 0.502240814068 | 1.6926502795323 | 0.00195568002023806 | 0.0175435749566335 | 0.0132751531793968 | tags=45%, list=21%, signal=36% | 5795/51701/2241/4008/6093/57154/7414/57493/64750/1500/1495/1387/6934/56288/3480/87/25945/2260/5908/10163/4088/1956/81/4301/7082/9475/7046/83439/10810 |
| hsa05224 | Breast cancer | 59 | 0.503082181309634 | 1.66730625241336 | 0.00199784483491912 | 0.0175435749566335 | 0.0132751531793968 | tags=37%, list=20%, signal=30% | 2932/1452/3280/5728/8202/1021/1871/2247/4040/10000/6934/5894/3480/8323/2260/7474/1956/324/4616/6654/4853/83439 |
| hsa04512 | ECM-receptor interaction | 29 | 0.610278912780284 | 1.77338975588063 | 0.00237691586223226 | 0.0199396265104119 | 0.0150882358310665 | tags=62%, list=30%, signal=44% | 1282/3685/1284/3339/3371/3915/3688/3913/7058/7057/1278/960/3912/375790/6385/1291/3678/3693 |
| hsa03020 | RNA polymerase | 15 | -0.724023472011505 | -1.74163465290606 | 0.00241262384823489 | 0.0199396265104119 | 0.0150882358310665 | tags=47%, list=5%, signal=44% | 5435/5437/51082/5441/5436/5440/5439 |
| hsa04660 | T cell receptor signaling pathway | 56 | 0.499721893541457 | 1.64215859756717 | 0.00249975389128819 | 0.0200694526700566 | 0.0151864747681267 | tags=21%, list=12%, signal=19% | 2932/4790/5529/5599/1739/4792/10000/5530/868/5894/5520/7410 |
| hsa04936 | Alcoholic liver disease | 44 | 0.560125854888314 | 1.76557691186551 | 0.00291532868868142 | 0.0227557600422078 | 0.0172191928395803 | tags=41%, list=24%, signal=31% | 2932/4790/5599/31/4792/10000/6934/7187/50507/4217/5606/51422/1374/83439/6416/9663/2309/6434 |
| hsa05165 | Human papillomavirus infection | 143 | 0.392250271318081 | 1.50836505364749 | 0.00306192635240561 | 0.0232540893250264 | 0.017596276619799 | tags=29%, list=20%, signal=24% | 2932/4790/1452/3280/5728/5529/9223/1282/3685/3516/5747/1739/1284/1021/84441/10000/64398/1387/6934/7187/3371/56288/5894/55534/3915/5520/3716/523/7337/1385/3688/8323/5523/4193/3133/7474/1956/324/3913/6654/4853/83439 |
| hsa04071 | Sphingolipid signaling pathway | 62 | 0.469923165769809 | 1.57048440080153 | 0.00324447870191965 | 0.0239920661905111 | 0.0181547007697443 | tags=23%, list=12%, signal=20% | 2776/4790/5728/5529/5599/6093/23236/4363/10000/5894/5520/9846/4217/9517 |
| hsa04014 | Ras signaling pathway | 87 | 0.426712756956003 | 1.52092477123094 | 0.00345829821522667 | 0.0249174820122742 | 0.0188549592301157 | tags=33%, list=18%, signal=28% | 4790/2113/9462/22821/5922/5599/4763/2247/27/4254/25/10000/627/6789/5894/3480/9846/2114/22800/2260/2321/5921/56034/5578/5908/7424/1946/1956/4301 |
| hsa05215 | Prostate cancer | 53 | 0.51286299616674 | 1.66487191019416 | 0.00389793282079378 | 0.0273829780660763 | 0.0207205902579038 | tags=36%, list=20%, signal=29% | 2932/6935/4790/5728/1871/4792/10000/1387/6934/5894/3480/1385/2260/4193/56034/5327/1956/6654/83439 |
| hsa05142 | Chagas disease | 38 | 0.538774384246488 | 1.64890290007342 | 0.00448027318998595 | 0.0307062625947817 | 0.0232353064024945 | tags=21%, list=12%, signal=19% | 2776/4790/5599/23236/4792/10000/5520/5054 |
| hsa04934 | Cushing syndrome | 56 | 0.478212337579347 | 1.57147507797448 | 0.00607685569002684 | 0.0406570583070843 | 0.0307650338191835 | tags=29%, list=20%, signal=23% | 2776/2932/817/23236/1021/3949/1871/6934/4297/1385/8323/5908/7474/1956/324/83439 |
| hsa00480 | Glutathione metabolism | 22 | -0.623469747806642 | -1.62508664360019 | 0.00678426811984855 | 0.0414430291669009 | 0.0313597748331672 | tags=73%, list=33%, signal=49% | 6241/6611/3417/2539/2730/2936/4953/9588/4259/6723/51056/9446/2950/2876/2879/3418 |
| hsa04550 | Signaling pathways regulating pluripotency of stem cells | 56 | 0.477618195263033 | 1.56952264017753 | 0.00665103542535143 | 0.0414430291669009 | 0.0313597748331672 | tags=36%, list=18%, signal=30% | 2932/5978/7994/3624/2247/10000/659/5894/3480/90/3716/6498/8323/2260/4211/657/463/7474/4088/324 |
| hsa04919 | Thyroid hormone signaling pathway | 58 | 0.467537875511849 | 1.54808192703708 | 0.00637007391431854 | 0.0414430291669009 | 0.0313597748331672 | tags=26%, list=16%, signal=22% | 2932/23389/9882/3685/8202/23236/10000/1387/9611/9969/5894/9282/4193/481/5578 |
| hsa05135 | Yersinia infection | 70 | 0.449262921248889 | 1.52605364973621 | 0.00656911494983349 | 0.0414430291669009 | 0.0313597748331672 | tags=26%, list=15%, signal=22% | 2776/2932/4790/5586/1793/5599/5747/6093/10096/4792/10000/64283/6197/23365/7410/5606/3688/10097 |
| hsa04931 | Insulin resistance | 42 | 0.515657560532728 | 1.62209060221888 | 0.00741602297169694 | 0.0443383501073796 | 0.0335506526379123 | tags=24%, list=10%, signal=21% | 2932/4790/5728/9882/5599/4792/10000/6197/2673/10724 |
| hsa04151 | PI3K-Akt signaling pathway | 130 | 0.37574540898308 | 1.42238416295995 | 0.00827265858155192 | 0.0484295221128352 | 0.0366464261726642 | tags=33%, list=22%, signal=26% | 2932/4790/5586/5728/5529/9223/1282/3685/5747/1284/1021/2247/4254/10000/9863/3371/627/5894/3480/3915/5520/3716/1385/3688/2260/5523/2321/4193/56034/5578/7424/1946/1956/3913/6654/57521/6446/1969/23239/7058/7057/64764/1278 |
