## Supplemental Table 2 for "Donor-matched iPSC model reveals context-dependent T2D genetic signals in fibro-adipogenic progenitors"

| Rank | Motif | Name | P-value | log P-pvalue | q-value (Benjamini) | # Target Sequences with Motif | % of Targets Sequences with Motif | # Background Sequences with Motif | % of Background Sequences with Motif |
| --- | --- | --- | --- | --- | --- | --- | --- | --- | --- |
| 1 | NDATGASTCAYN | Fos(bZIP)/TSC-Fos-ChIP-Seq(GSE110950)/Homer | 1e-202 | -4.666e+02 | 0.0000 | 514.0 | 21.55% | 4038.9 | 4.24% |
| 2 | DATGASTCATHN | Atf3(bZIP)/GBM-ATF3-ChIP-Seq(GSE33912)/Homer | 1e-197 | -4.553e+02 | 0.0000 | 545.0 | 22.85% | 4697.1 | 4.93% |
| 3 | DATGASTCAT | BATF(bZIP)/Th17-BATF-ChIP-Seq(GSE39756)/Homer | 1e-195 | -4.509e+02 | 0.0000 | 535.0 | 22.43% | 4561.5 | 4.79% |
| 4 | NNATGASTCATH | Fra1(bZIP)/BT549-Fra1-ChIP-Seq(GSE46166)/Homer | 1e-192 | -4.431e+02 | 0.0000 | 488.0 | 20.46% | 3814.6 | 4.00% |
| 5 | VTGACTCATC | AP-1(bZIP)/ThioMac-PU.1-ChIP-Seq(GSE21512)/Homer | 1e-180 | -4.154e+02 | 0.0000 | 568.0 | 23.82% | 5587.2 | 5.86% |
| 6 | RATGASTCAT | JunB(bZIP)/DendriticCells-Junb-ChIP-Seq(GSE36099)/Homer | 1e-175 | -4.035e+02 | 0.0000 | 473.0 | 19.83% | 3939.0 | 4.13% |
| 7 | GGATGACTCATC | Fra2(bZIP)/Striatum-Fra2-ChIP-Seq(GSE43429)/Homer | 1e-161 | -3.707e+02 | 0.0000 | 426.0 | 17.86% | 3446.2 | 3.62% |
| 8 | NATGASTCABNN | Fosl2(bZIP)/3T3L1-Fosl2-ChIP-Seq(GSE56872)/Homer | 1e-108 | -2.507e+02 | 0.0000 | 301.0 | 12.62% | 2498.4 | 2.62% |
| 9 | GATGASTCATCN | Jun-AP1(bZIP)/K562-cJun-ChIP-Seq(GSE31477)/Homer | 1e-81 | -1.872e+02 | 0.0000 | 227.0 | 9.52% | 1885.1 | 1.98% |
| 10 | TGCTGAGTCA | Bach2(bZIP)/OCILy7-Bach2-ChIP-Seq(GSE44420)/Homer | 1e-51 | -1.184e+02 | 0.0000 | 163.0 | 6.83% | 1525.6 | 1.60% |
| 11 | TRCATTCCAG | TEAD3(TEA)/HepG2-TEAD3-ChIP-Seq(Encode)/Homer | 1e-30 | -7.009e+01 | 0.0000 | 417.0 | 17.48% | 9302.4 | 9.76% |
| 12 | CYRCATTCCA | TEAD1(TEAD)/HepG2-TEAD1-ChIP-Seq(Encode)/Homer | 1e-26 | -6.129e+01 | 0.0000 | 360.0 | 15.09% | 7962.3 | 8.36% |
| 13 | YCWGGAATGY | TEAD(TEA)/Fibroblast-PU.1-ChIP-Seq(Unpublished)/Homer | 1e-25 | -5.793e+01 | 0.0000 | 261.0 | 10.94% | 5189.3 | 5.45% |
| 14 | RGCCAATSRG | NFY(CCAAT)/Promoter/Homer | 1e-23 | -5.380e+01 | 0.0000 | 296.0 | 12.41% | 6372.6 | 6.69% |
| 15 | CCWGGAATGY | TEAD4(TEA)/Tropoblast-Tead4-ChIP-Seq(GSE37350)/Homer | 1e-19 | -4.498e+01 | 0.0000 | 319.0 | 13.38% | 7490.2 | 7.86% |
| 16 | AWWNTGCTGAGTCAT | Bach1(bZIP)/K562-Bach1-ChIP-Seq(GSE31477)/Homer | 1e-16 | -3.759e+01 | 0.0000 | 45.0 | 1.89% | 377.7 | 0.40% |
| 17 | GCTGASTCAGCA | MafK(bZIP)/C2C12-MafK-ChIP-Seq(GSE36030)/Homer | 1e-16 | -3.755e+01 | 0.0000 | 121.0 | 5.07% | 2061.8 | 2.16% |
| 18 | CCWGGAATGY | TEAD2(TEA)/Py2T-Tead2-ChIP-Seq(GSE55709)/Homer | 1e-16 | -3.722e+01 | 0.0000 | 213.0 | 8.93% | 4641.6 | 4.87% |
| 19 | GCTGTGGTTW | RUNX-AML(Runt)/CD4+-PolII-ChIP-Seq(Barski_et_al.)/Homer | 1e-15 | -3.561e+01 | 0.0000 | 256.0 | 10.73% | 6025.6 | 6.32% |
| 20 | RACCGGAAGT | GABPA(ETS)/Jurkat-GABPa-ChIP-Seq(GSE17954)/Homer | 1e-15 | -3.511e+01 | 0.0000 | 328.0 | 13.75% | 8341.5 | 8.75% |
| 21 | GGAAATTCCC | NFkB-p65-Rel(RHD)/ThioMac-LPS-Expression(GSE23622)/Homer | 1e-15 | -3.491e+01 | 0.0000 | 46.0 | 1.93% | 424.9 | 0.45% |
| 22 | SAAACCACAG | RUNX(Runt)/HPC7-Runx1-ChIP-Seq(GSE22178)/Homer | 1e-14 | -3.415e+01 | 0.0000 | 246.0 | 10.31% | 5793.2 | 6.08% |
| 23 | GATGACTCAGCA | NF-E2(bZIP)/K562-NFE2-ChIP-Seq(GSE31477)/Homer | 1e-14 | -3.384e+01 | 0.0000 | 45.0 | 1.89% | 419.2 | 0.44% |
| 24 | NRYTTCCGGH | Fli1(ETS)/CD8-FLI-ChIP-Seq(GSE20898)/Homer | 1e-13 | -3.209e+01 | 0.0000 | 387.0 | 16.23% | 10499.4 | 11.02% |
| 25 | ACCGGAAGTG | ETV4(ETS)/HepG2-ETV4-ChIP-Seq(ENCODE)/Homer | 1e-13 | -3.183e+01 | 0.0000 | 416.0 | 17.44% | 11501.0 | 12.07% |
| 26 | NNAYTTCCTGHN | Etv2(ETS)/ES-ER71-ChIP-Seq(GSE59402)/Homer | 1e-13 | -3.077e+01 | 0.0000 | 323.0 | 13.54% | 8462.5 | 8.88% |
| 27 | NWAACCACADNN | RUNX2(Runt)/PCa-RUNX2-ChIP-Seq(GSE33889)/Homer | 1e-13 | -3.050e+01 | 0.0000 | 268.0 | 11.24% | 6694.2 | 7.03% |
| 28 | AAACCACARM | RUNX1(Runt)/Jurkat-RUNX1-ChIP-Seq(GSE29180)/Homer | 1e-13 | -3.045e+01 | 0.0000 | 315.0 | 13.21% | 8221.6 | 8.63% |
| 29 | GGTTGCCATGGCAA | X-box(HTH)/NPC-H3K4me1-ChIP-Seq(GSE16256)/Homer | 1e-12 | -2.967e+01 | 0.0000 | 63.0 | 2.64% | 849.9 | 0.89% |
| 30 | NRYTTCCGGY | Elk4(ETS)/Hela-Elk4-ChIP-Seq(GSE31477)/Homer | 1e-12 | -2.907e+01 | 0.0000 | 219.0 | 9.18% | 5230.8 | 5.49% |
| 31^a^ | TWGTCTGV | Smad3(MAD)/NPC-Smad3-ChIP-Seq(GSE36673)/Homer | 1e-12 | -2.841e+01 | 0.0000 | 863.0 | 36.18% | 27985.2 | 29.37% |
| 32 | ACTTCCKGKT | Elf4(ETS)/BMDM-Elf4-ChIP-Seq(GSE88699)/Homer | 1e-12 | -2.838e+01 | 0.0000 | 338.0 | 14.17% | 9130.4 | 9.58% |
| 33 | WGGGGATTTCCC | NFkB-p65(RHD)/GM12787-p65-ChIP-Seq(GSE19485)/Homer | 1e-12 | -2.822e+01 | 0.0000 | 213.0 | 8.93% | 5090.2 | 5.34% |
| 34 | TGCTGACTCA | MafA(bZIP)/Islet-MafA-ChIP-Seq(GSE30298)/Homer | 1e-12 | -2.769e+01 | 0.0000 | 275.0 | 11.53% | 7099.9 | 7.45% |
| 35 | ACAGGAAGTG | ETS1(ETS)/Jurkat-ETS1-ChIP-Seq(GSE17954)/Homer | 1e-11 | -2.747e+01 | 0.0000 | 358.0 | 15.01% | 9873.6 | 10.36% |
| 36 | ATTTCCTGTN | EWS:ERG-fusion(ETS)/CADO_ES1-EWS:ERG-ChIP-Seq(SRA014231)/Homer | 1e-11 | -2.678e+01 | 0.0000 | 222.0 | 9.31% | 5455.7 | 5.73% |
| 37 | HACTTCCGGY | Elk1(ETS)/Hela-Elk1-ChIP-Seq(GSE31477)/Homer | 1e-11 | -2.631e+01 | 0.0000 | 219.0 | 9.18% | 5388.8 | 5.66% |
| 38 | GGCCCCGCCCCC | Sp1(Zf)/Promoter/Homer | 1e-11 | -2.588e+01 | 0.0000 | 174.0 | 7.30% | 4014.9 | 4.21% |
| 39 | AACCGGAAGT | ETV1(ETS)/GIST48-ETV1-ChIP-Seq(GSE22441)/Homer | 1e-11 | -2.546e+01 | 0.0000 | 449.0 | 18.83% | 13186.6 | 13.84% |
| 40 | VACAGGAAAT | EWS:FLI1-fusion(ETS)/SK_N_MC-EWS:FLI1-ChIP-Seq(SRA014231)/Homer | 1e-11 | -2.541e+01 | 0.0000 | 206.0 | 8.64% | 5031.2 | 5.28% |
| 41 | RGKGGGCGGAGC | Sp5(Zf)/mES-Sp5.Flag-ChIP-Seq(GSE72989)/Homer | 1e-10 | -2.452e+01 | 0.0000 | 491.0 | 20.59% | 14762.1 | 15.49% |
| 42 | AVCCGGAAGT | ELF1(ETS)/Jurkat-ELF1-ChIP-Seq(SRA014231)/Homer | 1e-10 | -2.415e+01 | 0.0000 | 203.0 | 8.51% | 5011.4 | 5.26% |
| 43 | GTTGCCATGGCAACM | Rfx2(HTH)/LoVo-RFX2-ChIP-Seq(GSE49402)/Homer | 1e-10 | -2.384e+01 | 0.0000 | 55.0 | 2.31% | 789.9 | 0.83% |
| 44 | GCYCCGCCCCYH | KLF17(Zf)/2cell-Klf17-CutnTag(GSE211845)/Homer | 1e-10 | -2.368e+01 | 0.0000 | 280.0 | 11.74% | 7546.7 | 7.92% |
| 45 | AACCGGAAGT | ETS(ETS)/Promoter/Homer | 1e-10 | -2.345e+01 | 0.0000 | 138.0 | 5.79% | 3051.6 | 3.20% |
| 46 | CGGTTGCCATGGCAAC | RFX(HTH)/K562-RFX3-ChIP-Seq(SRA012198)/Homer | 1e-10 | -2.334e+01 | 0.0000 | 51.0 | 2.14% | 708.3 | 0.74% |
| 47 | YCATYAATCA | Pdx1(Homeobox)/Islet-Pdx1-ChIP-Seq(SRA008281)/Homer | 1e-9 | -2.256e+01 | 0.0000 | 232.0 | 9.73% | 6045.4 | 6.34% |
| 48 | ACAGGAAGTG | ERG(ETS)/VCaP-ERG-ChIP-Seq(GSE14097)/Homer | 1e-9 | -2.237e+01 | 0.0000 | 501.0 | 21.01% | 15343.5 | 16.10% |
| 49 | SCCTAGCAACAG | Rfx5(HTH)/GM12878-Rfx5-ChIP-Seq(GSE31477)/Homer | 1e-9 | -2.197e+01 | 0.0000 | 119.0 | 4.99% | 2559.1 | 2.69% |
| 50 | AWWWTGCTGAGTCAT | NFE2L2(bZIP)/HepG2-NFE2L2-ChIP-Seq(Encode)/Homer | 1e-9 | -2.151e+01 | 0.0000 | 30.0 | 1.26% | 300.9 | 0.32% |
| 51 | KGTTGCCATGGCAA | Rfx1(HTH)/NPC-H3K4me1-ChIP-Seq(GSE16256)/Homer | 1e-9 | -2.115e+01 | 0.0000 | 91.0 | 3.82% | 1794.8 | 1.88% |
| 52 | MACAGGAAGT | Ets1-distal(ETS)/CD4+-PolII-ChIP-Seq(Barski_et_al.)/Homer | 1e-9 | -2.080e+01 | 0.0000 | 113.0 | 4.74% | 2436.0 | 2.56% |
| 53 | ASWTCCTGBT | SPDEF(ETS)/VCaP-SPDEF-ChIP-Seq(SRA014231)/Homer | 1e-8 | -2.063e+01 | 0.0000 | 323.0 | 13.54% | 9251.5 | 9.71% |
| 54 | RGKGGGCGKGGC | KLF14(Zf)/HEK293-KLF14.GFP-ChIP-Seq(GSE58341)/Homer | 1e-8 | -1.863e+01 | 0.0000 | 786.0 | 32.96% | 26365.1 | 27.67% |
| 55 | HTGCTGAGTCAT | Nrf2(bZIP)/Lymphoblast-Nrf2-ChIP-Seq(GSE37589)/Homer | 1e-7 | -1.823e+01 | 0.0000 | 31.0 | 1.30% | 368.4 | 0.39% |
| 56 | TGTTTAYTTAGC | FoxD3(forkhead)/ZebrafishEmbryo-Foxd3.biotin-ChIP-seq(GSE106676)/Homer | 1e-7 | -1.804e+01 | 0.0000 | 172.0 | 7.21% | 4408.1 | 4.63% |
| 57 | DGGGYGKGGC | KLF5(Zf)/LoVo-KLF5-ChIP-Seq(GSE49402)/Homer | 1e-7 | -1.752e+01 | 0.0000 | 602.0 | 25.24% | 19621.8 | 20.59% |
| 58 | AVCAGGAAGT | EHF(ETS)/LoVo-EHF-ChIP-Seq(GSE49402)/Homer | 1e-7 | -1.749e+01 | 0.0000 | 361.0 | 15.14% | 10880.7 | 11.42% |
| 59 | NGCGTGGGCGGR | Egr2(Zf)/Thymocytes-Egr2-ChIP-Seq(GSE34254)/Homer | 1e-7 | -1.702e+01 | 0.0000 | 95.0 | 3.98% | 2080.2 | 2.18% |
| 60 | RTGATTKATRGN | PBX2(Homeobox)/K562-PBX2-ChIP-Seq(Encode)/Homer | 1e-6 | -1.612e+01 | 0.0000 | 184.0 | 7.71% | 4935.7 | 5.18% |
| 61 | AGAGGAAGTG | PU.1(ETS)/ThioMac-PU.1-ChIP-Seq(GSE21512)/Homer | 1e-6 | -1.605e+01 | 0.0000 | 160.0 | 6.71% | 4156.7 | 4.36% |
| 62 | WAAGTAAACA | FOXA1(Forkhead)/MCF7-FOXA1-ChIP-Seq(GSE26831)/Homer | 1e-6 | -1.595e+01 | 0.0000 | 186.0 | 7.80% | 5014.6 | 5.26% |
| 63 | ANCAGGAAGT | ELF3(ETS)/PDAC-ELF3-ChIP-Seq(GSE64557)/Homer | 1e-6 | -1.548e+01 | 0.0000 | 211.0 | 8.85% | 5886.3 | 6.18% |
| 64 | WAAGTAAACA | FOXA1(Forkhead)/LNCAP-FOXA1-ChIP-Seq(GSE27824)/Homer | 1e-6 | -1.539e+01 | 0.0000 | 224.0 | 9.39% | 6333.3 | 6.65% |
| 65 | VDGGGYGGGGCY | KLF1(Zf)/HUDEP2-KLF1-CutnRun(GSE136251)/Homer | 1e-6 | -1.527e+01 | 0.0000 | 438.0 | 18.36% | 13898.0 | 14.59% |
| 66 | VBSYGTCTGG | Smad4(MAD)/ESC-SMAD4-ChIP-Seq(GSE29422)/Homer | 1e-6 | -1.527e+01 | 0.0000 | 504.0 | 21.13% | 16310.3 | 17.12% |
| 67 | GCTATTTTTAGC | Mef2d(MADS)/Retina-Mef2d-ChIP-Seq(GSE61391)/Homer | 1e-6 | -1.522e+01 | 0.0000 | 59.0 | 2.47% | 1129.4 | 1.19% |
| 68 | CCATATATGGNM | CArG(MADS)/PUER-Srf-ChIP-Seq(Sullivan_et_al.)/Homer | 1e-6 | -1.516e+01 | 0.0000 | 85.0 | 3.56% | 1872.3 | 1.97% |
| 69 | GCCACRCCCACY | Klf9(Zf)/GBM-Klf9-ChIP-Seq(GSE62211)/Homer | 1e-6 | -1.492e+01 | 0.0000 | 191.0 | 8.01% | 5262.0 | 5.52% |
| 70 | TGCGTGGGYG | Egr1(Zf)/K562-Egr1-ChIP-Seq(GSE32465)/Homer | 1e-6 | -1.446e+01 | 0.0000 | 289.0 | 12.12% | 8667.9 | 9.10% |
| 71 | NRGCCCCRCCCHBNN | KLF3(Zf)/MEF-Klf3-ChIP-Seq(GSE44748)/Homer | 1e-6 | -1.439e+01 | 0.0000 | 256.0 | 10.73% | 7524.2 | 7.90% |
| 72 | DGTAAACA | Foxo3(Forkhead)/U2OS-Foxo3-ChIP-Seq(E-MTAB-2701)/Homer | 1e-6 | -1.405e+01 | 0.0000 | 144.0 | 6.04% | 3777.9 | 3.97% |
| 73 | YGGCCCCGCCCC | Sp2(Zf)/HEK293-Sp2.eGFP-ChIP-Seq(Encode)/Homer | 1e-6 | -1.399e+01 | 0.0000 | 674.0 | 28.26% | 22845.0 | 23.98% |
| 74 | ACVAGGAAGT | ELF5(ETS)/T47D-ELF5-ChIP-Seq(GSE30407)/Homer | 1e-6 | -1.384e+01 | 0.0000 | 208.0 | 8.72% | 5927.4 | 6.22% |
| 75 | NYYTGTTTACHN | FOXP1(Forkhead)/H9-FOXP1-ChIP-Seq(GSE31006)/Homer | 1e-5 | -1.346e+01 | 0.0000 | 87.0 | 3.65% | 2016.9 | 2.12% |
| 76 | CTGTCTGG | Smad2(MAD)/ES-SMAD2-ChIP-Seq(GSE29422)/Homer | 1e-5 | -1.318e+01 | 0.0000 | 483.0 | 20.25% | 15832.1 | 16.62% |
| 77 | WWATRTAAACAN | Foxf1(Forkhead)/Lung-Foxf1-ChIP-Seq(GSE77951)/Homer | 1e-5 | -1.301e+01 | 0.0000 | 167.0 | 7.00% | 4615.7 | 4.84% |
| 78 | RGGGGGCGGGGC | Klf15(Zf)/Liver-Klf15-ChIP-Seq(GSE166083)/Homer | 1e-5 | -1.295e+01 | 0.0000 | 493.0 | 20.67% | 16236.5 | 17.04% |
| 79 | WWTRTAAACAVG | FoxL2(Forkhead)/Ovary-FoxL2-ChIP-Seq(GSE60858)/Homer | 1e-5 | -1.285e+01 | 0.0000 | 153.0 | 6.42% | 4162.9 | 4.37% |
| 80 | ATGCATWATGCATRW | OCT:OCT-short(POU,Homeobox)/NPC-OCT6-ChIP-Seq(GSE43916)/Homer | 1e-5 | -1.283e+01 | 0.0000 | 152.0 | 6.37% | 4130.0 | 4.33% |
| 81 | MCTCCCMCRCAB | WT1(Zf)/Kidney-WT1-ChIP-Seq(GSE90016)/Homer | 1e-5 | -1.208e+01 | 0.0000 | 252.0 | 10.57% | 7625.2 | 8.00% |
| 82 | RTCATGTGAC | MITF(bHLH)/MastCells-MITF-ChIP-Seq(GSE48085)/Homer | 1e-5 | -1.189e+01 | 0.0000 | 282.0 | 11.82% | 8708.0 | 9.14% |
| 83 | NVWTGTTTAC | FOXK1(Forkhead)/HEK293-FOXK1-ChIP-Seq(GSE51673)/Homer | 1e-5 | -1.178e+01 | 0.0000 | 199.0 | 8.34% | 5812.1 | 6.10% |
| 84 | MKGGGYGTGGCC | KLF6(Zf)/PDAC-KLF6-ChIP-Seq(GSE64557)/Homer | 1e-5 | -1.175e+01 | 0.0000 | 469.0 | 19.66% | 15529.6 | 16.30% |
| 85 | WNTGCTGASTCAGCANWTTY | MafB(bZIP)/BMM-Mafb-ChIP-Seq(GSE75722)/Homer | 1e-4 | -1.145e+01 | 0.0001 | 123.0 | 5.16% | 3281.7 | 3.44% |
| 86 | BSNTGTTTACWYWGN | Foxa3(Forkhead)/Liver-Foxa3-ChIP-Seq(GSE77670)/Homer | 1e-4 | -1.082e+01 | 0.0001 | 63.0 | 2.64% | 1427.9 | 1.50% |
| 87 | ACCRTGACTAATTNN | PAX3:FKHR-fusion(Paired,Homeobox)/Rh4-PAX3:FKHR-ChIP-Seq(GSE19063)/Homer | 1e-4 | -1.079e+01 | 0.0001 | 51.0 | 2.14% | 1076.1 | 1.13% |
| 88 | SCHTGTTTACAT | FOXK2(Forkhead)/U2OS-FOXK2-ChIP-Seq(E-MTAB-2204)/Homer | 1e-4 | -1.024e+01 | 0.0002 | 136.0 | 5.70% | 3809.2 | 4.00% |
| 89 | TRTTTACTTW | FOXM1(Forkhead)/MCF7-FOXM1-ChIP-Seq(GSE72977)/Homer | 1e-4 | -1.020e+01 | 0.0002 | 185.0 | 7.76% | 5487.3 | 5.76% |
| 90 | WNTGTTTRYTTTGGCA | NF1:FOXA1(CTF,Forkhead)/LNCAP-FOXA1-ChIP-Seq(GSE27824)/Homer | 1e-4 | -1.016e+01 | 0.0002 | 18.0 | 0.75% | 233.0 | 0.24% |
| 91 | NNNVCTGWGYAAACASN | Fox:Ebox(Forkhead,bHLH)/Panc1-Foxa2-ChIP-Seq(GSE47459)/Homer | 1e-4 | -1.009e+01 | 0.0002 | 214.0 | 8.97% | 6512.4 | 6.83% |
| 92 | WTTTTCCATTGS | NFATC2(RHD)/Islets-NFATC2-ChIP-Seq(GSE158496)/Homer | 1e-3 | -8.630e+00 | 0.0009 | 413.0 | 17.32% | 13963.8 | 14.66% |
| 93 | RCAGGATGTGGT | ETS:RUNX(ETS,Runt)/Jurkat-RUNX1-ChIP-Seq(GSE17954)/Homer | 1e-3 | -8.543e+00 | 0.0010 | 47.0 | 1.97% | 1061.7 | 1.11% |
| 94 | HTTTCCCASG | Rbpj1(?)/Panc1-Rbpj1-ChIP-Seq(GSE47459)/Homer | 1e-3 | -8.432e+00 | 0.0011 | 409.0 | 17.15% | 13849.5 | 14.54% |
| 95 | NDCTAATTAS | En1(Homeobox)/SUM149-EN1-ChIP-Seq(GSE120957)/Homer | 1e-3 | -8.350e+00 | 0.0012 | 349.0 | 14.63% | 11635.2 | 12.21% |
| 96 | SARTGGAAAAWRTGAGTCAB | NFAT:AP1(RHD,bZIP)/Jurkat-NFATC1-ChIP-Seq(Jolma_et_al.)/Homer | 1e-3 | -7.957e+00 | 0.0017 | 44.0 | 1.84% | 1000.9 | 1.05% |
| 97 | CYTGTTTACWYW | Foxa2(Forkhead)/Liver-Foxa2-ChIP-Seq(GSE25694)/Homer | 1e-3 | -7.871e+00 | 0.0019 | 145.0 | 6.08% | 4343.8 | 4.56% |
| 98 | GGGGGAATCCCC | NFkB-p50,p52(RHD)/Monocyte-p50-ChIP-Chip(Schreiber_et_al.)/Homer | 1e-3 | -7.863e+00 | 0.0019 | 58.0 | 2.43% | 1435.6 | 1.51% |
| 99 | NAHCAGCTGD | Ap4(bHLH)/AML-Tfap4-ChIP-Seq(GSE45738)/Homer | 1e-3 | -7.782e+00 | 0.0020 | 416.0 | 17.44% | 14230.6 | 14.94% |
| 100 | RGCAATNAAA | Hoxa9(Homeobox)/ChickenMSG-Hoxa9.Flag-ChIP-Seq(GSE86088)/Homer | 1e-3 | -7.440e+00 | 0.0028 | 545.0 | 22.85% | 19178.9 | 20.13% |
| 101 | ATGCATAATTCA | Pit1+1bp(Homeobox)/GCrat-Pit1-ChIP-Seq(GSE58009)/Homer | 1e-3 | -7.358e+00 | 0.0030 | 59.0 | 2.47% | 1498.5 | 1.57% |
| 102 | AYAGTGCCMYCTRGTGGCCA | CTCF(Zf)/CD4+-CTCF-ChIP-Seq(Barski_et_al.)/Homer | 1e-3 | -7.309e+00 | 0.0031 | 47.0 | 1.97% | 1125.7 | 1.18% |
| 103 | CTGTTTAC | Foxo1(Forkhead)/RAW-Foxo1-ChIP-Seq(Fan_et_al.)/Homer | 1e-3 | -7.262e+00 | 0.0032 | 417.0 | 17.48% | 14368.9 | 15.08% |
| 104 | CACAGCAGGGGG | Unknown-ESC-element(?)/mES-Nanog-ChIP-Seq(GSE11724)/Homer | 1e-3 | -7.236e+00 | 0.0033 | 234.0 | 9.81% | 7595.9 | 7.97% |
| 105 | VVATGACGTCAT | CREB5(bZIP)/LNCaP-CREB5.V5-ChIP-Seq(GSE137775)/Homer | 1e-3 | -7.232e+00 | 0.0033 | 108.0 | 4.53% | 3132.3 | 3.29% |
| 106 | ATGACGTCATCN | JunD(bZIP)/K562-JunD-ChIP-Seq/Homer | 1e-3 | -6.970e+00 | 0.0042 | 31.0 | 1.30% | 662.3 | 0.70% |
| 107 | TGATTRATGGCY | Hoxb4(Homeobox)/ES-Hoxb4-ChIP-Seq(GSE34014)/Homer | 1e-3 | -6.955e+00 | 0.0042 | 47.0 | 1.97% | 1145.3 | 1.20% |
| 108 | CNNBRGCGCCCCCTGSTGGC | BORIS(Zf)/K562-CTCFL-ChIP-Seq(GSE32465)/Homer | 1e-2 | -6.676e+00 | 0.0055 | 84.0 | 3.52% | 2369.8 | 2.49% |
| 109 | TGTTKCCTAGCAACM | Rfx6(HTH)/Min6b1-Rfx6.HA-ChIP-Seq(GSE62844)/Homer | 1e-2 | -6.671e+00 | 0.0055 | 303.0 | 12.70% | 10221.1 | 10.73% |
| 110 | GGCMATGAAA | Hoxd10(Homeobox)/ChickenMSG-Hoxd10.Flag-ChIP-Seq(GSE86088)/Homer | 1e-2 | -6.637e+00 | 0.0056 | 223.0 | 9.35% | 7285.7 | 7.65% |
| 111 | YCTTTGTTCC | Sox4(HMG)/proB-Sox4-ChIP-Seq(GSE50066)/Homer | 1e-2 | -6.604e+00 | 0.0058 | 175.0 | 7.34% | 5555.9 | 5.83% |
| 112 | GGGGGTGTGTCC | KLF10(Zf)/HEK293-KLF10.GFP-ChIP-Seq(GSE58341)/Homer | 1e-2 | -6.524e+00 | 0.0062 | 215.0 | 9.01% | 7011.8 | 7.36% |
| 113 | GGGGGGGG | Maz(Zf)/HepG2-Maz-ChIP-Seq(GSE31477)/Homer | 1e-2 | -6.489e+00 | 0.0063 | 533.0 | 22.35% | 18936.5 | 19.87% |
| 114 | ATGACGTCATCY | c-Jun-CRE(bZIP)/K562-cJun-ChIP-Seq(GSE31477)/Homer | 1e-2 | -6.404e+00 | 0.0068 | 109.0 | 4.57% | 3249.2 | 3.41% |
| 115 | NGYCATAAAWCH | CDX4(Homeobox)/ZebrafishEmbryos-Cdx4.Myc-ChIP-Seq(GSE48254)/Homer | 1e-2 | -6.268e+00 | 0.0078 | 150.0 | 6.29% | 4706.5 | 4.94% |
| 116 | TTTTATKRGG | HOXB13(Homeobox)/ProstateTumor-HOXB13-ChIP-Seq(GSE56288)/Homer | 1e-2 | -6.261e+00 | 0.0078 | 181.0 | 7.59% | 5818.4 | 6.11% |
| 117 | RGCMGCCTYC | ZNF91(Zf)/HEK-ZNF91.HA-ChIP-Seq(GSE162571)/Homer | 1e-2 | -6.221e+00 | 0.0080 | 332.0 | 13.92% | 11385.6 | 11.95% |
| 118 | GGVTCTCGCGAGAAC | ZBTB33(Zf)/GM12878-ZBTB33-ChIP-Seq(GSE32465)/Homer | 1e-2 | -6.123e+00 | 0.0088 | 18.0 | 0.75% | 331.0 | 0.35% |
| 119 | GGMGCTGTCCATGGTGCTGA | REST-NRSF(Zf)/Jurkat-NRSF-ChIP-Seq/Homer | 1e-2 | -6.021e+00 | 0.0096 | 8.0 | 0.34% | 91.7 | 0.10% |
| 120 | AVCAGCTG | SCL(bHLH)/HPC7-Scl-ChIP-Seq(GSE13511)/Homer | 1e-2 | -6.008e+00 | 0.0097 | 1282.0 | 53.75% | 48455.4 | 50.85% |
| 121 | TGGAACAGMA | ZNF189(Zf)/HEK293-ZNF189.GFP-ChIP-Seq(GSE58341)/Homer | 1e-2 | -5.866e+00 | 0.0111 | 218.0 | 9.14% | 7224.5 | 7.58% |
| 122 | NCYAATAAAA | Hoxd13(Homeobox)/ChickenMSG-Hoxd13.Flag-ChIP-Seq(GSE86088)/Homer | 1e-2 | -5.785e+00 | 0.0119 | 281.0 | 11.78% | 9566.2 | 10.04% |
| 123 | CMATAAATCA | Hoxc10(Homeobox)/EB-Hoxc10.iFlag-ChIP-Seq(GSE142377)/Homer | 1e-2 | -5.732e+00 | 0.0124 | 200.0 | 8.39% | 6587.3 | 6.91% |
| 124 | TGGTACATTCCA | PRDM10(Zf)/HEK293-PRDM10.eGFP-ChIP-Seq(Encode)/Homer | 1e-2 | -5.704e+00 | 0.0127 | 214.0 | 8.97% | 7104.3 | 7.46% |
| 125 | CVGTSCTCCC | Znf263(Zf)/K562-Znf263-ChIP-Seq(GSE31477)/Homer | 1e-2 | -5.661e+00 | 0.0131 | 665.0 | 27.88% | 24234.7 | 25.43% |
| 126 | VVRRAACAATGG | Sox7(HMG)/ESC-Sox7-ChIP-Seq(GSE133899)/Homer | 1e-2 | -5.616e+00 | 0.0136 | 67.0 | 2.81% | 1889.5 | 1.98% |
| 127 | NRRTGACGTCAT | Atf2(bZIP)/3T3L1-Atf2-ChIP-Seq(GSE56872)/Homer | 1e-2 | -5.560e+00 | 0.0143 | 111.0 | 4.65% | 3412.5 | 3.58% |
| 128 | GGCCYCCTGCTGDGH | Zic3(Zf)/mES-Zic3-ChIP-Seq(GSE37889)/Homer | 1e-2 | -5.463e+00 | 0.0156 | 257.0 | 10.78% | 8734.2 | 9.17% |
| 129 | ATGATTGATGGS | HOXA3(Homeobox)/mEmbryo-Hoxa3-ChIP-Seq(E-MTAB-8607)/Homer | 1e-2 | -5.376e+00 | 0.0169 | 37.0 | 1.55% | 925.9 | 0.97% |
| 130 | NVCTAATTAG | Emx2(Homeobox)/Cortex-Emx2-ChIP-Seq(GSE183130)/Homer | 1e-2 | -5.312e+00 | 0.0179 | 225.0 | 9.43% | 7576.7 | 7.95% |
| 131 | CHCAGCRGGRGG | Zic2(Zf)/ESC-Zic2-ChIP-Seq(SRP197560)/Homer | 1e-2 | -5.249e+00 | 0.0189 | 196.0 | 8.22% | 6519.1 | 6.84% |
| 132 | ATGMATATDC | Pit1(Homeobox)/GCrat-Pit1-ChIP-Seq(GSE58009)/Homer | 1e-2 | -5.167e+00 | 0.0204 | 173.0 | 7.25% | 5691.1 | 5.97% |
| 133 | TTTTATKRGN | Hoxc13(Homeobox)/EB-Hoxc13.HA-ChIP-Seq(GSE142377)/Homer | 1e-2 | -5.099e+00 | 0.0217 | 235.0 | 9.85% | 7985.8 | 8.38% |
| 134 | GYMATAAAAH | Cdx2(Homeobox)/mES-Cdx2-ChIP-Seq(GSE14586)/Homer | 1e-2 | -4.943e+00 | 0.0251 | 111.0 | 4.65% | 3487.4 | 3.66% |
| 135 | ATTGCATCAT | Chop(bZIP)/MEF-Chop-ChIP-Seq(GSE35681)/Homer | 1e-2 | -4.854e+00 | 0.0273 | 57.0 | 2.39% | 1619.0 | 1.70% |
| 136 | NNTGTGGATTSS | Foxh1(Forkhead)/hESC-FOXH1-ChIP-Seq(GSE29422)/Homer | 1e-2 | -4.849e+00 | 0.0273 | 123.0 | 5.16% | 3928.2 | 4.12% |
| 137 | GGCVGTTR | MYB(HTH)/ERMYB-Myb-ChIPSeq(GSE22095)/Homer | 1e-2 | -4.715e+00 | 0.0309 | 364.0 | 15.26% | 12919.8 | 13.56% |
| 138 | DRTGTTGCAA | CEBP:AP1(bZIP)/ThioMac-CEBPb-ChIP-Seq(GSE21512)/Homer | 1e-2 | -4.696e+00 | 0.0312 | 151.0 | 6.33% | 4963.3 | 5.21% |
| 139 | CCWTTGTY | Sox3(HMG)/NPC-Sox3-ChIP-Seq(GSE33059)/Homer | 1e-2 | -4.654e+00 | 0.0323 | 326.0 | 13.67% | 11492.5 | 12.06% |
