## Supplemental Table 3 for "Donor-matched iPSC model reveals context-dependent T2D genetic signals in fibro-adipogenic progenitors"

| Rank | Motif | Name | P-value | log P-pvalue | q-value (Benjamini) | # Target Sequences with Motif | % of Targets Sequences with Motif | # Background Sequences with Motif | % of Background Sequences with Motif |
| --- | --- | --- | --- | --- | --- | --- | --- | --- | --- |
| 1 | NDATGASTCAYN | Fos(bZIP)/TSC-Fos-ChIP-Seq(GSE110950)/Homer | 1e-320 | -7.370e+02 | 0.0000 | 552.0 | 28.19% | 3426.6 | 3.52% |
| 2 | NNATGASTCATH | Fra1(bZIP)/BT549-Fra1-ChIP-Seq(GSE46166)/Homer | 1e-315 | -7.266e+02 | 0.0000 | 535.0 | 27.32% | 3223.6 | 3.31% |
| 3 | GGATGACTCATC | Fra2(bZIP)/Striatum-Fra2-ChIP-Seq(GSE43429)/Homer | 1e-311 | -7.168e+02 | 0.0000 | 506.0 | 25.84% | 2837.5 | 2.92% |
| 4 | DATGASTCATHN | Atf3(bZIP)/GBM-ATF3-ChIP-Seq(GSE33912)/Homer | 1e-301 | -6.950e+02 | 0.0000 | 567.0 | 28.96% | 3989.6 | 4.10% |
| 5 | NATGASTCABNN | Fosl2(bZIP)/3T3L1-Fosl2-ChIP-Seq(GSE56872)/Homer | 1e-299 | -6.894e+02 | 0.0000 | 431.0 | 22.01% | 1943.2 | 2.00% |
| 6 | RATGASTCAT | JunB(bZIP)/DendriticCells-Junb-ChIP-Seq(GSE36099)/Homer | 1e-296 | -6.817e+02 | 0.0000 | 521.0 | 26.61% | 3303.2 | 3.39% |
| 7 | GATGASTCATCN | Jun-AP1(bZIP)/K562-cJun-ChIP-Seq(GSE31477)/Homer | 1e-282 | -6.511e+02 | 0.0000 | 373.0 | 19.05% | 1417.7 | 1.46% |
| 8 | DATGASTCAT | BATF(bZIP)/Th17-BATF-ChIP-Seq(GSE39756)/Homer | 1e-282 | -6.498e+02 | 0.0000 | 543.0 | 27.73% | 3922.3 | 4.03% |
| 9 | VTGACTCATC | AP-1(bZIP)/ThioMac-PU.1-ChIP-Seq(GSE21512)/Homer | 1e-277 | -6.379e+02 | 0.0000 | 575.0 | 29.37% | 4622.9 | 4.75% |
| 10 | TGCTGAGTCA | Bach2(bZIP)/OCILy7-Bach2-ChIP-Seq(GSE44420)/Homer | 1e-130 | -3.003e+02 | 0.0000 | 221.0 | 11.29% | 1237.9 | 1.27% |
| 11 | AYAGTGCCMYCTRGTGGCCA | CTCF(Zf)/CD4+-CTCF-ChIP-Seq(Barski_et_al.)/Homer | 1e-81 | -1.888e+02 | 0.0000 | 165.0 | 8.43% | 1167.3 | 1.20% |
| 12 | AWWNTGCTGAGTCAT | Bach1(bZIP)/K562-Bach1-ChIP-Seq(GSE31477)/Homer | 1e-81 | -1.885e+02 | 0.0000 | 95.0 | 4.85% | 267.8 | 0.28% |
| 13 | AWWWTGCTGAGTCAT | NFE2L2(bZIP)/HepG2-NFE2L2-ChIP-Seq(Encode)/Homer | 1e-66 | -1.536e+02 | 0.0000 | 78.0 | 3.98% | 223.2 | 0.23% |
| 14 | GATGACTCAGCA | NF-E2(bZIP)/K562-NFE2-ChIP-Seq(GSE31477)/Homer | 1e-65 | -1.502e+02 | 0.0000 | 87.0 | 4.44% | 322.6 | 0.33% |
| 15 | HTGCTGAGTCAT | Nrf2(bZIP)/Lymphoblast-Nrf2-ChIP-Seq(GSE37589)/Homer | 1e-60 | -1.394e+02 | 0.0000 | 79.0 | 4.03% | 281.8 | 0.29% |
| 16 | CNNBRGCGCCCCCTGSTGGC | BORIS(Zf)/K562-CTCFL-ChIP-Seq(GSE32465)/Homer | 1e-50 | -1.160e+02 | 0.0000 | 218.0 | 11.13% | 3323.2 | 3.41% |
| 17 | GCTGASTCAGCA | MafK(bZIP)/C2C12-MafK-ChIP-Seq(GSE36030)/Homer | 1e-49 | -1.129e+02 | 0.0000 | 149.0 | 7.61% | 1674.5 | 1.72% |
| 18 | GGCCCCGCCCCC | Sp1(Zf)/Promoter/Homer | 1e-43 | -1.012e+02 | 0.0000 | 356.0 | 18.18% | 8023.8 | 8.24% |
| 19 | GCYCCGCCCCYH | KLF17(Zf)/2cell-Klf17-CutnTag(GSE211845)/Homer | 1e-36 | -8.315e+01 | 0.0000 | 468.0 | 23.90% | 12946.9 | 13.30% |
| 20 | NRGCCCCRCCCHBNN | KLF3(Zf)/MEF-Klf3-ChIP-Seq(GSE44748)/Homer | 1e-35 | -8.204e+01 | 0.0000 | 378.0 | 19.31% | 9606.4 | 9.87% |
| 21 | RGKGGGCGGAGC | Sp5(Zf)/mES-Sp5.Flag-ChIP-Seq(GSE72989)/Homer | 1e-35 | -8.092e+01 | 0.0000 | 649.0 | 33.15% | 20435.1 | 21.00% |
| 22 | RACTACAACTCCCAGVAKGC | Ronin(THAP)/ES-Thap11-ChIP-Seq(GSE51522)/Homer | 1e-33 | -7.691e+01 | 0.0000 | 62.0 | 3.17% | 395.8 | 0.41% |
| 23 | VDGGGYGGGGCY | KLF1(Zf)/HUDEP2-KLF1-CutnRun(GSE136251)/Homer | 1e-32 | -7.535e+01 | 0.0000 | 564.0 | 28.80% | 17253.0 | 17.73% |
| 24 | RGGGGGCGGGGC | Klf15(Zf)/Liver-Klf15-ChIP-Seq(GSE166083)/Homer | 1e-31 | -7.228e+01 | 0.0000 | 620.0 | 31.66% | 19804.6 | 20.35% |
| 25 | GCCACRCCCACY | Klf9(Zf)/GBM-Klf9-ChIP-Seq(GSE62211)/Homer | 1e-31 | -7.189e+01 | 0.0000 | 293.0 | 14.96% | 7011.8 | 7.20% |
| 26 | RACTACAATTCCCAGAAKGC | GFY-Staf(?,Zf)/Promoter/Homer | 1e-28 | -6.485e+01 | 0.0000 | 66.0 | 3.37% | 567.3 | 0.58% |
| 27 | ACAGGAAGTG | ERG(ETS)/VCaP-ERG-ChIP-Seq(GSE14097)/Homer | 1e-26 | -6.122e+01 | 0.0000 | 502.0 | 25.64% | 15649.0 | 16.08% |
| 28 | WNTGCTGASTCAGCANWTTY | MafB(bZIP)/BMM-Mafb-ChIP-Seq(GSE75722)/Homer | 1e-24 | -5.631e+01 | 0.0000 | 149.0 | 7.61% | 2835.8 | 2.91% |
| 29 | NRYTTCCGGH | Fli1(ETS)/CD8-FLI-ChIP-Seq(GSE20898)/Homer | 1e-24 | -5.557e+01 | 0.0000 | 402.0 | 20.53% | 11975.7 | 12.31% |
| 30 | ACAGGAAGTG | ETS1(ETS)/Jurkat-ETS1-ChIP-Seq(GSE17954)/Homer | 1e-24 | -5.556e+01 | 0.0000 | 365.0 | 18.64% | 10518.1 | 10.81% |
| 31 | AACCGGAAGT | ETV1(ETS)/GIST48-ETV1-ChIP-Seq(GSE22441)/Homer | 1e-23 | -5.496e+01 | 0.0000 | 452.0 | 23.08% | 14034.1 | 14.42% |
| 32 | NNAYTTCCTGHN | Etv2(ETS)/ES-ER71-ChIP-Seq(GSE59402)/Homer | 1e-23 | -5.464e+01 | 0.0000 | 318.0 | 16.24% | 8765.6 | 9.01% |
| 33 | ACCGGAAGTG | ETV4(ETS)/HepG2-ETV4-ChIP-Seq(ENCODE)/Homer | 1e-23 | -5.306e+01 | 0.0000 | 423.0 | 21.60% | 12986.0 | 13.34% |
| 34 | YGGCCCCGCCCC | Sp2(Zf)/HEK293-Sp2.eGFP-ChIP-Seq(Encode)/Homer | 1e-21 | -4.918e+01 | 0.0000 | 759.0 | 38.76% | 27884.6 | 28.65% |
| 35 | MKGGGYGTGGCC | KLF6(Zf)/PDAC-KLF6-ChIP-Seq(GSE64557)/Homer | 1e-21 | -4.865e+01 | 0.0000 | 558.0 | 28.50% | 18988.0 | 19.51% |
| 36 | ACTACAATTCCC | GFY(?)/Promoter/Homer | 1e-19 | -4.578e+01 | 0.0000 | 60.0 | 3.06% | 678.7 | 0.70% |
| 37 | TGCTGACTCA | MafA(bZIP)/Islet-MafA-ChIP-Seq(GSE30298)/Homer | 1e-19 | -4.494e+01 | 0.0000 | 240.0 | 12.26% | 6380.3 | 6.56% |
| 38 | VACAGGAAAT | EWS:FLI1-fusion(ETS)/SK_N_MC-EWS:FLI1-ChIP-Seq(SRA014231)/Homer | 1e-19 | -4.488e+01 | 0.0000 | 209.0 | 10.67% | 5260.8 | 5.41% |
| 39 | RACCGGAAGT | GABPA(ETS)/Jurkat-GABPa-ChIP-Seq(GSE17954)/Homer | 1e-19 | -4.412e+01 | 0.0000 | 316.0 | 16.14% | 9321.0 | 9.58% |
| 40 | ACTTCCKGKT | Elf4(ETS)/BMDM-Elf4-ChIP-Seq(GSE88699)/Homer | 1e-18 | -4.344e+01 | 0.0000 | 332.0 | 16.96% | 9997.8 | 10.27% |
| 41 | DGGGYGKGGC | KLF5(Zf)/LoVo-KLF5-ChIP-Seq(GSE49402)/Homer | 1e-18 | -4.335e+01 | 0.0000 | 637.0 | 32.53% | 22949.6 | 23.58% |
| 42 | ANCAGGAAGT | ELF3(ETS)/PDAC-ELF3-ChIP-Seq(GSE64557)/Homer | 1e-17 | -4.028e+01 | 0.0000 | 218.0 | 11.13% | 5819.4 | 5.98% |
| 43 | AVCAGGAAGT | EHF(ETS)/LoVo-EHF-ChIP-Seq(GSE49402)/Homer | 1e-17 | -3.953e+01 | 0.0000 | 364.0 | 18.59% | 11569.6 | 11.89% |
| 44 | HACTTCCGGY | Elk1(ETS)/Hela-Elk1-ChIP-Seq(GSE31477)/Homer | 1e-16 | -3.848e+01 | 0.0000 | 245.0 | 12.51% | 6932.1 | 7.12% |
| 45 | ATTTCCTGTN | EWS:ERG-fusion(ETS)/CADO_ES1-EWS:ERG-ChIP-Seq(SRA014231)/Homer | 1e-15 | -3.646e+01 | 0.0000 | 198.0 | 10.11% | 5290.4 | 5.44% |
| 46 | MACAGGAAGT | Ets1-distal(ETS)/CD4+-PolII-ChIP-Seq(Barski_et_al.)/Homer | 1e-15 | -3.454e+01 | 0.0000 | 108.0 | 5.52% | 2264.7 | 2.33% |
| 47 | GCCACACCCA | Klf4(Zf)/mES-Klf4-ChIP-Seq(GSE11431)/Homer | 1e-14 | -3.385e+01 | 0.0000 | 202.0 | 10.32% | 5584.7 | 5.74% |
| 48 | RGKGGGCGKGGC | KLF14(Zf)/HEK293-KLF14.GFP-ChIP-Seq(GSE58341)/Homer | 1e-14 | -3.332e+01 | 0.0000 | 812.0 | 41.47% | 32133.5 | 33.02% |
| 49 | CTGCGCATGCGC | NRF1(NRF)/MCF7-NRF1-ChIP-Seq(Unpublished)/Homer | 1e-13 | -3.073e+01 | 0.0000 | 130.0 | 6.64% | 3147.1 | 3.23% |
| 50 | GGGGGTGTGTCC | KLF10(Zf)/HEK293-KLF10.GFP-ChIP-Seq(GSE58341)/Homer | 1e-13 | -3.019e+01 | 0.0000 | 231.0 | 11.80% | 6927.5 | 7.12% |
| 51 | AVCCGGAAGT | ELF1(ETS)/Jurkat-ELF1-ChIP-Seq(SRA014231)/Homer | 1e-12 | -2.945e+01 | 0.0000 | 220.0 | 11.24% | 6548.6 | 6.73% |
| 52 | STGCGCATGCGC | NRF(NRF)/Promoter/Homer | 1e-12 | -2.878e+01 | 0.0000 | 121.0 | 6.18% | 2924.5 | 3.00% |
| 53 | CCWGGAATGY | TEAD4(TEA)/Tropoblast-Tead4-ChIP-Seq(GSE37350)/Homer | 1e-11 | -2.679e+01 | 0.0000 | 222.0 | 11.34% | 6815.2 | 7.00% |
| 54 | ASWTCCTGBT | SPDEF(ETS)/VCaP-SPDEF-ChIP-Seq(SRA014231)/Homer | 1e-11 | -2.583e+01 | 0.0000 | 272.0 | 13.89% | 8909.1 | 9.15% |
| 55 | CCCCTCCCCCAC | ZNF148(Zf)/MDAMB231-ZNF148-ChIP-Seq(GSE147020)/Homer | 1e-11 | -2.565e+01 | 0.0000 | 347.0 | 17.72% | 12067.9 | 12.40% |
| 56 | VTTACGTAAYNNNNN | NFIL3(bZIP)/HepG2-NFIL3-ChIP-Seq(Encode)/Homer | 1e-10 | -2.532e+01 | 0.0000 | 136.0 | 6.95% | 3628.5 | 3.73% |
| 57 | CTTCCGGGAA | Stat3(Stat)/mES-Stat3-ChIP-Seq(GSE11431)/Homer | 1e-10 | -2.522e+01 | 0.0000 | 154.0 | 7.87% | 4298.3 | 4.42% |
| 58 | ACVAGGAAGT | ELF5(ETS)/T47D-ELF5-ChIP-Seq(GSE30407)/Homer | 1e-10 | -2.500e+01 | 0.0000 | 205.0 | 10.47% | 6275.6 | 6.45% |
| 59 | NRYTTCCGGY | Elk4(ETS)/Hela-Elk4-ChIP-Seq(GSE31477)/Homer | 1e-10 | -2.470e+01 | 0.0000 | 216.0 | 11.03% | 6733.6 | 6.92% |
| 60 | AACCGGAAGT | ETS(ETS)/Promoter/Homer | 1e-9 | -2.253e+01 | 0.0000 | 136.0 | 6.95% | 3787.2 | 3.89% |
| 61 | GGGGGGGG | Maz(Zf)/HepG2-Maz-ChIP-Seq(GSE31477)/Homer | 1e-9 | -2.214e+01 | 0.0000 | 657.0 | 33.55% | 26413.6 | 27.14% |
| 62 | YCWGGAATGY | TEAD(TEA)/Fibroblast-PU.1-ChIP-Seq(Unpublished)/Homer | 1e-9 | -2.196e+01 | 0.0000 | 160.0 | 8.17% | 4734.8 | 4.86% |
| 63 | DGTAAACA | Foxo3(Forkhead)/U2OS-Foxo3-ChIP-Seq(E-MTAB-2701)/Homer | 1e-9 | -2.143e+01 | 0.0000 | 134.0 | 6.84% | 3779.6 | 3.88% |
| 64 | TRCATTCCAG | TEAD3(TEA)/HepG2-TEAD3-ChIP-Seq(Encode)/Homer | 1e-9 | -2.121e+01 | 0.0000 | 247.0 | 12.61% | 8289.9 | 8.52% |
| 65 | CYRCATTCCA | TEAD1(TEAD)/HepG2-TEAD1-ChIP-Seq(Encode)/Homer | 1e-9 | -2.093e+01 | 0.0000 | 224.0 | 11.44% | 7366.7 | 7.57% |
| 66 | CCWTTGTYYB | Sox10(HMG)/SciaticNerve-Sox3-ChIP-Seq(GSE35132)/Homer | 1e-8 | -2.015e+01 | 0.0000 | 300.0 | 15.32% | 10629.8 | 10.92% |
| 67 | CTGTTTAC | Foxo1(Forkhead)/RAW-Foxo1-ChIP-Seq(Fan_et_al.)/Homer | 1e-8 | -1.869e+01 | 0.0000 | 371.0 | 18.95% | 13889.4 | 14.27% |
| 68 | GWAAYHTGAKMC | Six2(Homeobox)/NephronProgenitor-Six2-ChIP-Seq(GSE39837)/Homer | 1e-7 | -1.790e+01 | 0.0000 | 190.0 | 9.70% | 6242.6 | 6.41% |
| 69 | SAAACCACAG | RUNX(Runt)/HPC7-Runx1-ChIP-Seq(GSE22178)/Homer | 1e-7 | -1.790e+01 | 0.0000 | 152.0 | 7.76% | 4715.1 | 4.84% |
| 70 | SVYTTCCNGGAARB | Stat3+il21(Stat)/CD4-Stat3-ChIP-Seq(GSE19198)/Homer | 1e-7 | -1.773e+01 | 0.0000 | 171.0 | 8.73% | 5486.1 | 5.64% |
| 71 | AAACCACARM | RUNX1(Runt)/Jurkat-RUNX1-ChIP-Seq(GSE29180)/Homer | 1e-7 | -1.653e+01 | 0.0000 | 200.0 | 10.21% | 6781.3 | 6.97% |
| 72 | CACAGCAGGGGG | Unknown-ESC-element(?)/mES-Nanog-ChIP-Seq(GSE11724)/Homer | 1e-6 | -1.546e+01 | 0.0000 | 201.0 | 10.27% | 6928.1 | 7.12% |
| 73 | WWATRTAAACAN | Foxf1(Forkhead)/Lung-Foxf1-ChIP-Seq(GSE77951)/Homer | 1e-6 | -1.544e+01 | 0.0000 | 150.0 | 7.66% | 4833.4 | 4.97% |
| 74 | CCCCTCCCCCAC | Zfp281(Zf)/ES-Zfp281-ChIP-Seq(GSE81042)/Homer | 1e-6 | -1.502e+01 | 0.0000 | 116.0 | 5.92% | 3518.7 | 3.62% |
| 75 | CCWGGAATGY | TEAD2(TEA)/Py2T-Tead2-ChIP-Seq(GSE55709)/Homer | 1e-6 | -1.418e+01 | 0.0000 | 128.0 | 6.54% | 4056.3 | 4.17% |
| 76 | MTGATGCAAT | Atf4(bZIP)/MEF-Atf4-ChIP-Seq(GSE35681)/Homer | 1e-6 | -1.401e+01 | 0.0000 | 66.0 | 3.37% | 1708.7 | 1.76% |
| 77 | CSGTGACGTCAC | CRE(bZIP)/Promoter/Homer | 1e-6 | -1.394e+01 | 0.0000 | 77.0 | 3.93% | 2110.3 | 2.17% |
| 78 | TGWAAYCTGABACCB | Six4(Homeobox)/MCF7-SIX4-ChIP-Seq(Encode)/Homer | 1e-5 | -1.377e+01 | 0.0000 | 21.0 | 1.07% | 293.7 | 0.30% |
| 79 | YCATYAATCA | Pdx1(Homeobox)/Islet-Pdx1-ChIP-Seq(SRA008281)/Homer | 1e-5 | -1.266e+01 | 0.0000 | 157.0 | 8.02% | 5375.6 | 5.52% |
| 80 | AGAGGAAGTG | PU.1(ETS)/ThioMac-PU.1-ChIP-Seq(GSE21512)/Homer | 1e-5 | -1.254e+01 | 0.0000 | 128.0 | 6.54% | 4195.8 | 4.31% |
| 81 | WWTRTAAACAVG | FoxL2(Forkhead)/Ovary-FoxL2-ChIP-Seq(GSE60858)/Homer | 1e-5 | -1.235e+01 | 0.0000 | 135.0 | 6.89% | 4497.0 | 4.62% |
| 82 | GKVTCADRTTWC | Six1(Homeobox)/Myoblast-Six1-ChIP-Chip(GSE20150)/Homer | 1e-5 | -1.210e+01 | 0.0000 | 55.0 | 2.81% | 1415.2 | 1.45% |
| 83 | RTGATTKATRGN | PBX2(Homeobox)/K562-PBX2-ChIP-Seq(Encode)/Homer | 1e-5 | -1.154e+01 | 0.0001 | 132.0 | 6.74% | 4450.2 | 4.57% |
| 84 | CCWTTGTY | Sox3(HMG)/NPC-Sox3-ChIP-Seq(GSE33059)/Homer | 1e-5 | -1.153e+01 | 0.0001 | 287.0 | 14.66% | 11145.0 | 11.45% |
| 85 | ATTGCATCAT | Chop(bZIP)/MEF-Chop-ChIP-Seq(GSE35681)/Homer | 1e-4 | -1.126e+01 | 0.0001 | 53.0 | 2.71% | 1388.0 | 1.43% |
| 86 | YCTTTGTTCC | Sox4(HMG)/proB-Sox4-ChIP-Seq(GSE50066)/Homer | 1e-4 | -1.122e+01 | 0.0001 | 156.0 | 7.97% | 5483.6 | 5.63% |
| 87 | NYYTGTTTACHN | FOXP1(Forkhead)/H9-FOXP1-ChIP-Seq(GSE31006)/Homer | 1e-4 | -1.084e+01 | 0.0001 | 72.0 | 3.68% | 2116.2 | 2.17% |
| 88 | TGGGGAAGGGCM | ZNF467(Zf)/HEK293-ZNF467.GFP-ChIP-Seq(GSE58341)/Homer | 1e-4 | -1.081e+01 | 0.0001 | 339.0 | 17.31% | 13603.3 | 13.98% |
| 89 | NVWTGTTTAC | FOXK1(Forkhead)/HEK293-FOXK1-ChIP-Seq(GSE51673)/Homer | 1e-4 | -1.080e+01 | 0.0001 | 160.0 | 8.17% | 5699.7 | 5.86% |
| 90 | ATTTCCCAGVAKSCY | ZNF143\|STAF(Zf)/CUTLL-ZNF143-ChIP-Seq(GSE29600)/Homer | 1e-4 | -1.070e+01 | 0.0001 | 111.0 | 5.67% | 3669.5 | 3.77% |
| 91 | NWAACCACADNN | RUNX2(Runt)/PCa-RUNX2-ChIP-Seq(GSE33889)/Homer | 1e-4 | -1.068e+01 | 0.0001 | 145.0 | 7.41% | 5078.6 | 5.22% |
| 92 | VVATGACGTCAT | CREB5(bZIP)/LNCaP-CREB5.V5-ChIP-Seq(GSE137775)/Homer | 1e-4 | -1.049e+01 | 0.0001 | 84.0 | 4.29% | 2606.9 | 2.68% |
| 93 | NNNCTTTCCAGGAAA | Bcl6(Zf)/Liver-Bcl6-ChIP-Seq(GSE31578)/Homer | 1e-4 | -1.049e+01 | 0.0001 | 264.0 | 13.48% | 10278.0 | 10.56% |
| 94 | NYTTCCWGGAAR | STAT4(Stat)/CD4-Stat4-ChIP-Seq(GSE22104)/Homer | 1e-4 | -1.027e+01 | 0.0002 | 176.0 | 8.99% | 6445.6 | 6.62% |
| 95 | BSNTGTTTACWYWGN | Foxa3(Forkhead)/Liver-Foxa3-ChIP-Seq(GSE77670)/Homer | 1e-4 | -1.023e+01 | 0.0002 | 56.0 | 2.86% | 1554.8 | 1.60% |
| 96 | TTCCKNAGAA | STAT6(Stat)/Macrophage-Stat6-ChIP-Seq(GSE38377)/Homer | 1e-4 | -9.798e+00 | 0.0003 | 103.0 | 5.26% | 3426.1 | 3.52% |
| 97 | CCTGCTGAGH | Zic(Zf)/Cerebellum-ZIC1.2-ChIP-Seq(GSE60731)/Homer | 1e-4 | -9.552e+00 | 0.0003 | 244.0 | 12.46% | 9530.9 | 9.79% |
| 98 | RTTATGYAAB | HLF(bZIP)/HSC-HLF.Flag-ChIP-Seq(GSE69817)/Homer | 1e-3 | -9.097e+00 | 0.0005 | 129.0 | 6.59% | 4578.2 | 4.70% |
| 99 | AGGVNCCTTTGT | Sox9(HMG)/Limb-SOX9-ChIP-Seq(GSE73225)/Homer | 1e-3 | -9.033e+00 | 0.0006 | 153.0 | 7.81% | 5609.6 | 5.76% |
| 100 | TGTTTAYTTAGC | FoxD3(forkhead)/ZebrafishEmbryo-Foxd3.biotin-ChIP-seq(GSE106676)/Homer | 1e-3 | -9.011e+00 | 0.0006 | 135.0 | 6.89% | 4842.2 | 4.98% |
| 101 | BCCWTTGTBYKV | Sox21(HMG)/ESC-SOX21-ChIP-Seq(GSE110505)/Homer | 1e-3 | -8.996e+00 | 0.0006 | 279.0 | 14.25% | 11196.0 | 11.50% |
| 102 | CCATTGTTYB | Sox17(HMG)/Endoderm-Sox17-ChIP-Seq(GSE61475)/Homer | 1e-3 | -8.916e+00 | 0.0006 | 123.0 | 6.28% | 4345.2 | 4.46% |
| 103 | GCTGTGGTTW | RUNX-AML(Runt)/CD4+-PolII-ChIP-Seq(Barski_et_al.)/Homer | 1e-3 | -8.904e+00 | 0.0006 | 132.0 | 6.74% | 4727.8 | 4.86% |
| 104 | ATTTGCATAW | Oct4(POU,Homeobox)/mES-Oct4-ChIP-Seq(GSE11431)/Homer | 1e-3 | -8.881e+00 | 0.0006 | 79.0 | 4.03% | 2537.6 | 2.61% |
| 105 | SARTGGAAAAWRTGAGTCAB | NFAT:AP1(RHD,bZIP)/Jurkat-NFATC1-ChIP-Seq(Jolma_et_al.)/Homer | 1e-3 | -8.844e+00 | 0.0006 | 39.0 | 1.99% | 1013.2 | 1.04% |
| 106 | CAAGATGGCGGC | YY1(Zf)/Promoter/Homer | 1e-3 | -8.517e+00 | 0.0009 | 33.0 | 1.69% | 818.2 | 0.84% |
| 107 | GGCCYCCTGCTGDGH | Zic3(Zf)/mES-Zic3-ChIP-Seq(GSE37889)/Homer | 1e-3 | -8.094e+00 | 0.0013 | 233.0 | 11.90% | 9274.7 | 9.53% |
| 108 | CCATTGTTNY | Sox6(HMG)/Myotubes-Sox6-ChIP-Seq(GSE32627)/Homer | 1e-3 | -8.036e+00 | 0.0014 | 246.0 | 12.56% | 9870.8 | 10.14% |
| 109 | ATGACGTCATCN | JunD(bZIP)/K562-JunD-ChIP-Seq/Homer | 1e-3 | -7.804e+00 | 0.0018 | 28.0 | 1.43% | 678.9 | 0.70% |
| 110 | NATTTCCNGGAAAT | STAT1(Stat)/HelaS3-STAT1-ChIP-Seq(GSE12782)/Homer | 1e-3 | -7.722e+00 | 0.0019 | 59.0 | 3.01% | 1837.1 | 1.89% |
| 111 | ATGAATATTC | Brn2(POU,Homeobox)/NPC-Brn2-ChIP-Seq(GSE35496)/Homer | 1e-3 | -7.617e+00 | 0.0021 | 22.0 | 1.12% | 485.6 | 0.50% |
| 112 | NRRTGACGTCAT | Atf2(bZIP)/3T3L1-Atf2-ChIP-Seq(GSE56872)/Homer | 1e-3 | -7.575e+00 | 0.0022 | 81.0 | 4.14% | 2735.6 | 2.81% |
| 113 | ATGACGTCATCY | c-Jun-CRE(bZIP)/K562-cJun-ChIP-Seq(GSE31477)/Homer | 1e-3 | -7.505e+00 | 0.0023 | 76.0 | 3.88% | 2537.3 | 2.61% |
| 114 | MGGAAGTGAAAC | PU.1-IRF(ETS:IRF)/Bcell-PU.1-ChIP-Seq(GSE21512)/Homer | 1e-3 | -7.354e+00 | 0.0027 | 281.0 | 14.35% | 11591.6 | 11.91% |
| 115 | GGGGCTYGKCTGGGA | Zfp809(Zf)/ES-Zfp809-ChIP-Seq(GSE70799)/Homer | 1e-3 | -7.349e+00 | 0.0027 | 111.0 | 5.67% | 4016.1 | 4.13% |
| 116 | SCHTGTTTACAT | FOXK2(Forkhead)/U2OS-FOXK2-ChIP-Seq(E-MTAB-2204)/Homer | 1e-3 | -7.183e+00 | 0.0031 | 102.0 | 5.21% | 3653.2 | 3.75% |
| 117 | MCTCCCMCRCAB | WT1(Zf)/Kidney-WT1-ChIP-Seq(GSE90016)/Homer | 1e-3 | -7.149e+00 | 0.0032 | 245.0 | 12.51% | 9986.3 | 10.26% |
| 118 | ATGCATWATGCATRW | OCT:OCT-short(POU,Homeobox)/NPC-OCT6-ChIP-Seq(GSE43916)/Homer | 1e-3 | -7.004e+00 | 0.0036 | 106.0 | 5.41% | 3844.6 | 3.95% |
| 119 | CCATTGTTCB | SOX1(HMG)/NPC-SOX1-ChIP-Seq(GSE138215)/Homer | 1e-2 | -6.891e+00 | 0.0040 | 341.0 | 17.42% | 14463.9 | 14.86% |
| 120 | TGTTKCCTAGCAACM | Rfx6(HTH)/Min6b1-Rfx6.HA-ChIP-Seq(GSE62844)/Homer | 1e-2 | -6.868e+00 | 0.0041 | 253.0 | 12.92% | 10405.9 | 10.69% |
| 121 | WAAGTAAACA | FOXA1(Forkhead)/MCF7-FOXA1-ChIP-Seq(GSE26831)/Homer | 1e-2 | -6.850e+00 | 0.0041 | 142.0 | 7.25% | 5423.9 | 5.57% |
| 122 | RAACAATGGN | Sox15(HMG)/CPA-Sox15-ChIP-Seq(GSE62909)/Homer | 1e-2 | -6.668e+00 | 0.0049 | 181.0 | 9.24% | 7181.4 | 7.38% |
| 123 | NGCGTGGGCGGR | Egr2(Zf)/Thymocytes-Egr2-ChIP-Seq(GSE34254)/Homer | 1e-2 | -6.411e+00 | 0.0063 | 95.0 | 4.85% | 3446.9 | 3.54% |
| 124 | BCCATTGTTC | Sox2(HMG)/mES-Sox2-ChIP-Seq(GSE11431)/Homer | 1e-2 | -6.409e+00 | 0.0063 | 141.0 | 7.20% | 5444.7 | 5.59% |
| 125 | TTTTATKRGN | Hoxc13(Homeobox)/EB-Hoxc13.HA-ChIP-Seq(GSE142377)/Homer | 1e-2 | -6.368e+00 | 0.0065 | 200.0 | 10.21% | 8089.7 | 8.31% |
| 126 | WAAGTAAACA | FOXA1(Forkhead)/LNCAP-FOXA1-ChIP-Seq(GSE27824)/Homer | 1e-2 | -6.300e+00 | 0.0069 | 168.0 | 8.58% | 6661.2 | 6.84% |
| 127 | ATGCATAATTCA | Pit1+1bp(Homeobox)/GCrat-Pit1-ChIP-Seq(GSE58009)/Homer | 1e-2 | -6.193e+00 | 0.0076 | 52.0 | 2.66% | 1682.4 | 1.73% |
| 128 | DRTGTTGCAA | CEBP:AP1(bZIP)/ThioMac-CEBPb-ChIP-Seq(GSE21512)/Homer | 1e-2 | -6.085e+00 | 0.0084 | 120.0 | 6.13% | 4569.8 | 4.70% |
| 129 | ATGAATWATTCATGA | OCT:OCT(POU,Homeobox,IR1)/NPC-Brn2-ChIP-Seq(GSE35496)/Homer | 1e-2 | -5.818e+00 | 0.0109 | 7.0 | 0.36% | 92.8 | 0.10% |
| 130 | TGCAGTTCCMVNWRTGGCCA | CTCF-SatelliteElement(Zf?)/CD4+-CTCF-ChIP-Seq(Barski_et_al.)/Homer | 1e-2 | -5.724e+00 | 0.0119 | 8.0 | 0.41% | 119.1 | 0.12% |
| 131 | GTCACGTGACYV | TFE3(bHLH)/MEF-TFE3-ChIP-Seq(GSE75757)/Homer | 1e-2 | -5.655e+00 | 0.0126 | 23.0 | 1.17% | 605.4 | 0.62% |
| 132 | CGGTTGCCATGGCAAC | RFX(HTH)/K562-RFX3-ChIP-Seq(SRA012198)/Homer | 1e-2 | -5.607e+00 | 0.0131 | 25.0 | 1.28% | 681.0 | 0.70% |
| 133 | NNNNNBAGATAWYATCTVHN | GATA(Zf),IR3/iTreg-Gata3-ChIP-Seq(GSE20898)/Homer | 1e-2 | -5.587e+00 | 0.0133 | 32.0 | 1.63% | 943.4 | 0.97% |
| 134 | AVCAGCTG | SCL(bHLH)/HPC7-Scl-ChIP-Seq(GSE13511)/Homer | 1e-2 | -5.494e+00 | 0.0145 | 975.0 | 49.80% | 45535.1 | 46.79% |
| 135 | NWGGGTGTGGCY | EKLF(Zf)/Erythrocyte-Klf1-ChIP-Seq(GSE20478)/Homer | 1e-2 | -5.438e+00 | 0.0152 | 62.0 | 3.17% | 2162.4 | 2.22% |
| 136 | NNTGTGGATTSS | Foxh1(Forkhead)/hESC-FOXH1-ChIP-Seq(GSE29422)/Homer | 1e-2 | -5.405e+00 | 0.0156 | 96.0 | 4.90% | 3617.5 | 3.72% |
| 137 | CHCAGCRGGRGG | Zic2(Zf)/ESC-Zic2-ChIP-Seq(SRP197560)/Homer | 1e-2 | -5.330e+00 | 0.0167 | 185.0 | 9.45% | 7605.9 | 7.82% |
| 138 | GTTGCCATGGCAACM | Rfx2(HTH)/LoVo-RFX2-ChIP-Seq(GSE49402)/Homer | 1e-2 | -5.242e+00 | 0.0181 | 26.0 | 1.33% | 738.2 | 0.76% |
| 139 | RTTTCTNAGAAA | STAT5(Stat)/mCD4+-Stat5-ChIP-Seq(GSE12346)/Homer | 1e-2 | -5.232e+00 | 0.0181 | 60.0 | 3.06% | 2100.9 | 2.16% |
| 140 | GSCTWTAAAAGG | TATA-Box(TBP)/Promoter/Homer | 1e-2 | -5.068e+00 | 0.0212 | 192.0 | 9.81% | 7978.6 | 8.20% |
| 141 | NGYCATAAAWCH | CDX4(Homeobox)/ZebrafishEmbryos-Cdx4.Myc-ChIP-Seq(GSE48254)/Homer | 1e-2 | -5.067e+00 | 0.0212 | 120.0 | 6.13% | 4723.2 | 4.85% |
| 142 | GATGACGTCA | Atf1(bZIP)/K562-ATF1-ChIP-Seq(GSE31477)/Homer | 1e-2 | -5.035e+00 | 0.0216 | 145.0 | 7.41% | 5848.2 | 6.01% |
| 143 | GGCVGTTR | MYB(HTH)/ERMYB-Myb-ChIPSeq(GSE22095)/Homer | 1e-2 | -4.993e+00 | 0.0224 | 302.0 | 15.42% | 13103.2 | 13.46% |
| 144 | NRSCWCGYGC | Zfp541(Zf)/Sperm-Zfp541-ChIP-Seq(GSE163916)/Homer | 1e-2 | -4.930e+00 | 0.0237 | 394.0 | 20.12% | 17469.3 | 17.95% |
| 145 | SSGGTCACGTGA | E-box(bHLH)/Promoter/Homer | 1e-2 | -4.919e+00 | 0.0238 | 27.0 | 1.38% | 796.6 | 0.82% |
| 146^a^ | GGCCATAAATCA | Hoxc9(Homeobox)/Ainv15-Hoxc9-ChIP-Seq(GSE21812)/Homer | 1e-2 | -4.821e+00 | 0.0260 | 70.0 | 3.58% | 2570.7 | 2.64% |
| 147 | GTCACGTGGT | Usf2(bHLH)/C2C12-Usf2-ChIP-Seq(GSE36030)/Homer | 1e-2 | -4.781e+00 | 0.0269 | 75.0 | 3.83% | 2790.4 | 2.87% |
| 148 | AGGCCTRG | ZFX(Zf)/mES-Zfx-ChIP-Seq(GSE11431)/Homer | 1e-2 | -4.752e+00 | 0.0275 | 402.0 | 20.53% | 17905.0 | 18.40% |
| 149 | RTGATTTATGGY | Hoxd9(Homeobox)/EB-Hoxd9.HA-ChIP-Seq(GSE142377)/Homer | 1e-2 | -4.748e+00 | 0.0275 | 50.0 | 2.55% | 1733.4 | 1.78% |
| 150 | ABTTCYYRRGAA | STAT6(Stat)/CD4-Stat6-ChIP-Seq(GSE22104)/Homer | 1e-2 | -4.745e+00 | 0.0275 | 87.0 | 4.44% | 3316.2 | 3.41% |
| 151 | NGRTGACGTCAY | Atf7(bZIP)/3T3L1-Atf7-ChIP-Seq(GSE56872)/Homer | 1e-2 | -4.728e+00 | 0.0276 | 110.0 | 5.62% | 4333.8 | 4.45% |
